# Micropatterned Organoids Enable Modeling of the Earliest Stages of Human Cardiac Vascularization

**DOI:** 10.1101/2022.07.08.499233

**Authors:** Oscar J. Abilez, Huaxiao Yang, Lei Tian, Kitchener D. Wilson, Evan H. Lyall, Mengcheng Shen, Rahulkumar Bhoi, Yan Zhuge, Fangjun Jia, Hung Ta Wo, Gao Zhou, Yuan Guan, Bryan Aldana, Detlef Obal, Gary Peltz, Christopher K. Zarins, Joseph C. Wu

## Abstract

Although model organisms have provided insight into the earliest stages of cardiac vascularization, we know very little about this process in humans. Here we show that spatially micropatterned human pluripotent stem cells (hPSCs) enable *in vitro* modeling of this process, corresponding to the first three weeks of *in vivo* human development. Using four hPSC fluorescent reporter lines, we create cardiac vascular organoids (cVOs) by identifying conditions that simultaneously give rise to spatially organized and branched vascular networks within endocardial, myocardial, and epicardial cells. Using single-cell transcriptomics, we show that the cellular composition of cVOs resembles that of a 6.5 post-conception week (PCW) human heart. We find that NOTCH and BMP pathways are upregulated in cVOs, and their inhibition disrupts vascularization. Finally, using the same vascular-inducing factors to create cVOs, we produce hepatic vascular organoids (hVOs). This suggests there is a conserved developmental program for creating vasculature within different organ systems.

**Graphic Abstract:** 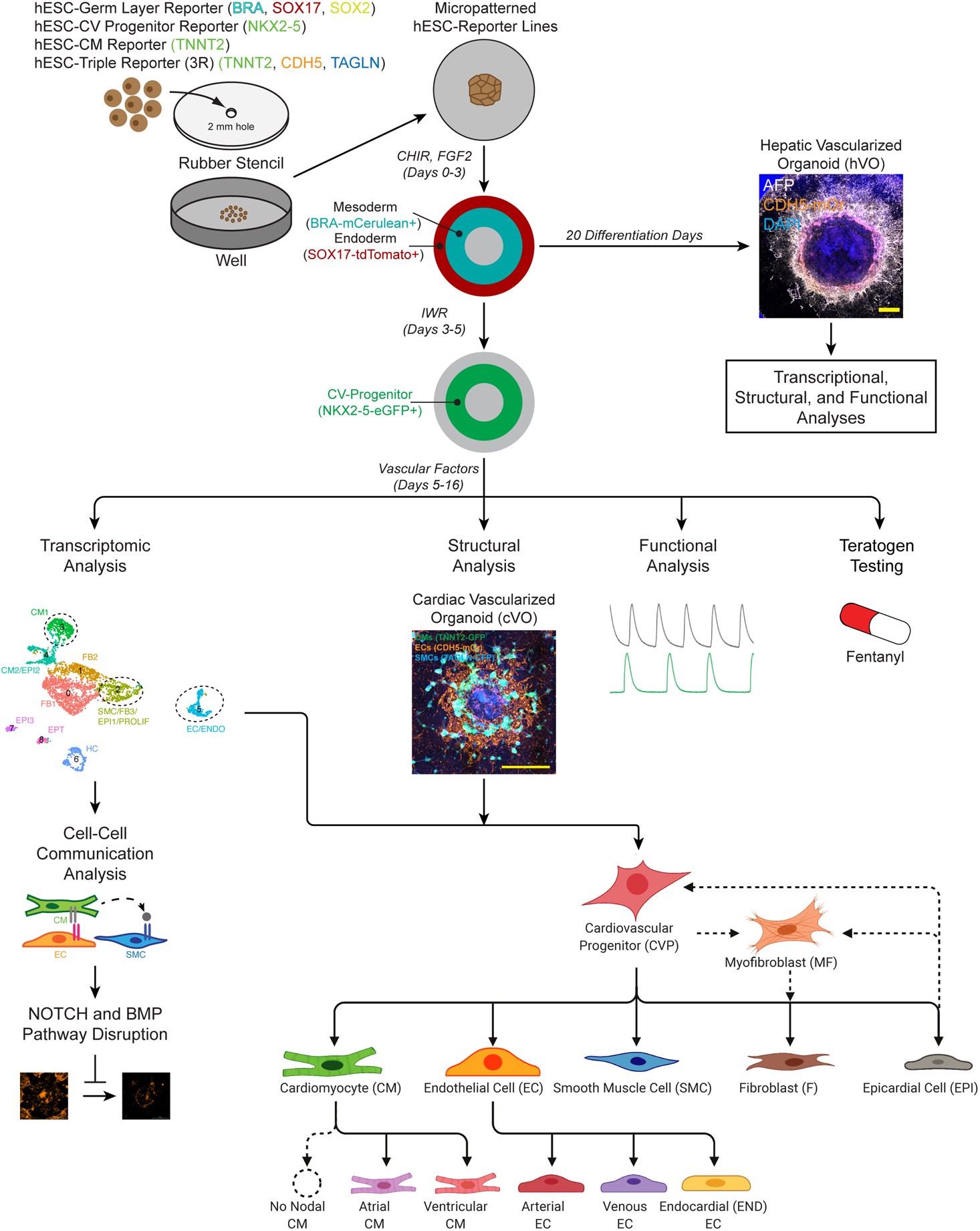

## Introduction

Human pluripotent stem cells (hPSCs), including human embryonic stem cells (hESCs) and human induced pluripotent stem cells (hiPSCs) can differentiate into any cell type in the body^1, 2^. Through specification of mesoderm and endoderm^3, 4^, and then progenitor intermediates^5, 6^, labs around the world can routinely differentiate hPSCs into cardiomyocytes (CMs)^7^, hepatocytes (HCs)^8^ and individual cardiovascular cell types^9^, including endothelial cells (ECs)^10^, smooth muscle cells (SMCs)^11, 12^, fibroblast cells (FBs)^13^, endocardial cells (ENDOs)^14^, and epicardial cells (EPIs)^15^. Furthermore, engineered cardiac^16–18^ and hepatic^19^ tissues with integrated ECs and stromal cells have been created through classic tissue engineering techniques while vascular networks^20, 21^ have been produced using stereolithography and 3D bioprinting. Moreover, labs have used self-organizing techniques to create either cardiac^22–29^, hepatic^30^, or vascular^31^ organoids. The *de novo* vascularization of kidney^32^ and brain^33^ organoids has also been achieved through *in vivo* induction. However, the lack of a prospective and directed *in vitro* strategy to create *de novo* vasculature with robust branching and hierarchical organization within organoids is a major bottleneck in the stem cell field^34–36^. Although model organisms have provided insight into the earliest stages of cardiac vascularization^37, 38^, we know very little about this process in humans, due to ethical restrictions and the technical difficulty of obtaining embryos at such early stages of development^39^.

Here we show that clues from developmental biology^5, 40, 41^ enable *in vitro* modeling of the earliest developmental stages of cardiac vascularization, roughly corresponding to the first three weeks of *in vivo* human development (Carnegie Stages 9 and 10)^42^. Using the four hPSC fluorescent reporter systems hESC-RUES-GLR^43^, hESC-NKX2-5-eGFP^6^, hESC-TNNT2-GFP^44^, and hESC-3R (this paper), along with spatially micropatterned hPSCs^22, 45^, we create cardiac vascularized organoids (cVOs) in a repeatable and scalable fashion. Importantly, the four hPSC reporter systems enable temporal identification of germ layer, progenitor, and cardiovascular cell types *in situ* without disturbing the architecture or orientation of developing cVOs. Addition of a combination of growth factors to micropatterned hPSCs simultaneously generates a spatially organized and branched vascular network within endocardial, myocardial, epicardial, and progenitor cells, along with numerous extracellular matrix (ECM) proteins.

Using single-cell RNA-sequencing (scRNA-seq), we show that the cellular composition of cVOs is similar to that of a 6.5 post-conception week (PCW) human heart (Carnegie Stages 19 and 20)^46^. Furthermore, we use machine learning^47^ to characterize differences in CM and EC formation. We find that NOTCH and BMP pathways are upregulated in cVOs, and inhibition of these pathways disrupts vascularization. Finally, after the generation of liver-specific progenitor pools, we produce hepatic vascularized organoids (hVOs) using the same vascular-inducing factors to create cVOs. This suggests that there is a conserved developmental program for creating vasculature within different organ systems. In summary, our *in vitro* model provides a significant technical advance for addressing questions regarding organ vascularization.

## Results

### Micropatterning of hPSCs results in spatially organized cardiomyocytes

Geometric micropatterning of human pluripotent stem cells (hPSCs) has been shown by us and others to enable repeatable and scalable formation of spatially organized germ layers (endoderm, mesoderm, and ectoderm)^40, 48^, primitive streak^43, 49^, cardiomyocytes^22, 23, 45^, and cardiac organoids^24^. In each well of a multi-well plate, a single plasma-treated rubber stencil with a central hole was used to create a 2 mm circular hPSC micropattern for cardiovascular differentiation (**Fig. 1a**). Phase (**Fig. 1b** and **Extended Data Fig. 1a**) and confocal fluorescence imaging (**Fig. 1c**) shows hPSC micropatterns were pluripotent throughout a confluent colony. However, there were areas with incomplete hESC confluence in the 6 mm colonies (**Fig. 1c**). For a sense of scale, a photograph shows 2 mm hPSC micropatterns (blue) in the centers of four wells of a 48-well dish (**Fig. 1d**). Using an hESC-TNNT2-GFP fluorescent reporter line^44^, we show that 6 mm micropatterns could be differentiated into CMs (**Fig. 1e-f**) and the progression of this differentiation could be tracked *in situ* over days 0-9 (**Supplementary Video 1**).

**Fig. 1.**
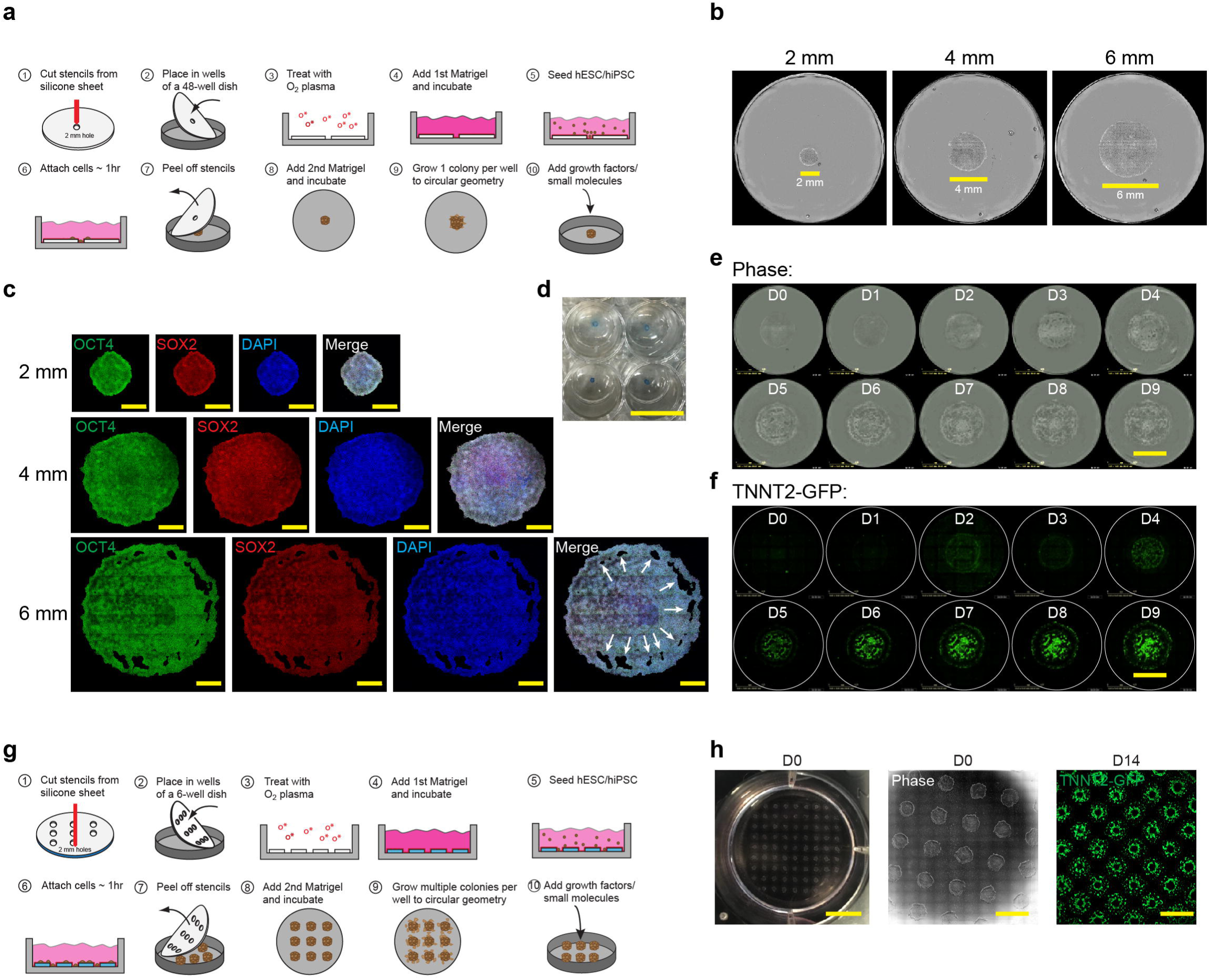
Micropatterning of hPSCs results in spatially organized cardiomyocytes. **a,** A single circular 2 mm hPSC micropattern in each well of a multi-well plate is created from a silicone stencil. Differentiation begins when hPSC micropatterns are 100 % confluent, typically 2-3 days post seeding. **b,** Phase imaging showing 2, 4, and 6 mm undifferentiated hPSC micropatterns. Scale bars, 2, 4, 6 mm, respectively. **c,** Confocal imaging showing 2, 4, and 6 mm hPSC micropatterns are pluripotent by immunostaining for OCT4 (green), SOX2 (red), and nuclear staining with DAPI (blue). White arrows show areas of incomplete hPSC confluence in 6 mm micropatterns. Scale bar, 1 mm. **d,** Photograph showing 2 mm hPSC micropatterns (blue) in the centers of four wells of a 48-well dish. Scale bar, 9 mm. **e,** Phase imaging showing cardiomyocyte (CM) differentiation of a 6 mm hESC-TNNT2-GFP micropattern from Day 0-9 (D0-9). Scale bar, 6 mm. **f,** Fluorescence imaging of (E) showing GFP+ CMs (green). Scale bar, 6 mm. **g,** An array of circular 2 mm hPSC micropatterns in each well of a multi-well plate is created from a silicone stencil. For a 6-well dish, up to 77 hPSC micropatterns can be created per well for a total of 462 micropatterns. **h,** (left) Photograph showing an array of 77 hPSC micropatterns in 1 well of a 6-well dish; (middle) phase imaging showing undifferentiated hPSC micropatterns; (right) fluorescence imaging showing organized and arrayed GFP+ CMs. Scale bars, 10, 4, 4 mm, respectively. See also **Supplementary Video 1**.

To scale up micropattern formation, a single plasma-treated rubber stencil with an array of 2 mm holes was used to create arrayed hPSC micropatterns. For a 6-well dish, up to 77 micropatterns (7 ⨉ 7 array + 7 ⨉ 2 columns + 7 ⨉ 2 rows) could be created per well for a total of 462 micropatterns resulting in 462 GFP^+^ CM colonies (**Fig. 1g-h**). Thus, spatial micropatterning of hPSCs produces spatially organized cardiomyocytes in single and arrayed colony formats. We confirmed that the total fluorescence area of pluripotency markers of our micropatterns was highly correlated with micropattern number (R^2^ = 0.989) and micropattern theoretical total area (R^2^ = 0.99) (**Extended Data Fig. 1b-d**). Of note, validation of our total fluorescence area quantification method was important for our screening experiments described below.

### Micropatterning gives rise to organized germ layer and cardiovascular progenitor formation

To vascularize cardiac organoids, we reasoned that we would only need to induce mesoderm from our hPSC micropatterns to simultaneously produce mesoderm-derived CMs, ECs, and SMCs as previously described^5^. However, to vascularize endoderm-derived hepatic organoids, we would need to simultaneously induce endoderm to produce hepatocytes and mesoderm to produce ECs and SMCs. With this in mind, we used the hESC-RUES-GLR (germ layer reporter) cell line^43^ to visualize germ layer formation under different CHIR-99021 (CHIR) concentrations and unpatterned/micropatterned conditions (**Fig. 2a-b**, and **Extended Data Fig. 2a**).

**Fig. 2.**
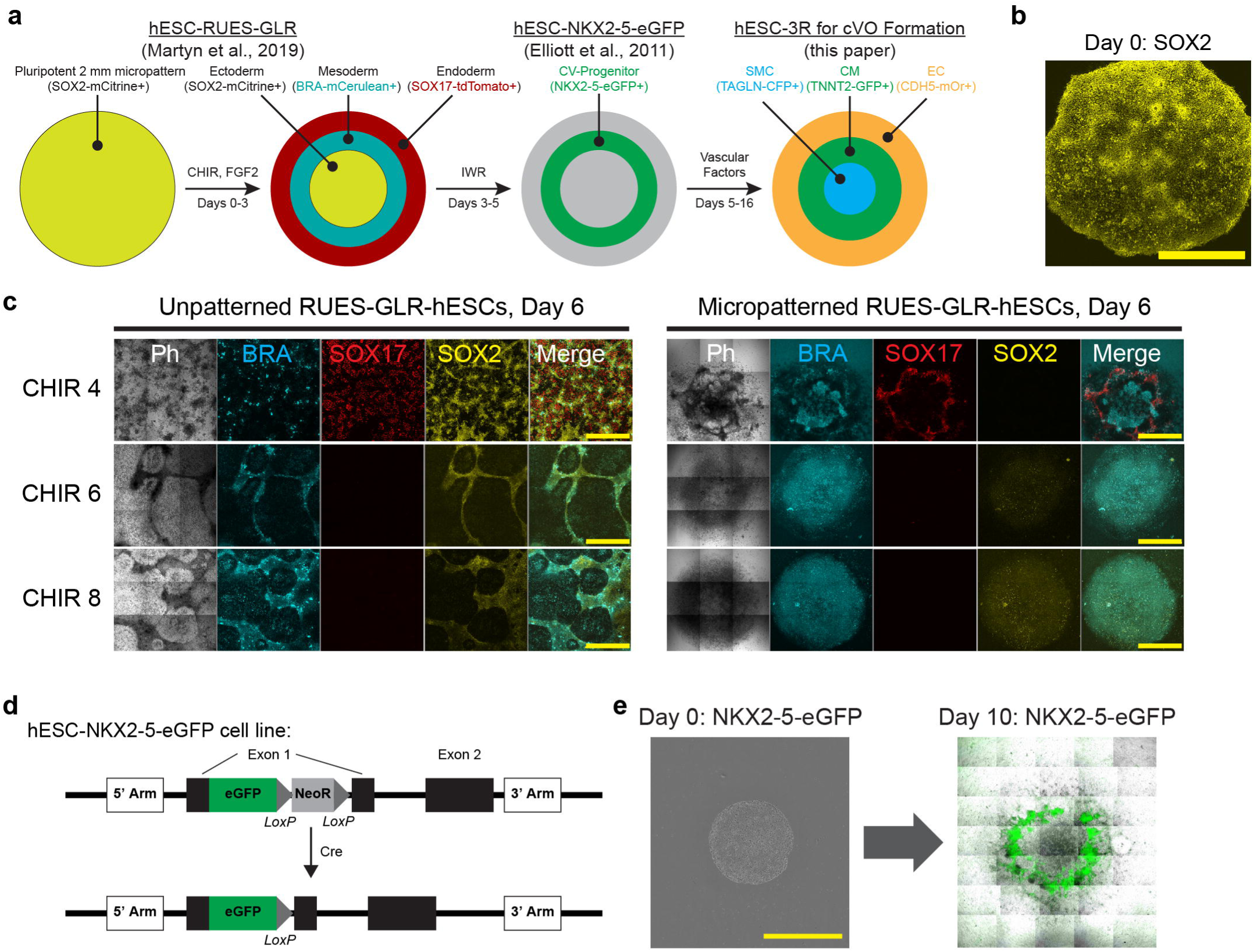
Micropatterning gives rise to organized germ layer and cardiovascular progenitor formation. **a,** The hESC-RUES-GLR (germ layer reporter) cell line ^43^ expresses SOX2, BRA, and SOX17 to identify undifferentiated cells and, upon differentiation, ectoderm, mesoderm, and endoderm arranged in stereotypical concentric rings as shown. The hESC-NKX2-5-eGFP cell line^6^ identifies cardiovascular progenitors in the middle ring arising from mesoderm. The hESC-3R cell line (this paper) identifies SMCs, CMs, and ECs arising from cardiovascular progenitors. **b,** Confocal imaging at day 0 showing SOX2 expression throughout a single undifferentiated micropattern. Scale bar, 1 mm. **c,** Confocal imaging showing unpatterned (left) and micropatterned (right) RUES-GLR hESCs at day 6 with CHIR 4, 6, and 8 µM added at day 0. Unpatterned hESCs give rise to disorganized germ layers (BRA^+^, SOX17^+^, SOX2^+^ expression) and SOX2 is expressed at all CHIR concentrations. In contrast, micropatterned hESCs give rise to organized germ layers with minimal SOX2 expression. For CHIR 4 µM, micropatterned hESCs give rise to a BRA^+^ mesodermal central area surrounded by a SOX17+ endodermal ring, with no SOX2 ectodermal expression. Scale bar, 1 mm. **j,** The hESC-NKX2-5-eGFP cell line expresses eGFP under the cardiovascular progenitor transcription factor NKX2-5 as previously described ^6^. **k,** A single NKX2-5-eGFP micropattern at day 0 (left) differentiated over 10 days showing organized ring formation of eGFP+ cardiovascular progenitors leading to beating cardiomyocytes (right). Scale bar, 2 mm. See also **Supplementary Video 2**.

While unpatterned RUES-GLR hESCs at day 6 gave rise to disorganized germ layer formation at different CHIR concentrations, the micropatterned group gave rise to organized germ layer formation (**Fig. 2c**). With 4 µM of CHIR induction, a central BRA^+^ mesodermal area was surrounded by a SOX17^+^ endodermal ring, and no SOX2 ectodermal expression was detected. In contrast, SOX2 was expressed in the unpatterned group at all CHIR concentrations and in a disorganized fashion. Immunostaining of BRA and SOX17 confirmed the reporter expression seen in the CHIR 4 µM micropattern group (**Extended Data Fig. 2b**).

Arrays of micropatterned RUES-GLR hESCs differentiated under the same conditions as single micropatterns also showed centrally located BRA^+^ expression surrounded by maximum SOX17^+^ expression, which peaked at day 11 for both conditions (**Extended Data Fig. 2c-d**). Thus, single and arrayed micropatterns gave rise to similar BRA and SOX17 temporal and spatial expression, but different from the unpatterned group.

To visualize the effects of micropatterning on cardiovascular progenitor formation, we used the hESC-NKX2-5-eGFP cell line^6^ (**Fig. 2d**). Using our baseline CM differentiation protocol (see Methods), we showed that single NKX2-5-eGFP micropatterns differentiated over 10 days gave rise to organized ring formation of NKX2-5-eGFP+ cardiovascular progenitors, leading to beating cardiomyocytes (**Fig. 2e**; **Supplementary Video 2**). At this juncture, using three fluorescence hPSC reporter lines (hESC-RUES-GLR, hESC-NKX2-5-eGFP, hESC-TNNT2-GFP (from the previous section)), we demonstrated that micropatterning gave rise to organized germ layer, cardiovascular progenitor, and CM formation, thus providing a foundation for screening conditions leading to cVO formation below.

### Creation of a triple reporter line enables identification of differentiation conditions leading to cVO formation

To facilitate screening differentiation conditions leading to cVO formation, we created a hESC-triple reporter line (hESC^TNNT2-GFP/CDH5-mOrange/TAGLN-CFP^) (hESC-3R) comprising the TNNT2 (troponin T) promoter-driven GFP to identify CMs, the CDH5 (VE-Cadherin) promoter-driven mOrange to identify ECs, and the TAGLN (SM22α) promoter-driven CFP to identify SMCs (**Fig. 3a**). Two lentiviral vectors, one for the CDH5 promoter-driven mOrange and the other for the TAGLN promoter-driven CFP (**Extended Data Fig. 3a-d**) were transduced into an hESC-TNNT2-GFP fluorescent reporter line^44^. We created two polyclonal and six monoclonal lines and confirmed pluripotency of the monoclonal lines (**Extended Data Fig. 3e-g**). Using differentiation protocols specific for CMs^7^, ECs^50^, and SMCs^11, 12^, we confirmed the differentiation of the hESC-3R line into CMs, ECs, and SMCs (**Fig. 3b**).

**Fig. 3.**
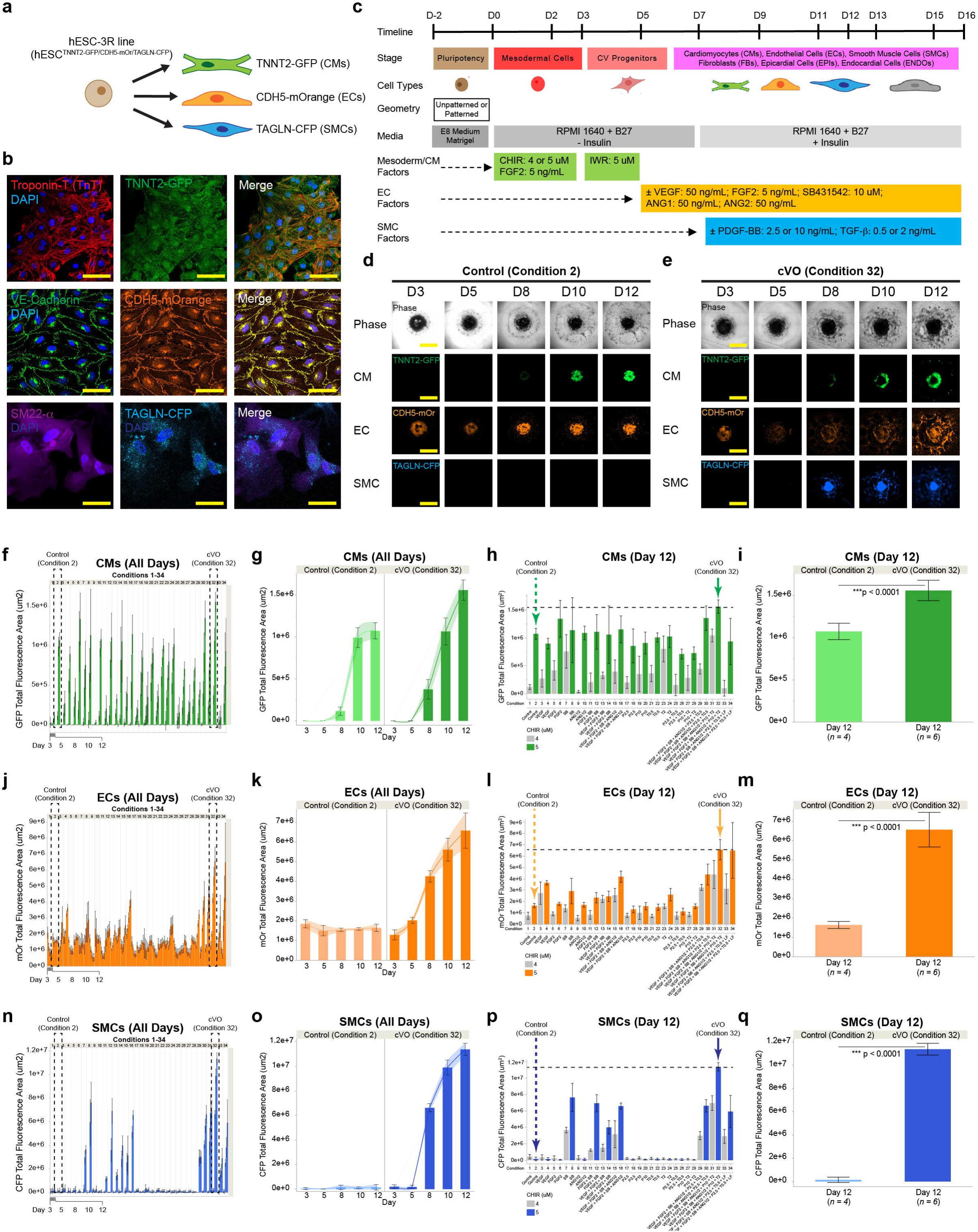
Creation of a triple-reporter line enables screening differentiation conditions leading to cVO formation. **a,** The hESC-3R line comprises the TNNT2 (Troponin-T) promoter driving GFP to identify CMs, the CDH5 (VE-Cadherin) promoter driving mOrange to identify ECs, and the TAGLN (SM22⍺) promoter driving CFP to identify SMCs. **b,** Schematic showing timeline, stage, cell types, geometry, media, and growth factors for co-differentiating CMs, ECs, SMCs, leading to formation of cVOs. **c,** Confocal fluorescence imaging and immunostaining confirms the differentiation of hESC-3R into CMs (co-expression of Troponin-T with TNNT2-GFP) (top row), ECs (co-expression of VE-Cadherin with CDH5-mOrange) (middle row), and SMCs (expression of TAGLN-CFP) (bottom row). Nuclei are stained with DAPI. Scale bar, 50 µm. **d-e,** Representative time-lapse imaging shows CM, EC, and SMC formation over 12 days for Control (Condition 2) and cVO (Condition 32). In the cVO sample, note ring formation of CMs, with outward radiation of ECs, and central SMCs. **f, j, n,** A total of thirty-four (34) differentiation conditions (n = 175 total micropatterns, n = 4-6 per condition) were screened for those giving the highest simultaneous co-differentiation of CMs, ECs, and SMCs. Half of the conditions used CHIR 4 µM (n = 88) and half used 5 µM (n = 87). Overall, 5 µM gave the most CMs, ECs, and SMCs. Student’s t-test, ***p < 0.0001. **f-i,** CM formation is highest in Condition 32 over time and begins around day 8. At day 12 CM formation is significantly higher for Condition 32 (n = 6) compared to Control** (n = 4). Student’s t-test, ***p < 0.0001. Error bars and shaded bands ± 1 SD. **j-m,** EC formation is highest in Condition 32 over time and is already apparent at day 3 and begins to increase around day 5. At day 12 EC formation is significantly higher for Condition 32 (n = 6) compared to Control** (n = 4). Student’s t-test, ***p < 0.0001. Error bars and shaded bands ± 1 SD. **n-q,** SMC formation is highest in Condition 32 over time and begins to increase around day 8. At day 12 SMC formation is significantly higher for Condition 32 (n = 6) compared to Control** (n = 4). Student’s t-test, ***p < 0.0001. Error bars and shaded bands ± 1 SD. *Condition 32 (cVO) consists of the following small molecules and growth factors: CHIR, 5 µM (day 0); FGF2, 5 ng/mL (days 0, 7, 9, 11, 13); IWR-1, 5 µM (day 3); VEGF, 50 ng/mL (days 5, 7, 9, 11, 13); SB 10 µM (days 7, 9, 11); ANG2, 50 ng/mL (days 5, 7); ANG1, 50 ng/mL (days 9, 11); PDGF-BB, 10 ng/mL (days 7, 9, 11, 13); and TGFβ1, 2 ng/mL (days 13). **Condition 2 (Control) is the standard cardiomyocyte differentiation with only the factors CHIR, FGF2, and IWR. Error bars ± 1 SD. See also **Supplementary Videos 3 and 4**.

Next, we tested the effects of CHIR concentrations (3, 4, and 5 µM) and VEGF addition (50 ng/mL) on CM and EC co-differentiation over 14 days using the hESC-3R line. hESC-3R micropatterns treated with CHIR + FGF2 + IWR-1 (Control) differentiated to the most CMs with a CHIR concentration of 5 µM and a few ECs at all concentrations (**Extended Data Fig. 2e**) in the absence of VEGF. On the other hand, hESC-3R micropatterns treated with 5 µM of CHIR and the addition of 50 ng/mL VEGF at days 6, 8, 10, and 12 of differentiation gave rise to the most CMs and ECs (**Extended Data Fig. 2f**). These initial results were the basis for our additional screening conditions described below for co-differentiating CMs, ECs, and SMCs to create cVOs. A schematic shows timeline, stage, cell types, geometry, media, and growth factors for creating CMs, ECs, SMCs, and cVOs (**Fig. 3c**). Our strategy for co-differentiation was initiated by first creating a cardiovascular (CV) progenitor pool that would give rise to all three cell types and then adding additional small molecules and growth factors to facilitate co-differentiation of CMs, ECs, and SMCs to produce cVOs (**Extended Data Fig. 3h**). We used previous established methods for guiding concentrations and timing for low and high production of CMs ^5, 7, 17^, ECs^10, 17, 51–53^, and SMCs^11, 54^ (**Extended Data Fig. 3i)**.

A total of thirty-four (34) screening conditions (n = 4-6 per micropatterns per condition, 175 total micropatterns) were tested to create various combinations of CMs, ECs, and SMCs in cVOs (**Supplementary Table 1, Extended Data Fig. 3j**). CHIR 4 µM were used in odd numbered conditions (n = 88 micropatterns) and CHIR 5 µM in even numbered conditions (n = 87 micropatterns) (**Extended Data Fig. 3k**).

Overall, CHIR 5 µM gave rise to more CMs (*p* < 0.0001), ECs (*p* < 0.0001), and SMCs (*p* < 0.0001) compared to CHIR 4 µM for each condition (**Extended Data Fig. 3l**; **Supplementary Video 3)**. Condition 32 produced cVOs with the most CMs, ECs and SMCs (**Fig. 3d-q,** and **Extended Data Fig. 3m-o**; **Supplementary Video 4**), which consists of the following small molecules and growth factors: CHIR, 5 µM (day 0); FGF2, 5 ng/mL (days 0, 7, 9, 11, 13); IWR-1, 5 µM (day 3); VEGF, 50 ng/mL (days 5, 7, 9, 11, 13); SB 10 µM (days 7, 9, 11); ANG2, 50 ng/mL (days 5, 7); ANG1, 50 ng/mL (days 9, 11); PDGF-BB, 10 ng/mL (days 7, 9, 11, 13); and TGFβ1, 2 ng/mL (days 13). Condition 2 (Control) is the baseline CM differentiation condition with only CHIR, FGF2, and IWR-1 being used.

Time-lapse microscopy showed more CM, EC, and SMC formation in Condition 32 compared to Control (**Fig. 3d-e**). Time-course CM formation for Control (n = 4) and Condition 32 (n = 6) showed that CMs began to emerge around day 8 for both conditions (**Fig. 3f-g**) and by day 12, there were statistically higher CMs in Condition 32 (*p* < 0.0001) (**Fig. 3h-i**). Time-course EC formation showed that ECs were already apparent at day 3 for both conditions and their formation began to increase around day 5 for Condition 32, a few days before CM formation (**Fig. 3j-k**); by day 12, there were statistically higher ECs in Condition 32 (*p* < 0.0001) (**Fig. 3l-m**). Time-course SMC formation showed that SMC formation began to increase around day 8 for Condition 32 (**Fig. 3n-o**) and by day 12, there were statistically higher SMCs in Condition 32 (*p* < 0.0001) (**Fig. 3p-q**).

As noted above, ECs CMs began to form around day 3 followed by CMs at day 8 in both conditions; in Condition 32, the rates of EC and SMC formation increased around days 5 and 8, respectively (**Extended Data Fig. 3p**). Interestingly, in terms of vascularization factor crosstalk with ECs and CMs, at day 12, VEGF alone generated the most ECs compared to Control (n =10, *p* < 0.0001), to FGF2 alone (n = 10, *p* < 0.0001), and to SB alone (n = 8, *p* < 0.05), while having no statistical differences on CM formation (**Extended Data Fig. 3q-r**). The combination of FGF2, SB, VEGF, along with the angiopoietins, produced the most ECs in Condition 32. For arrays of hESC-3R micropatterns, Condition 32 also gave rise to all the three cell types (**Extended Data Fig. 3s**).

We also found that the total fluorescence generated by the 34 screening conditions for cVO formation were well fit by simple machine learning models (general linear), encapsulating CM (R^2^=0.88; **Extended Data Fig. 7a**) and EC (R^2^=0.78; **Extended Data Fig. 7b**) formation over time. In summary, creation of the hESC-3R line enabled identification of differentiation conditions leading to robust cVO formation.

### cVOs comprise spatially and temporally self-organized cardiovascular cell types

We extended the differentiation of cVOs from 12 to 16 days (equivalent to ∼ 3 weeks of *in vivo* human development); these contained CMs, SMCs, and branching ECs arranged in a concentric fashion (**Fig. 4a**-**b**). We confirmed CMs co-expressed Troponin-T (TnT) and TNNT2-GFP, ECs co-expressed PECAM and CDH5-mOrange, and SMCs co-expressed Calponin and TAGLN-CFP (**Fig. 4c-e**). Compared to hESC-3R micropatterns, unpatterned hESC-3R cells resulted in unorganized and random distribution of CMs and ECs (**Extended Data Fig. 4a**). cVOs contained branched ECs intimately surrounding CMs and SMCs and moving in unison together with each CM contraction (**Extended Data Fig. 4b-d**; **Supplementary Videos 5** and **6**).

**Fig. 4.**
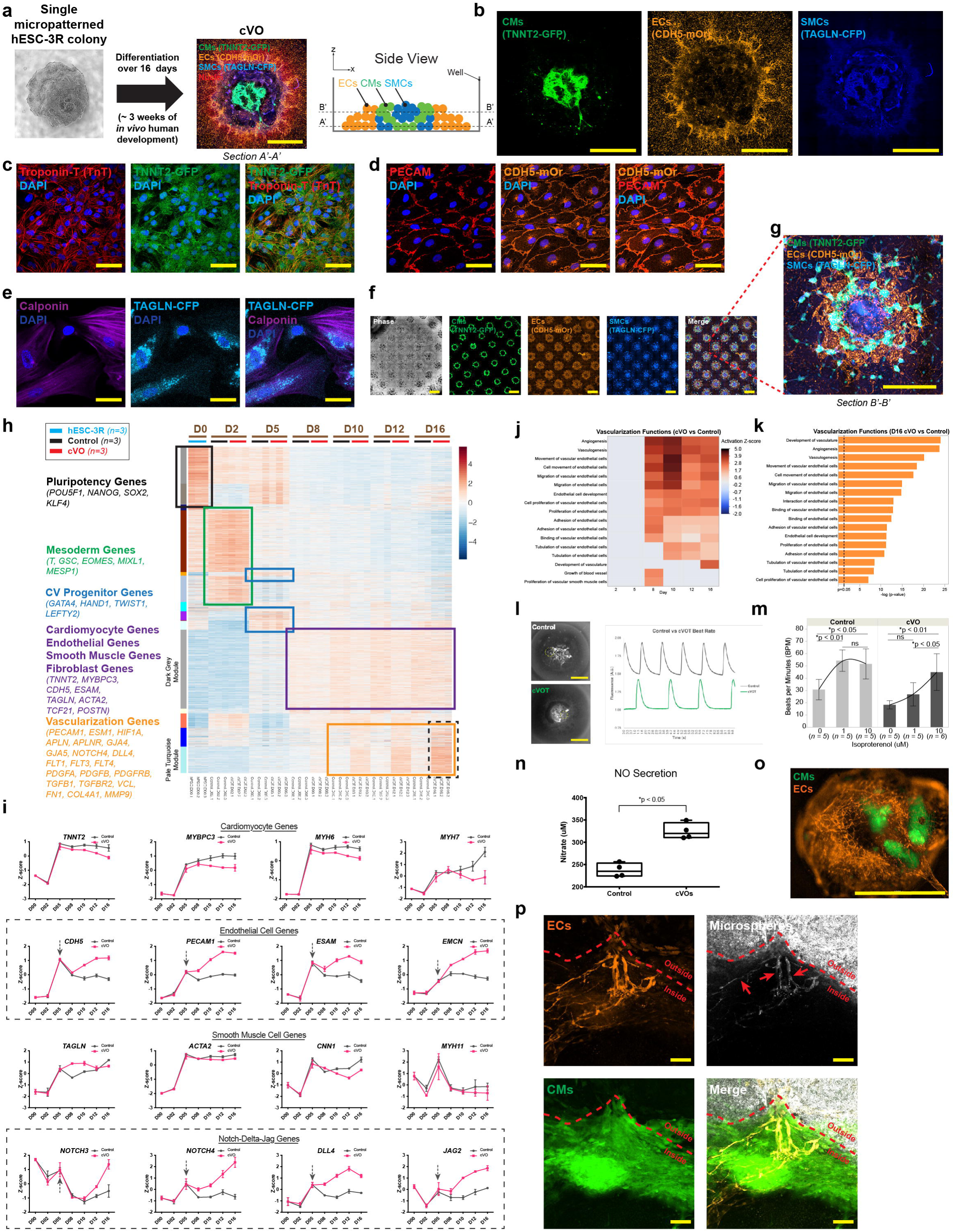
cVOs comprise spatially and temporally self-organized cardiovascular cell types. **a,** Confocal phase (left) and fluorescence (middle) imaging shows the differentiation of a single hESC-3R micropattern over 16 days (∼ 3 weeks of *in vivo* human development) into a cVO containing CMs (green), ECs (orange), and SMCs (blue). Nuclei are labeled with DRAQ5 nuclear stain (red). Note concentric organization of cell types and EC branching. This plane corresponds to section A’-A’, near the well bottom (right). Scale bar, 2 mm. **b,** Individual confocal channels showing CMs (green) (left), ECs (orange) (middle), and SMCs (blue) (right). Scale bar, 2 mm. **c,** Confocal fluorescence images showing hESC-3R CMs co-express Troponin-T (red) with TNNT2-GFP (green). Nuclei are labeled with DAPI (blue). Scale bar, 50 µm. **d,** Confocal fluorescence images showing hESC-3R ECs co-express PECAM (red) with CDH5-mOr (orange). Nuclei are labeled with DAPI (blue). Scale bar, 50 µm. **e,** Confocal fluorescence images showing hESC-3R SMCs co-express calponin (purple) with TAGLN-CFP (cyan). Nuclei are labeled with DAPI (blue). Scale bar, 50 µm. **f,** Phase (far left) and fluorescence imaging shows an array of hESC-3R micropatterns differentiated over 16 days into cVOs containing concentrically self-organized CMs (green), ECs (orange), and SMCs (blue). Note individual micropatterns give rise to an array of independently-formed cVOs despite sharing the same media. Scale bar, 2 mm. **g,** Enlarged area from red box in (**f**) showing concentric organization of CMs, ECs, and SMCs, with EC branching. This focal plane corresponds to section B’-B’ in (**a**). Scale bar, 2 mm. **h,** Temporal bulk RNA-sequencing (bRNA-seq) was performed for undifferentiated hESC-3R micropatterns (blue, n = 3), Control differentiation (black, n = 3), and cVO differentiation (red, n = 3). Weighted gene co-expression network analysis (WGCNA) heat map shows clusters of Pluripotency, Mesoderm, CV progenitor, Cardiomyocyte, Endothelial, Smooth Muscle, Fibroblast, and Vascularization Genes over 16 days (D0-16) for hESC-3R (blue, n = 3), Control (black, n = 3), and cVO (red, n = 3) groups. Note upregulated vascularization genes in D16 cVOs (black dashed rectangle). **I,** Comparison of CM, EC, SMC, and Notch-Delta-Jag gene groups from bulk RNA-seq most notably show that EC and Notch-Delta-Jag gene expression (dashed rectangles) is higher for cVOs and diverges from Controls around day 5 (dashed arrows), when vascular-inducing factors are added to the cultures. **j,** Ingenuity Pathway Analysis (IPA) of Days 2 to 16 cVOs compared to Controls showing a heat map of upregulated (red), equally regulated (gray), and downregulated (blue) vascularization functions. Note upregulation occurs between Days 5 and 8, when vascular-inducing factors were added. **k,** IPA of Day 16 cVOs compared to Controls showing significantly activated vascularization functions (-log (p-value) > 1.3). **l,** Fluo-4 calcium CM labeling of a Control (top) and cVO (bottom). Calcium transient rates of Controls are higher than cVOs (right). Note samples were differentiated from hiPSCs to avoid spectral overlap of Fluo-4 with GFP from the hESC-3R line. Scale bar, 2 mm. **m,** Beat rate of Controls (n = 15) and cVOs (n = 16) increases with isoproterenol (0, 1, and 10 uM) treatment, with Control rates consistently higher than cVO rates at each concentration. Two-way ANOVA with Tukey’s test for multiple comparisons, *p < 0.05 and *p < 0.01. **n,** hESC-3R EC nitric oxide (NO) secretion is higher in cVO ECs compared to Controls. Student’s t-test, *p < 0.05. **o,** 3D cVO differentiated over 16 days on a micropatterned substrate with a stiffness of 16 kPa showing CMs (green) surrounded by a branching network of ECs (orange). SMCs were not imaged. Scale bar, 1 mm. **p,** Microsphere (white) identification of vascular lumen (white vascular branches marked by red arrows) formed by ECs (orange). Note abundance of microspheres outside the cVO but only within vascular lumen inside the cVO. The red dashed line demarcates the outer cVO boundary, with CMs (green) shown inside the cVO. See also **Supplementary Videos 5**, **6**, and **7**.

Arrays of hESC-3R micropatterns also differentiated into cVOs and contained CMs, SMCs, and branched ECs arranged in a concentric fashion (**Fig. 4f**-**g**). Individual micropatterns gave rise to independent patterns of cVOs despite sharing the same media. Additionally, cVOs formed rings of CMs that beat in unison, both in a rotating and non-rotating fashion (**Supplementary Video 7**).

We performed temporal bulk RNA-sequencing (bRNA-seq) on I) single undifferentiated micropatterned hESC-3R colonies, ii) Controls, and iii) cVOs (n = 3 for all groups). Weighted gene co-expression network analysis (WGCNA) showed clustering of pluripotency, mesoderm, CV progenitor, CM, EC, SMC, and vascularization genes over 16 days. Vascularization genes were highest in cVOs and peaked at day 16 (**Fig. 4h**). Principal component analysis (PCA) showed developmental differences between all three groups over 16 days. Divergence between Control and cVO groups emerged at day 5, when vascular induction began (**Extended Data Fig. 4e**).

From our bRNA-seq analysis, we compared Control and cVO groups for select gene groups as follows: CM, EC, SMC, NOTCH-DELTA-JAG groups (**Fig. 4i**) along with pluripotent, mesoderm/CV progenitor, FB, and epicardial groups (**Extended Data Fig. 4f**). CM genes were higher for Controls compared to cVOs. For EC and NOTCH-DELTA-JAG genes, divergence of expression between Controls and cVOs began around day 5, when vascular specifying factors were added to the cultures. SMC and FB genes generally increased for both groups. Interestingly, the epicardial gene *WT1* was higher in cVOs than Controls, while the *TOP2A*, *UBE2C*, and *RRM2* genes decreased, suggesting a decrease in proliferation of epicardial cells in cVOs. As expected, pluripotent genes decreased, and mesoderm/CV progenitor genes increased in both Controls and cVOs.

Ingenuity Pathway Analysis (IPA) showed upregulated vascularization functions including angiogenesis and vasculogenesis (**Fig. 4j**). Upregulation over time generally began to increase at day 8, after vascularization factors were added in culture at day 5. IPA of day 16 cVOs vs Controls showed upregulation of vascularization functions (**Fig. 4k**).

In terms of cVO CM function, Fluo-4 calcium labeling of iPSC-derived Control and cVO groups revealed that calcium transient rates of Controls were higher than cVOs (**Fig. 4l**; **Supplementary Video 7**). Additionally, the beating rate of Controls (n = 15) and cVOs (n = 16) significantly increased with isoproterenol (0, 1, and 10 µM) treatment (*p* < 0.05 and *p* < 0.01 at 10 µM for Controls and cVOs, respectively) (**Fig. 4m**). Based on analysis of contraction-relaxation cycles of Controls and cVOs (**Extended Data Fig. 4g**), the beating rate, contraction velocity, relaxation velocity, and contraction-relaxation peak interval did not statistically differ (*p* = 0.65, 0.06, 0.10, and 0.52, respectively) between Controls (n = 4) and cVOs (n = 4) (**Extended Data Fig. 4h-k**). In contrast, EC nitric oxide (NO) secretion was significantly higher in cVO ECs compared to Controls (*p* < 0.05) (**Fig. 4n**). To create 3D cVOs we made micropatterns on a “soft” hydrogel substrate with a 16 kPa stiffness (compared to “hard” tissue culture plastic in the GPa range), which resulted in a network of ECs surrounding CMs and promoted spherical 3D formation (**Fig. 4o**). Additionally, we identified vascular branching and lumen within cVOs (**Fig. 4p**).

To demonstrate increased throughput and eliminate using stencils, we used the same differentiation protocol for 2D cVOs to create 3D cVOs over 14 days, using a range of CHIR at higher concentrations (6.0-8.5 µM) on a single hESC-3R colony formed in each well of a 96-well plate. CHIR 7.0 µM gave the most CMs and ECs, with CM and EC formation inversely related across all concentrations (**Extended Data Fig. 4l**). The higher CHIR needed in this format was not unexpected as we have previously needed to adjust CHIR concentrations depending on cell lines and multi-well plate formats. Finally, we validated our vascularization protocol with an hiPSC line (SCVI 113, Stanford CVI Biobank) and confirmed the presence of CMs, ECs and SMCs within hiPSC-derived cVOs (**Extended Data Fig. 4m-n**).

For cVOs, we also found a multitude of upregulated genes from our bRNA-seq analysis related to vascularization in EC, arterial, venous, endocardial, TGFβ pathway, VEGF pathway, PDGF pathway, gap junction, and paracrine gene groups (**Extended Data Fig. 5a-m**). Notably, we found that several members of the angiopoietin family *ANGPT2*, *ANGPTL1*, *ANGPTL4*, *ANGPTL6*, and *TIE1* were all upregulated in cVOs, consistent with their established role in vascularization^53, 55, 56^.

IPA showed upregulated Canonical Pathways also known to be important in cardiovascular development, including epithelial to mesenchymal transition (EMT), HIF1α signaling, and HOTAIR signaling (**Extended Data Fig. 6a**). IPA comparing Controls to cVOs from days 2-16 also showed upregulated Upstream Regulators known to be important in cardiovascular development, including *PDGFBB*, *TGFβ1*, *CD36*, *BMP4*, *VEGFA*, *FGF2*, and *HIF1A* (**Extended Data Fig. 6b**). Furthermore, IPA showed that the NOTCH pathway was upregulated in cVOs compared to Controls (**Extended Data Fig. 6c**), while the BMP pathway was upregulated in both groups (**Extended Data Fig. 6d**). IPA of the WGCNA Dark Grey module within the “Cardiomyocyte/Endothelial/Smooth Muscle/Fibroblast Genes” cluster in **Fig. 4h** confirmed a Cardiogenesis Regulator Effect Network with upstream regulators including *BMP*, *TGF*𝛽, and *WNT11* activating downstream genes including *MEF2C*, *NKX2-5*, *TBX5*, *MYH6*, and *NPPA,* leading to cardiogenesis (**Extended Data Fig. 6e**). IPA of the WGCNA Pale Turquoise module within the “Vascularization Genes” cluster in **Fig. 4h** confirmed a Vascularization Regulator Effect Network with upstream regulators including *VEGF*, *CD36*, *JAG2*, and *JUNB* activating genes including *PDGFB*, *FLT1*, *MMP9*, *TGF*𝛽*1*, *HIF1A*, *FN1*, and *DLL4*, leading to downstream Vascularization Functions (**Extended Data Fig. 6f**). IPA of day 16 cVOs vs Controls showed upregulated Vascularization Functions (**Extended Data Fig. 6g**) and a *VEGF* Regulator Effect network on *ITGA1*, *MMP9*, *CDH5*, *F3*, *NOTCH4*, *CD34*, and *DLL4,* leading to downstream development of vasculature (**Extended Data Fig. 6h**).

### Single-cell RNA-sequencing reveals multiple vascular, endocardial, myocardial, and epicardial cell types in cVOs

To obtain higher cellular resolution within Control and cVO groups, we next performed single-cell RNA-sequencing (scRNA-seq) and then primary downstream analysis with Seurat^57^. Non-linear dimensional reduction using UMAP of the cVO groups showed 9 clusters containing CMs, ECs, SMCs, FB, EPI, precursor cells (PRE), hepatic cells (HC), neural cells (NC), proliferating cells (PROLIF), and epithelial cells (EPT) (**Fig. 5a**), as also identified in UMAP feature plots (**Extended Data Fig. 8a**). Comparison with the cellular composition of a 6.5 post-conception week (PCW) (∼45 days) human heart (Carnegie Stages 19 and 20) from a recent study^46^ revealed that cVOs shared 9 of 14 cell types (**Fig. 5b**). This is the earliest *in vivo* scRNA-seq dataset to date but is still ∼3.5 weeks older than our cVOs and thus we expected our cVOs would not have all the cell types found in the 6.5 PCW heart. Further analysis revealed composition and lineage relationships of cellular subtypes of the 9 clusters (**Fig. 5c**). Notably, cVOs had 9% ECs, although based on our imaging data, we believe this is an underestimate and that a portion of ECs may have been lost during the dissociation process.

**Fig. 5.**
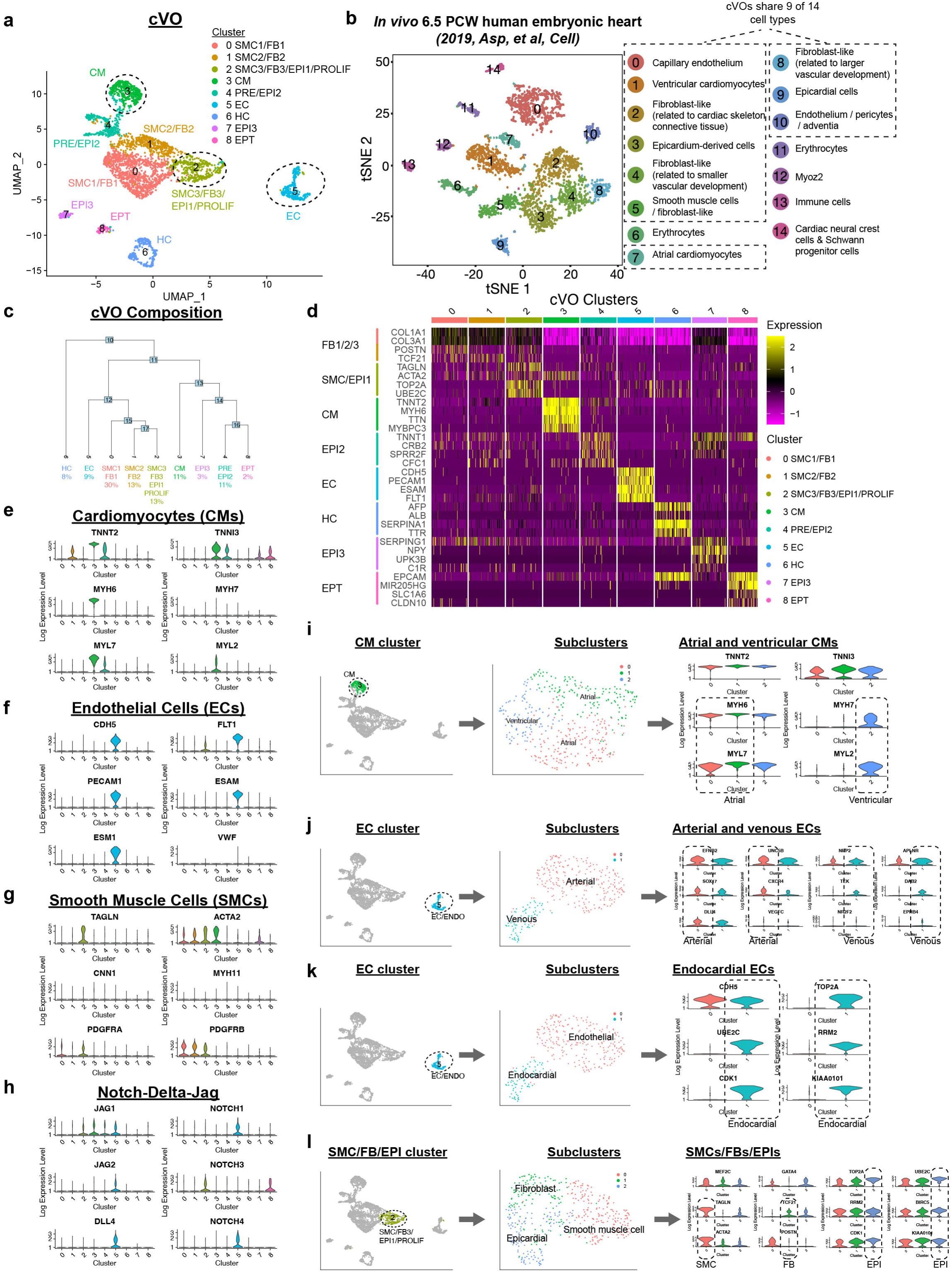
Single-cell RNA-sequencing reveals multiple vascular, endocardial, myocardial, and epicardial cell types in cVOs. **a,** cVO UMAP showing 8 clusters containing cardiomyocytes (CM), endothelial cells (EC), smooth muscle cells (SMC1, SMC2, SMC3), fibroblasts (FB1, FB2, FB3), epicardial cells (EPI1, EPI2, EPI3), precursor cells (PRE), proliferating cells (PROLIF), hepatic cells (HC), and epithelial cells (EPT). **b,** 6.5 PCW human heart t-SNE showing 14 cell types. cVOs share 9 of 14 cell types (dashed rectangles). PCW, post-conception week. **c,** cVO composition and lineage relationships of cellular subtypes from (**a**). **d,** cVO heatmap showing cell types in UMAP clusters in (**a**) according to scaled, log-normalized differentially expressed genes. **e-h,** cVO violin plots showing expression of CMs, ECs, SMCs, and NOTCH pathway genes in corresponding UMAP clusters in (**a**). **i,** cVO UMAP showing CM atrial (*MYH6*^+^, *MYL7*^+^) and ventricular (*MYH7*^+^, *MYL2*^+^) subtypes. **j,** cVO UMAP showing EC arterial (*EFNB2*^+^, *UNC5B*^+^, *SOX17*^+^, *CXCR4*^+^, *DLL4*^+^) and venous (*NRP2*^+^, *APLNR*^+^, *TEK*^+^, *DAB2*^+^, *EPHB4*^+^) subtypes. Note, the arterial subtype expresses a subset of venous genes and vice-versa. **k,** cVO UMAP showing EC (*CDH5^+^*) and endocardial (*TOP2A*^+^, *UBE2C*^+^, *RRM2*^+^, *CDK1*^+^, *KIAA0101*^+^) subtypes. **l,** cVO UMAP showing SMC (*TAGLN*^+^, *ACTA2*^+^), FB (*TCF21*^+^), and EPI (*TOP2A*^+^, *UBE2C*^+^, *RRM2*^+^, *BIRC5*^+^, *CDK1*^+^, *KIAA0101*^+^) subtypes.

We identified cell types in cVO UMAP clusters according to scaled, log-normalized differentially expressed genes (**Fig. 5d**). Violin plots of cVOs showed expression of CMs, ECs, SMCs, and NOTCH pathway genes (**Fig. 5e-h**) along with FBs, EPIs, HCs, PROLIFs, ECM, and VEGF pathway genes in corresponding UMAP clusters (**Extended Data Fig. 8b-g**). Violin plots also showed gene expression of cardiovascular progenitors (CV PROG), ECs, WNT/BMP/PDGF/TGFβ pathways, gap junctions (GJ) and ECMs (**Extended Data Fig. 8h**).

Subcluster analysis of CM cluster 3 revealed atrial (*MYH6^+^*, *MYL7^+^*) and ventricular (*MYH7*^+^, *MYL2*^+^) subtypes (**Fig. 5i**). Similarly, EC cluster 5 revealed arterial (*EFNB2*^+^, *SOX17*^+^, *DLL4*^+^, *UNC5B*^+^, *CXCR4*^+^) and venous (*NRP2*^+^, *TEK*^+^, *APLNR*^+^, *DAB2*^+^, *EPHB4*^+^) subtypes. Of note, the arterial subtype expressed a subset of venous genes and vice-versa (**Fig. 5j**). In addition, the EC cluster 5 revealed an endocardial (*CDH5*^+^, *UBE2C*^+^, *CDK1*^+^, *TOP2A*^+^, *RRM2*^+^, *KIAA0101*^+^) subtype (**Fig. 5k**). Finally, cluster 2 revealed SMC (*TAGLN*^+^, *ACTA2*^+^), FB (*TCF21*^+^), and EPI (*TOP2A*^+^, *RRM2*^+^, *CDK1*^+^, *UBE2C*^+^, *BIRC5*^+^, *KIAA0101*^+^) subtypes (**Fig. 5l**). In summary, scRNA-seq revealed multiple vascular, endocardial, myocardial, and epicardial cell types in cVOs, further improving the cellular subtype resolution identified by fluorescence imaging and bRNA-seq.

### Inhibition of NOTCH and BMP pathways disrupts vascularization within cVOs

The NOTCH pathway is vitally important for cardiovascular development, disease, and regeneration^58^. Based on our bRNA-seq and scRNA-seq data, members of the NOTCH pathway, including *JAG1*, *JAG2, DLL4, NOTCH1, NOTCH2, NOTCH3,* and *NOTCH4*, were upregulated in our cVOs (**Fig. 4i**, **Extended Data Fig. 6c, Fig. 5h,** and **Extended Data Fig. 8a**). Using our scRNA-seq data, we examined NOTCH-DLL-JAG receptor-ligand pairs between CMs, ECs, and SMCs to compare their putative cell-cell interactions in cVO and 6.5 PCW human heart groups (**Fig. 6a**). The number of receptors and ligands along with cell-cell interactions increased from cVOs to 6.5 PCW hearts, suggesting that increased NOTCH pathway activity is associated with increased vascularization and maturation.

**Fig. 6.**
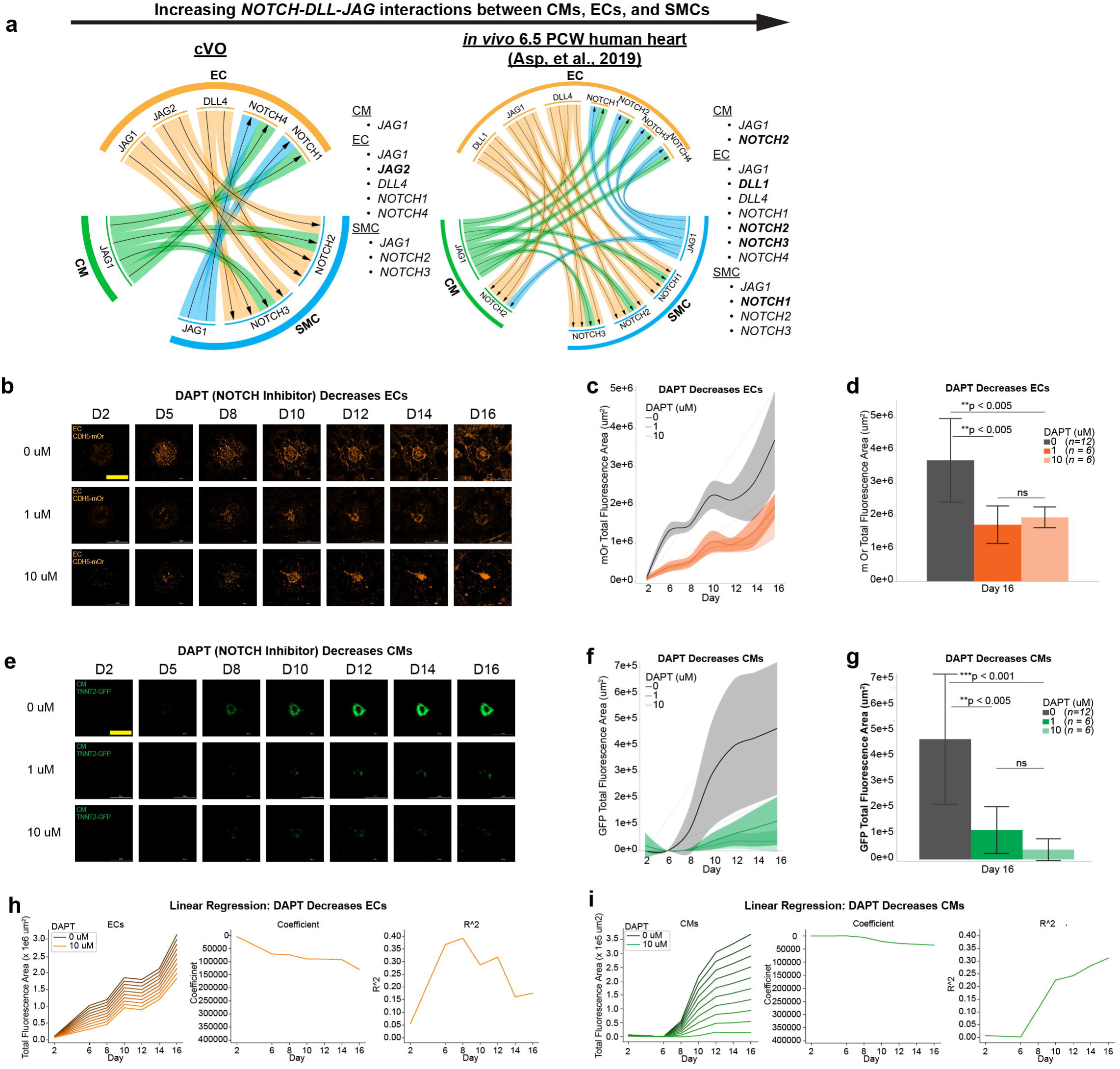
Inhibition of NOTCH pathway disrupts vasculature within cVOs. **a,** *NOTCH-DLL-JAG* receptor-ligand pair interactions between CMs, ECs, and SMCs are shown for cVO and an *in vivo* 6.5 PCW human heart^46^. cVO shows less receptors, ligands, and interactions than an *in vivo* 6.5 PCW heart. Bolded genes show differences between the cVO and heart groups, with the heart expressing more Notch receptors. PCW, post-conception week. **b-d,** DAPT, a NOTCH pathway antagonist, significantly decreased cVO EC formation over 16 days at 1 µM (n = 6, **p < 0.005) and 10 µM (n = 6, **p < 0.005) compared to 0 µM (Control) (n = 12). There was no significant (ns) difference between 1 and 10 µM. One-way ANOVA with Tukey’s test for multiple comparisons. Shaded bands and error bars ± 1 SD. Scale bar, 2 mm. **e-g,** DAPT significantly decreased cVO CM formation over 16 days at 1 µM (n = 6, **p < 0.005) and 10 µM (n = 6, ***p < 0.001) compared to 0 µM (Control) (n = 12). There was no significant (ns) difference between 1 and 10 µM. One-way ANOVA with Tukey’s test for multiple comparisons. Shaded bands and error bars ± 1 SD. Scale bar, 2 mm. **h-i,** Multiple linear regression model effects of DAPT on cVO EC and CM formation over 16 days (left) with linear regression coefficients (middle), and R^2^ (right). Based on the temporal graphs (**c, f, h, i**), the negative effect of DAPT on ECs was less than that on CMs.

To ascertain the effects of interfering with the NOTCH pathway on vascular and cardiac development, we next exposed cVOs to DAPT, a known NOTCH pathway antagonist. From days 2 to 16, DAPT decreased EC formation in a dose-independent manner (1 µM, n = 6, *p* < 0.005 and 10 µM, n = 6, *p* < 0.005) compared to (Control) (n = 12) (**Fig. 6b-d**). DAPT also significantly decreased CM formation at both 1 µM (n = 6, *p* < 0.005) and 10 µM (n = 6, *p* < 0.001) compared to 0 µM (Control) (n = 12) (**Fig. 6e-g**). In addition, we fit a simple machine learning model (multiple linear regression) to the changes in EC and CM formation caused by DAPT (**Fig. 6h-i**). Based on the temporal graphs (**Fig. 6 c, f, h, I**), the negative effect of DAPT on ECs was less than that on CMs.

Based on our bRNA-seq and scRNA-seq data, the BMP pathway was upregulated in cVOs. To ascertain the effects of interfering with the BMP pathway on vascular and cardiac development, we exposed cVOs to Dorsomorphin, a known BMP pathway antagonist. From days 2 to 16, Dorsomorphin decreased EC formation at 0.1 µM (n = 6, *p* < 0.0001) and 1.0 µM (n = 6, *p* < 0.0001) compared to 0 µM (Control) (n = 12). Similar to DAPT, a dose-dependent inhibitory effect of Dorsomorphin on EC formation was observed (**Extended Data Fig. 9a-c**). Dorsomorphin significantly decreased CM formation only at 1 µM (n = 6, *p* < 0.005) compared to 0 µM (Control) (n = 12), but 0.1 µM Dorsomorphin showed no significant inhibitory effects on CM formation (**Extended Data Fig. 9d-f**). In parallel, we also fit a multiple linear regression model to the changes in EC and CM formation caused by Dorsomorphin (**Extended Data Fig. 9g-h**). Based on the temporal graphs (**Extended Data Fig. 9b, e, g, h**), the negative effect of Dorsomorphin on ECs was more significant than that on CMs; this was the opposite for DAPT. The co-differentiation ECs and CMs in cVOs enabled this finding.

Fentanyl, a potent opioid agonist, has contributed to the opioid epidemic in the US in recent years^59^ and has the potential to be misused during pregnancy, possibly leading to congenital malformations as a teratogen. Since fentanyl has been shown to activate multiple pro-angiogenic signaling pathways^60^, we sought to observe any similar effects on cVO vascularization. Interestingly, at day 16, fentanyl significantly increased EC formation at 10 nM (n = 18, *p* < 0.001) compared to 0 nM (Control) (n = 18) (**Extended Data Fig.9i-j**), an opposite finding to DAPT and Dorsomorphin above.

### Vascularization factors used for creating cVOs enable creation of hVOs

We next asked if our vascularization strategy for cVOs could be applied to the hepatic system. Based on our results from hESC-RUES-GLR micropatterning, bRNA-seq, and scRNA-seq, we knew that we could produce mesoderm and endoderm, which we reasoned would be necessary for vascular and hepatic co-differentiation. Thus, our strategy for co-differentiation of ECs, SMCs, and HCs to create hVOs consisted of inducing mesendoderm and then co-differentiating a vascular progenitor pool and hepatoblast pool that would give rise to all the three cell types (**Extended Data Fig. 10a**). A schematic shows our timeline, stages, cell types, geometry, media, and growth factors for creating hVOs (**Extended Data Fig. 10b**). We used previous established methods to guide mesendoderm, foregut progenitors, hepatoblasts, and hepatocytes^3, 8^ (**Extended Data Fig. 10c**). We tested 3 differentiation conditions as follows: i) Control (day 20 baseline hepatic differentiation, with no vascularization factors added); ii) hVO-D3 (day 20 hVOs created by adding vascularization factors at day 3 of differentiation); and iii) hVO-D6 (day 20 hVOs created by adding vascularization factors at day 6 of differentiation) (**Supplementary Table 2**). We used the hESC-3R line to temporally observe vascular formation and then performed end-point immunostaining for hepatic markers; when applying our hepatic differentiation conditions, we did not observe any GFP fluorescence, thus indicating that we were not creating any “contaminating” CMs.

PCA of temporal bRNA-seq showed developmental differences between hESC-3R (day 0 undifferentiated micropatterns), Control, hVO-D3, and hVO-D6 groups (n = 3 for each group). A divergence between “Vascularization Factors” (D3 and D6 groups) versus “No Vascularization Factors” (Control group) was notable (**Extended Data Fig. 10d**). WGCNA showed clustering of pluripotency, structural, miscellaneous hepatic genes, hepatic and smooth muscle cell genes, and vascularization genes. Vascularization genes were most upregulated in the hVO-D3 group (**Extended Data Fig. 10e**).

From our bRNA-seq analysis, we compared the three groups for select gene groups as follows: pluripotent, HC, EC, SMC, FB, ECM, and NOTCH-DELTA-JAG (**Extended Data Fig. 10f-m**). Overall, pluripotency genes were upregulated for the hESC-3R group and downregulated for the Control, hVO-D3, and hVO-D6 groups. Hepatic genes were upregulated for all differentiation groups. Notably, all EC, SMC, ECM, and NOTCH-DELTA-JAG genes (except *DLL3* and *JAG1*) were most upregulated for the hVO-D3 group.

From days 3 to 19, time-lapse fluorescence microscopy showed EC formation for all three groups (n = 10, 12, 10, respectively). Adding vascular factors at day 3 (hVO-D3 group) to the baseline hepatic differentiation protocol gave the most EC formation at day 19 (*p* < 0.0001) (**Extended Data Fig. 10n-p**). Additionally, and importantly, confocal fluorescence imaging demonstrated the co-differentiation of HCs and ECs at day 19 of differentiation (**Extended Data Fig. 10q**). Thus, taken together, adding vascular factors at day 3 to developing hepatocytes resulted in the highest vascularization of hepatic organoids.

## Discussion

Using clues from developmental biology, we are able to create spatially self-organized 2D and 3D cVOs from hPSC micropatterns by identifying a combination of growth factors that simultaneously give rise to a spatially organized and branched vascular network within endocardial, myocardial, epicardial, and progenitor cells, along with numerous ECM proteins. Using scRNA-seq, we show similar cellular composition of cVOs to a 6.5 PCW human heart. Furthermore, we use machine learning to characterize key signals that affect CM and EC formation efficiency. We find that NOTCH and BMP pathways are upregulated in cVOs, and inhibition of these pathways disrupts vascularization. Finally, using the same vascular-inducing factors to create cVOs, we produce hVOs.

Spatial micropatterning of hPSCs has been shown by us and others to enable repeatable and scalable formation of spatially organized germ layers^40, 48^, primitive streak^43, 49^, cardiomyocytes^22, 23^, and cardiac organoids^24^. In our study here, we combined micropatterning with four fluorescent reporter cell lines to systematically phenotype conditions leading to simultaneous co-differentiation of self-organized CMs, ECs, and SMCs. Importantly, these reporter lines enabled temporal identification of germ layer, progenitor, and cardiovascular cell types *in situ* without disturbing the architecture of developing cVOs.

Recently, several impressive studies have shown the creation of 3D cardiac organoids^25–29^ with ECs and ENDOs lining central chambers. However, these vascular cell types were mostly found incidentally through transcriptomics and immunostaining; efforts to increase their numbers have been mostly with the addition of VEGF^17^. In our screening conditions, we added FGF2, SB, ANG1, ANG2, PDGF-BB, and TGFβ1 to systematically promote vasculogenesis, followed by angiogenesis, vessel branching, and lumen formation. Furthermore, our hESC-3R line enabled temporal recording of this developmental process, while micropatterning enabled spatial relationships to be identified. FGF2^61^, combined with VEGF and SB has recently been shown to produce vascular organoids^31, 62^. VEGF promotes vasculogenesis, while the addition of FGF2 with VEGF promotes angiogenesis^52^, and the addition of SB to these two factors increases hPSC-EC differentiation by up to 36-fold^10^. Additionally, ANG2 also promotes angiogenesis, while subsequent addition of ANG1 has been shown to promote vessel maturation^51, 52, 55^. Highlighting the contribution of the angiopoietins to cVO vascularization, we found that *ANGPT2*, *ANGPTL1*, *ANGPTL4*, *ANGPTL6*, and *TIE1* were all upregulated in cVOs. Finally, the addition of PDGF-BB has been shown to recruit perivascular cells (both pericytes and SMCs) to wrap around angiogenic sprouts, and addition of TGFβ1 has been shown to lead to vessel maturation by inhibiting EC proliferation, stimulating SMC differentiation, and promoting ECM deposition^51, 52^. In support of this, we found that several members of the PDGF and TGFβ pathways were upregulated in our cVOs and correlated with increased vascularization.

Our temporal bulk and endpoint single-cell transcriptomics revealed several cell types and pathways known to be important for cardiogenesis and vasculogenesis^63^, including the NOTCH, and BMP pathways. Our interference with relatively non-specific inhibitors of these pathways (DAPT, Dorsomorphin) all revealed their general disruptive effects on vascularization; the next step is to use more specific methods such as single-cell ATAC-seq, CRISPR activation/inhibition, CRISPR-mediated gene knockouts, and RNA-interference to dissect these pathways and cellular crosstalk even further to understand the etiology of congenital heart disease. For example, our system, with both cardiac and vascular cell types, could be used to further dissect the role of NOTCH1 in hypoplastic left heart syndrome (HLHS)^64^, the role of the TGFβ pathway in left ventricular non-compaction (LVNC)^65^, and the role of the PDGF pathway in lamin A/C (LMNA) cardiomyopathy^66^. All these pathways were upregulated in our cVOs, indicating their important role in cardiogenesis and vascularization.

Finally, using the same combination of vascular-inducing factors to create cVOs and hVOs implies a conserved developmental program for creating vasculature in different organ systems. With the RUES-GLR germ layer reporter line, various progenitor reporter lines, and parenchymal cell-specific reporter lines, we believe our vascularization method will serve as a starting point that can be added to established cell-specific differentiation protocols to achieve vascularization in other organ systems. Organoid vascularization will be necessary to i) avoid necrosis in the center of organoids where oxygen tension is low *in vitro* and *in vivo*^35^, ii) achieve larger organoid growth *in vitro* for improved systems for disease modeling, drug toxicity testing, and drug discovery^34^, and iii) increase the viability of implanted organoids for regenerative medicine applications *in vivo*^36^. See the **Extended Discussion** (Supplementary Information) for our suggested strategies for achieving improved vascularized organoid models moving forward.

## Acknowledgements

This work was supported by National Institutes of Health (NIH) K01 HL130608 (OJA); American Heart Association (AHA) Postdoctoral Fellowship Award 18POST34030106 (HY); K08 HL119251 (KDW); R01 HL150693, R01 HL141371, R01 HL146690, R01 HL145676, P01 HL141084 (JCW); California Institute for Regenerative Medicine (CIRM) Grant RC1-00151 (CKZ); Stanford Maternal & Child Heath Research Institute (MCHRI) Transdisciplinary Initiatives Program (TIP) Grant (OJA); Stanford Cardiovascular Institute (CVI) Seed Grant (OJA and HY); the Stanford Bio-X Program (OJA); and the Tobacco-Related Disease Research Program (TRDRP) 30FT0852 (MS). We thank the Stanford Neuroscience Microscopy Service (NMS) (Andrew Olson, Gordon Wang), the Stanford Cell Sciences Imaging Facility (CSIF) (Jon Mulholland, Kitty Lee), the Stanford Genomics (SG) Facility (John Coller, Dhananjay Wagh), the Stanford Genetics Bioinformatics Service Center (Ramesh Nair, Yue Zhang), the Computational Services and Bioinformatics Facility (CSBF) (Lee Kozar), the Stanford Protein and Nucleic Acid (PAN) Facility, the Stanford Neuroscience Gene Vector and Virus Core (GVVC), the Stanford Department of Physics Student Machine Shop (Mehmet Solyali, Karlheinz Merkle) for all their technical support, and Kevin Lam and Renata Tully for help with confocal imaging. OJA thanks Luke Abilez for several fruitful discussions on various aspects of this study. We thank Daylon James and Shahin Rafii (Reproductive Endocrinology Laboratory and Ansary Stem Cell Institute, Weill Cornell Medical College) for the pLV-hVPr-mOrange plasmid. We thank Ali Brivanlou (Laboratory of Synthetic Embryology, Rockefeller University) for the Human RUES2 GLR hESC line.

## Author Contributions

OJA conceived the project, designed experiments, performed experiments, analyzed data, interpreted results, and wrote the manuscript. HY performed experiments, analyzed data, and interpreted results. OJA, LT, and GZ performed bioinformatics analyses. KDW and EL designed and performed machine learning algorithms. MS, YZ, FJ, HTW, YG, BA, and DO performed experiments, analyzed data, and/or interpreted results. OJA, GP, CKZ, and JCW supervised the project. All authors reviewed the manuscript and agreed that the results support the conclusions of the project.

## Corresponding Authors

Correspondence to Oscar J. Abilez or Joseph C. Wu.

## Ethics Declarations

An invention disclosure has been filed with the Stanford Office of Technology Licensing (OTL). OJA is a consultant for Rosebud Biosciences.

## Extended Data Figure Legends

**Extended Data Fig. 1.**
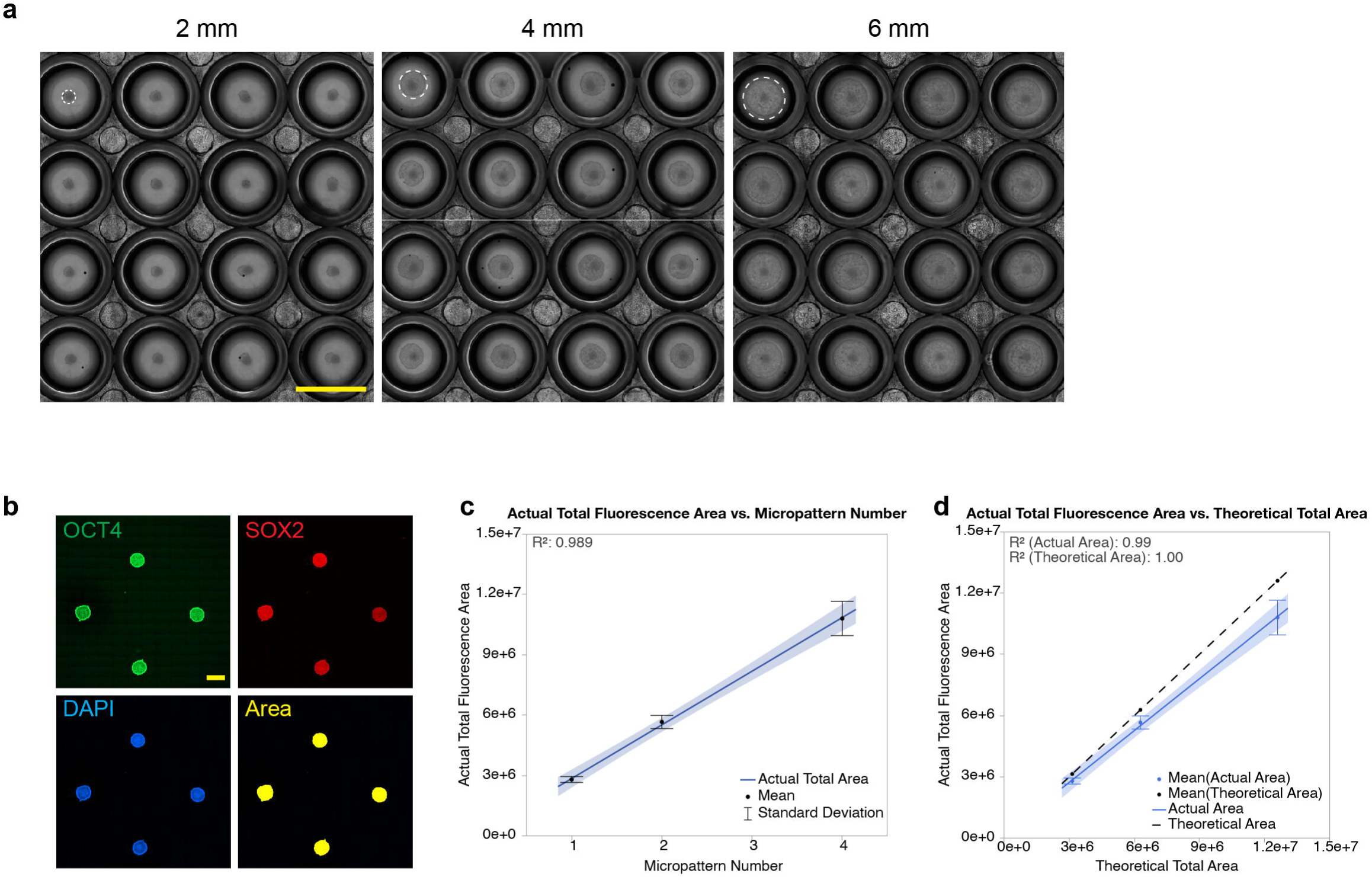
Micropatterning of hPSCs results in spatially organized cardiomyocytes, related to Fig. 1. **a,** Phase images showing reproducible 2 mm (left), 4 mm (middle), and 6 mm (right) undifferentiated hPSC micropatterns in 16 wells of a 48-well plate. Representative micropatterns are outlined by white dashed circles. Scale bar, 10 mm. **b,** Fluorescence imaging showing an array of four 2 mm hPSC micropatterns are pluripotent by immunostaining for OCT4 (green), SOX2 (red), and nuclear staining with DAPI (blue). The total fluorescence area of DAPI of each micropattern is shown in yellow. Scale bar, 2 mm. **c,** The actual total fluorescence area of the micropatterns correlates with micropattern count (R^2^ = 0.989) (n = 4 per micropattern count). Error bars ± SD. **d,** The actual total fluorescence area of the micropatterns correlates with the theoretical total micropattern area (multiples of pi*r^2^, where r is the radius of each micropattern) (R^2^ = 0.99) (n = 4 per micropattern count). Error bars ± SD.

**Extended Data Fig. 2.**
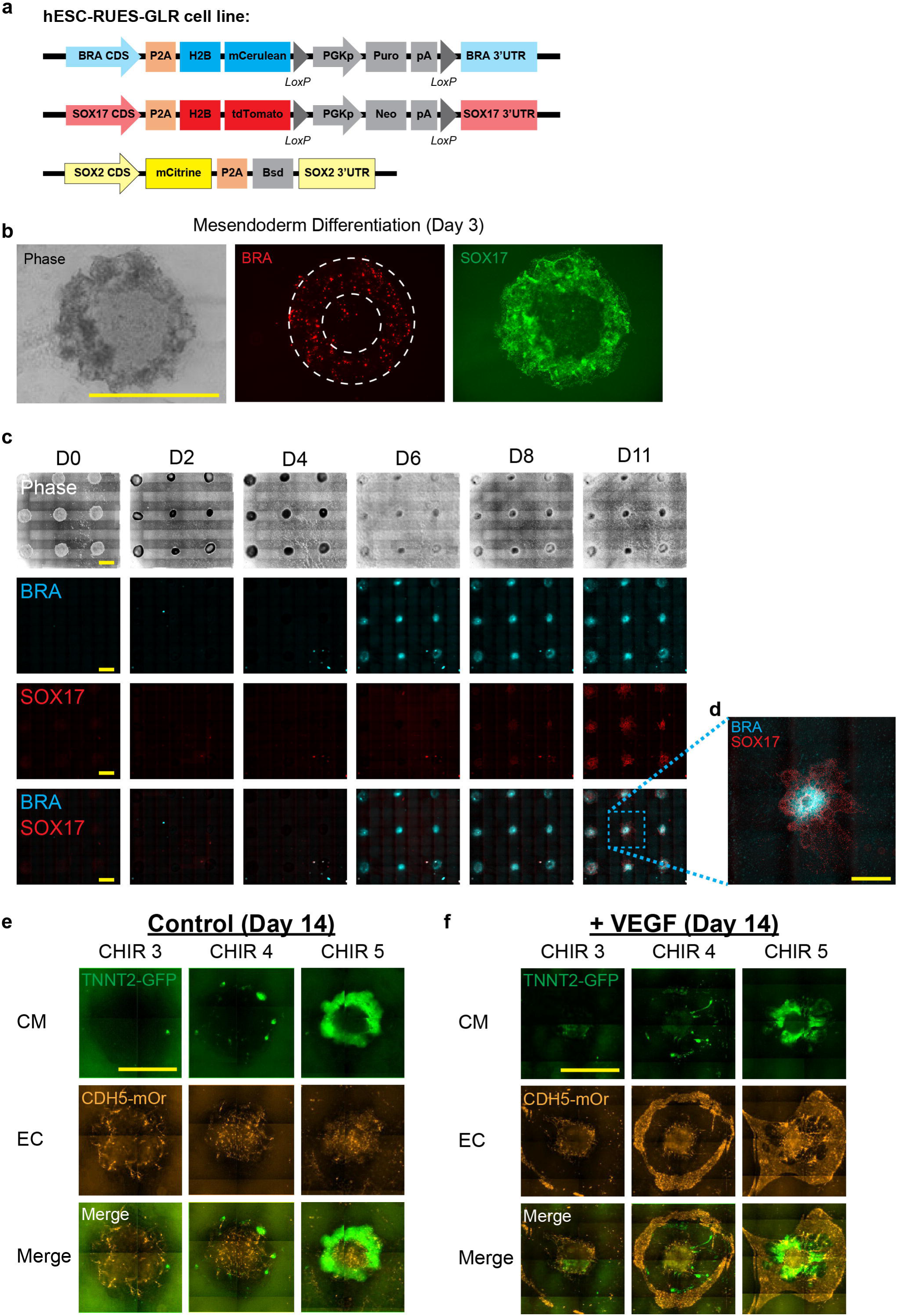
Micropatterning gives rise to organized germ layer formation, related to Figs. 2 and 3. **a,** The hESC-RUES-GLR (germ layer reporter) cell line ^43^ used to visualize germ layer formation comprises BRA driving mCerulean expression to visualize mesoderm formation, SOX17 driving tdTomato expression to visualize early endoderm formation (and late arterial formation), and SOX2 driving mCitrine to visualize ectoderm formation and pluripotency. **b,** Fluorescence images showing mesendoderm differentiation of a 2 mm hPSC micropattern at day 3 with circular distribution (white circular dashes) of BRA^+^ (mesoderm, red) and SOX17^+^ (endoderm, green). Scale bar 2 mm. **c,** Phase and fluorescence images showing temporal expression from days 0 to day 11 (D0-11) of the mesodermal marker BRA and the endodermal/arterial marker SOX17 of micropatterned arrays of RUES-GLR hESCs. Scale bar, 2 mm. **d,** Enlarged view of dashed square in (**c**). Note BRA^+^ central distributed expression relative to outer SOX17^+^ expression. Scale bar, 2 mm. **e,** Fluorescence images showing that hESC-3R (described in Fig. 2 and **Extended Data Fig. 3**) micropatterns treated with CHIR + FGF2 + IWR1 (Control) differentiated to the most CMs (green) at day 14 with a CHIR concentration of 5 µM. ECs (orange) are present at all concentrations but are punctate, with minimal branching. **f,** Fluorescence images showing that hESC-3R micropatterns treated with CHIR + FGF2 + IWR1 + VEGF (+VEGF at days 5, 7, 9, and 11) co-differentiated to the most CMs (green) and ECs (orange) at day 14 with a CHIR concentration of 5 µM and VEGF concentration of 50 ng/mL.

**Extended Data Fig. 3.**
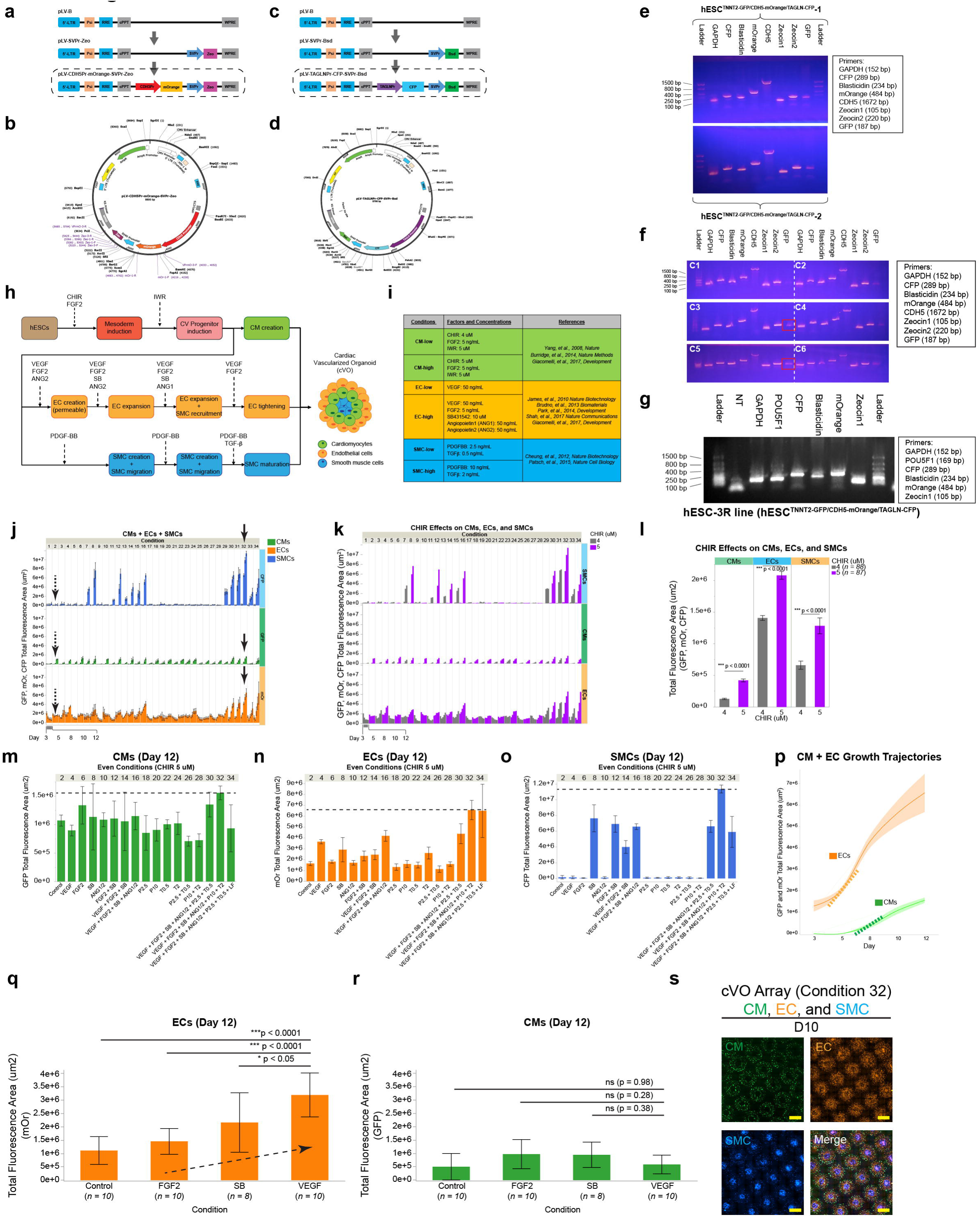
Creation of a triple-reporter line enables screening of differentiation conditions leading to cVO formation, related Fig. 3. **a,** Strategy for creating the lentivector pLV-CDH5Pr-mOrange-SVPr-Zeo used to create the hESC-3R line. The CDH5 (VE-Cadherin) promoter drives mOrange to identify ECs. **b,** Plasmid map of pLV-CDH5Pr-mOrange-SVPr-Zeo. **c,** Strategy for creating the lentivector pLV-TAGLNPr-CFP-SVPr-Bsd used to create the hESC-3R line. The TAGLN (SM22a) promoter drives CFP to identify SMCs. **d,** Plasmid map of pLV-TAGLNPr-CFP-SVPr-Bsd. **e,** Gel showing genomic PCR products of two polyclones of the hESC-3R line (hESC^TNNT2-GFP/CDH5-mOrange/TAGLN-CFP^ −1 and −2). A (DNA) Ladder, GAPDH (housekeeping gene), CFP, Blasticidin, mOrange, CDH5, Zeocin1, Zeocin2, and GFP are shown. **f,** Gel showing genomic PCR products of six monoclones (C1-C6) of the hESC-3R line. C3 and C5 (red rectangles) show the highest signals for all products. A (DNA) Ladder, GAPDH (housekeeping gene), CFP, Blasticidin, mOrange, CDH5, Zeocin1, Zeocin2, and GFP are shown. **g,** Gel showing genomic PCR products of a monoclonal hESC-3R line. A (DNA) Ladder, non-template (NT) control, GAPDH (housekeeping gene), POU5F1, CFP, Blasticidin, mOrange, and Zeocin1 are shown. **h,** Strategy for co-differentiating CMs, ECs, and SMCs to create cVOs. A key step is creating a cardiovascular (CV) progenitor pool that gives rise to all three cell types. **i,** Growth factor and small molecule characteristics for differentiation into CMs, ECs, and SMC. Select references are listed that were used as guidelines for concentrations and timing used in thirty-four (34) screening conditions. The detailed screening conditions are listed in **Supplementary Table 1j,** The thirty-four (34) screening conditions (n = 175 total micropatterns) showing total fluorescence area for CMs (GFP), ECs (mOr), and SMCs (CFP) from days 0-12 for each condition. Odd numbered conditions (1-33) used CHIR 4 µM and even numbered conditions (2-34) used 5 uM. Overall, Condition 32 (black solid arrows) gave rise to the most CMs, ECs, and SMCs. Condition 2 (black dashed arrows) is the standard Control CM differentiation with only the small molecules CHIR, FGF2, and IWR1. Error bars ± 1 SD. See also **Supplementary Videos 3** and **4**. **k,** Conditions using CHIR 4 (grey, n = 88 micropatterns) or 5 uM (purple, n = 87 micropatterns) showing largest CM, EC, and SMC formation (GFP, mOr, CFP total fluorescence area). **l**, Overall, 5 µM gave rise to the most CMs, ECs, and SMCs compared to 4 uM. Error bars ± 1 SD. **m,** Even conditions (2-34) using 5 µM (green) (n = 4-6 per condition) showing largest CM formation (GFP total fluorescence area) at day 12 for Condition 32 (dashed line). Error bars ± 1 SD. **n,** Even conditions (2-34) using 5 µM (orange) (n = 4-6 per condition) showing largest EC formation (mOr total fluorescence area) at day 12 for Condition 32 (dashed line). Error bars ± 1 SD. **o,** Even conditions (2-34) using 5 µM (blue) (n = 4-6 per condition) showing largest SMC formation (CFP total fluorescence area) at day 12 for Condition 32 (dashed line). Error bars ± 1 SD. **p,** Time-course CM formation (GFP total fluorescence area) and EC formation (mOr total fluorescence area) for Conditions 32 (n = 6). The rate of CM formation (dashed green line) increases around day 8 while the rate of EC formation (dashed orange line) increases around day 5. Note EC formation is already apparent at day 3. Shaded bands ± 1 SD. See also **Supplementary Videos 3** and **4**. **q,** At day 12, VEGF alone gave the most ECs compared to Control (n =10, ***p < 0.0001), to FGF2 alone (n = 10, ***p < 0.0001), and to SB alone (n = 8, *p < 0.05). **r,** FGF2, SB, and VEGF had no statistical (ns) effect on CM formation. The combination of FGF2, SB, VEGF, along with the angiopoietins, gave the most ECs in Condition 32 (see **n**). One-way ANOVA with Tukey’s test for multiple comparisons for GFP and mOrange channels separately. **s,** Fluorescence images showing cVO array formation at day 10 comprising CMs (green), ECs (orange), and SMCs (blue). Scale bar, 2 mm.

**Extended Data Fig. 4.**
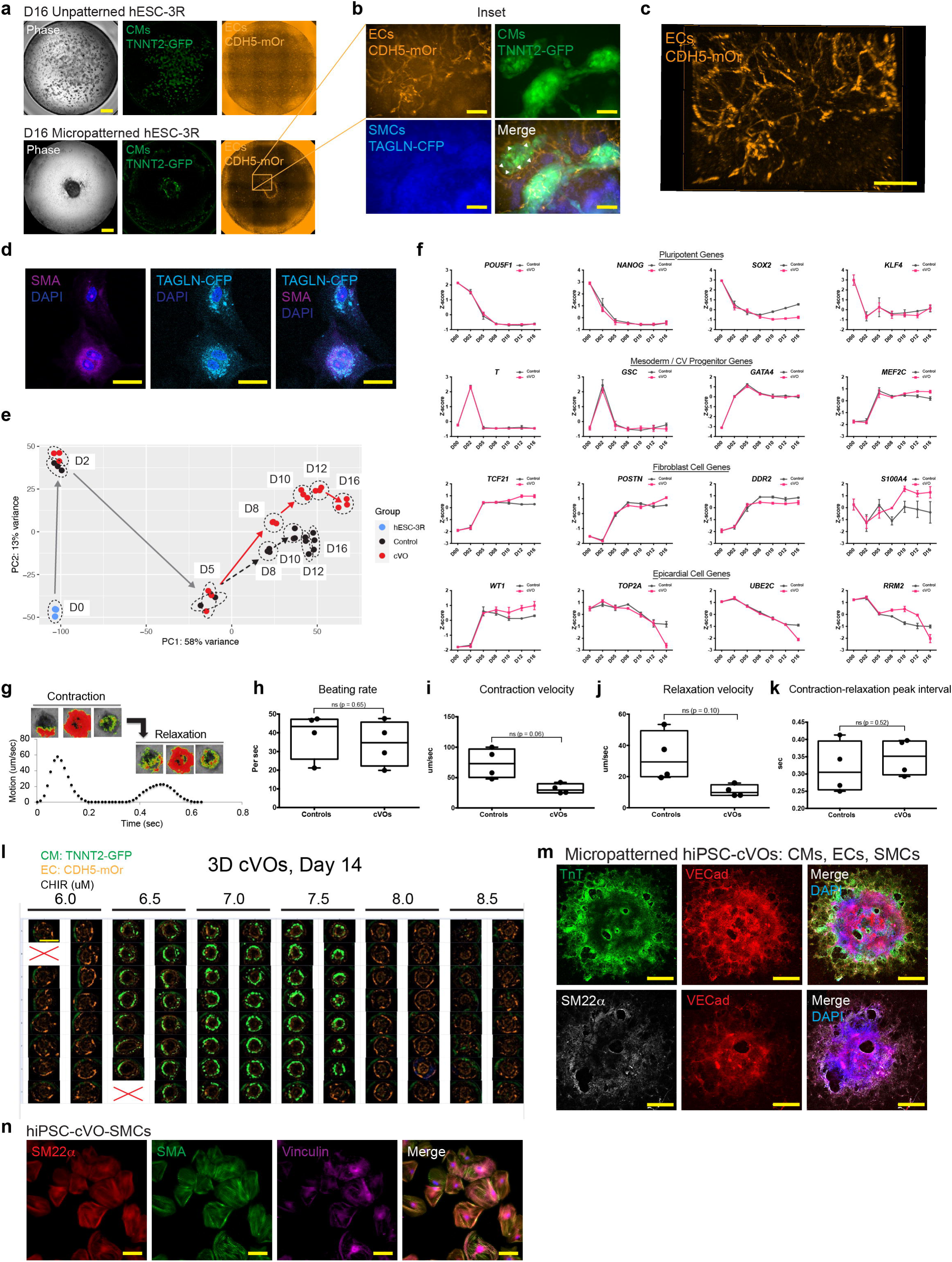
cVOs comprise spatially and temporally self-organized cardiovascular cell types, related to Fig. 4. **a,** Phase and fluorescence images showing the differentiation of unpatterned (top row) and micropatterned (bottom row) hESC-3R cells over 16 days containing CMs (TNNT2-GFP, green) and ECs (CDH5-mOr, orange). Note the unorganized distribution of CMs and ECs in the unpatterned group. Scale bar, 2 mm. **b,** An enlarged inset from the black rectangle in (**a**) showing CMs, ECs, and SMCs. White arrowheads show ECs surrounding a group of CMs. Scale bar, 500 µm. **c,** Maximum intensity image of ECs in (**b**) shows branching. Scale bar, 500 µm. **d,** Confocal fluorescence images showing hESC-3R SMCs co-express smooth muscle actin (SMA) (purple) with TAGLN-CFP (cyan). Nuclei are labeled with DAPI (blue). Scale bar, 50 µm. **e**, Principal component analysis (PCA) shows differences in differentiation trajectories between undifferentiated hESC-3R micropatterns (blue, n = 3), Control differentiation (black, n = 3), and cVO differentiation (red, n = 3) over 16 days (D0-16). Note trajectory divergence begins at day 5, when vascular induction begins. **f,** Comparison of Pluripotent, Mesoderm/CV Progenitor, Fibroblast, and Epicardial gene groups from bulk RNA-seq trend as expected. **g,** Representative contraction-relaxation cycle of a single cVO (shown in the three images above each phase, with maximum contraction and maximum relaxation shown in red). **h,** Beating rate of Controls (n = 4) was higher than cVOs (n = 4), but not statistically significant (p = 0.65). Student’s t-test, p < 0.05 considered significant. **i,** Contraction velocity of Controls (n = 4) was higher than cVOs (n = 4), but not statistically significant (p = 0.06). Student’s t-test, p < 0.05 considered significant. **j,** Relaxation velocity of Controls (n = 4) was higher than cVOs (n = 4), but not statistically significant (p = 0.10). Student’s t-test, p < 0.05 considered significant. **k,** Contraction-relaxation peak interval of Controls (n = 4) was lower than cVOs (n = 4), but not statistically significant (p = 0.52). Student’s t-test, p < 0.05 considered significant. **l,** 3D cVOs, day 14 created with varying amounts of CHIR (6.0-8.5 µM) with 7.0 µM giving the most CMs (green) and ECs (orange). n = 15-16 per group. **m,** A micropatterned hiPSC-derived cVO (Day 16) immunostained and confocal imaged for the CM marker Troponin-T (TnT), the EC marker VE-Cadherin (VECad), and the SMC marker smooth muscle 22α (SM22α). Nuclei are labeled with DAPI (blue). Scale bar, 1 mm. **n,** Confocal fluorescence images showing hiPSC-SMCs co-express SM22α (red), SMA (green), and vinculin (purple). Nuclei are labeled with DAPI (blue). Scale bar, 50 µm.

**Extended Data Fig. 5.**
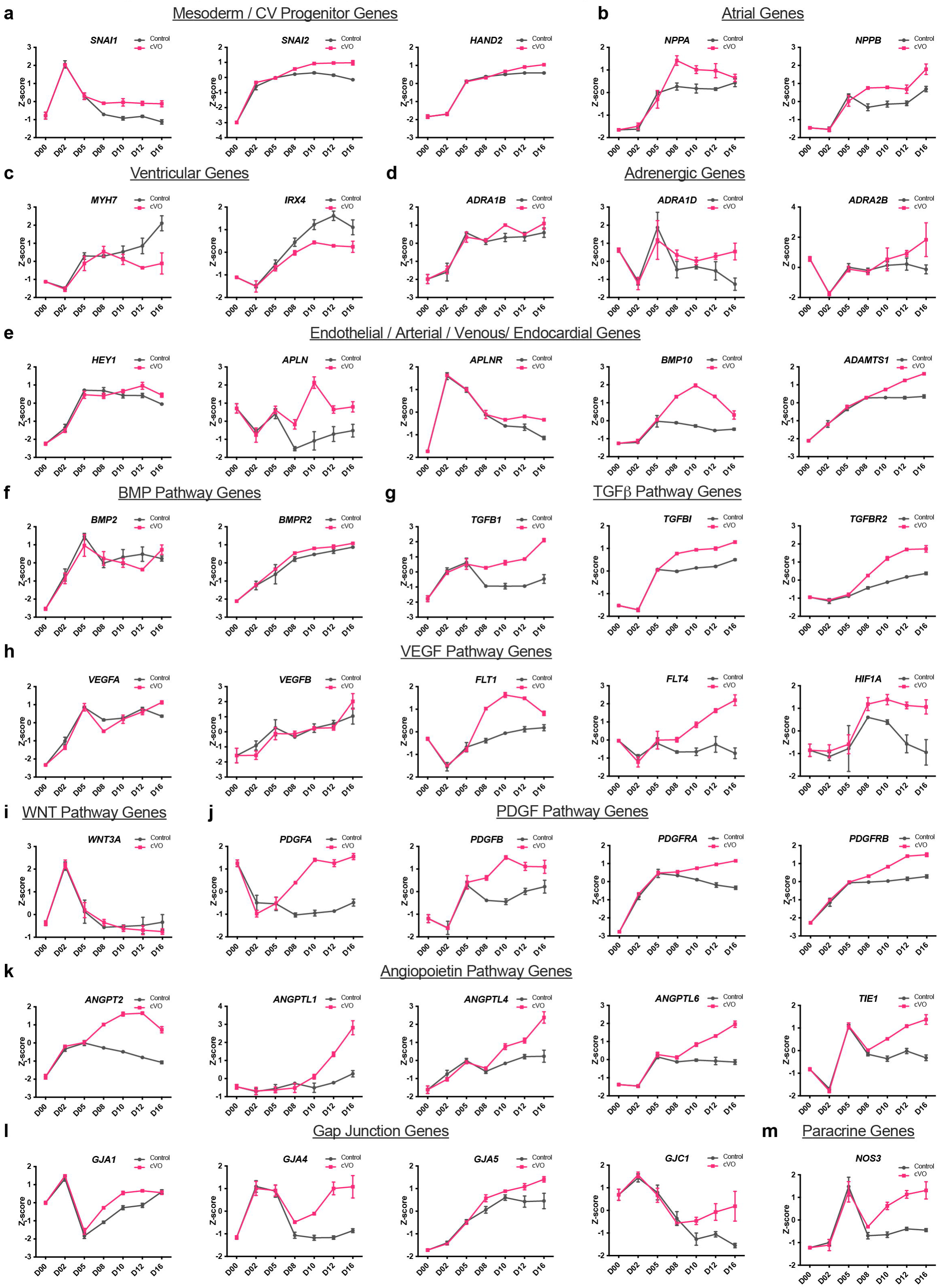
Temporal bulk RNA-seq expression for various cVO genes, related to Fig. 4. **a,** Mesoderm-precursor genes. **b,** Atrial genes. **c,** Ventricular genes. **d,** Adrenergic genes. **e,** Endothelial/arterial/venous/endocardial genes. **f,** BMP pathway genes. **g,** TGFβ pathway genes. **h,** VEGF pathway genes. **i,** WNT pathway genes. **j,** PDGF pathway genes. **k,** Angiopoietin pathway genes. **l,** Gap junction genes. **m,** Paracrine signaling genes.

**Extended Data Fig. 6.**
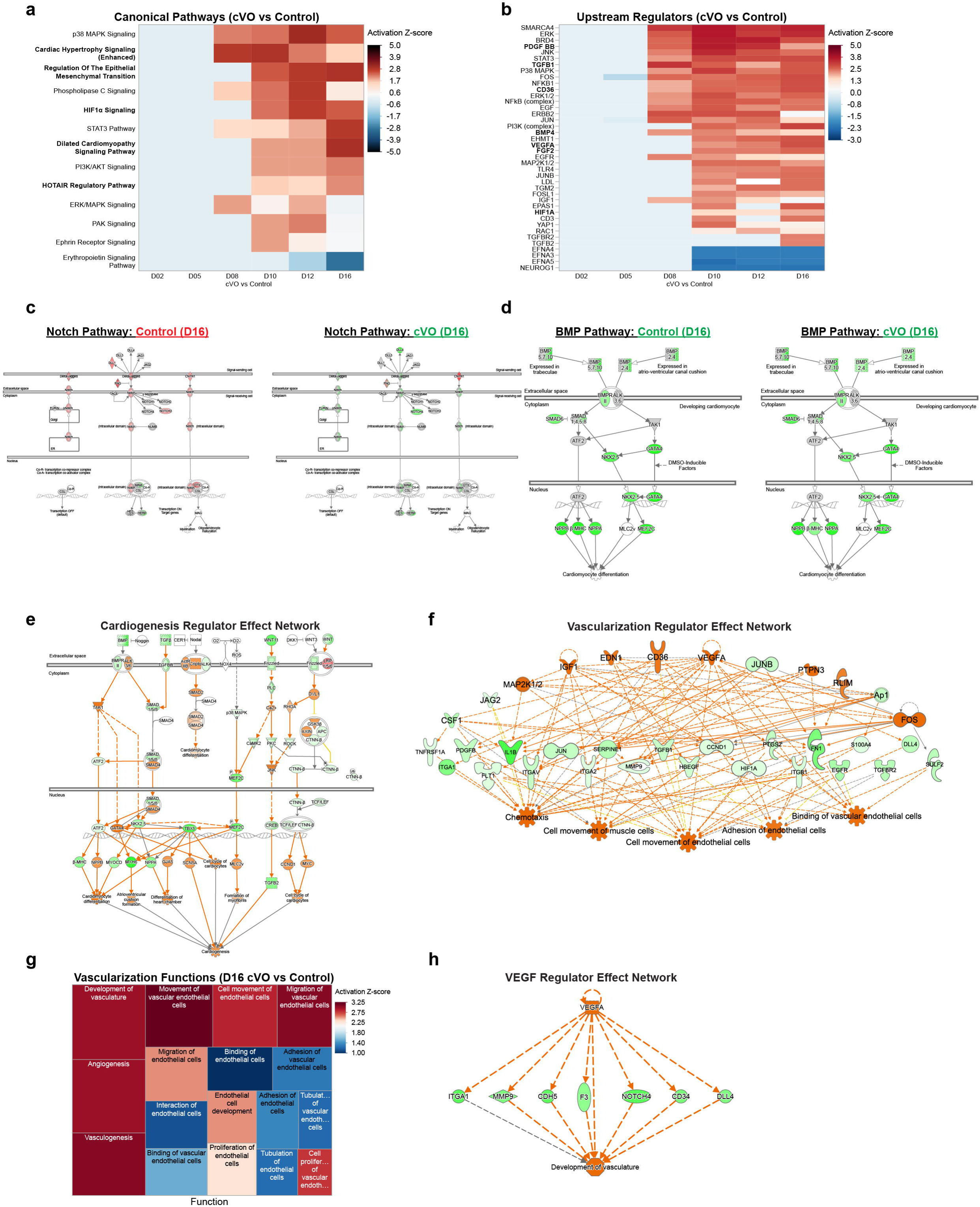
Ingenuity Pathway Analysis (IPA) of temporal bulk RNA-seq reveals cVO pathways, regulators, networks, and functions, related to Fig. 4. **a,** Days 2 to 16 cVO vs Control IPA showing a heat map of upregulated (red) and downregulated (blue) canonical pathways. **b,** Days 2 to 16 cVO vs Control IPA showing a heat map of upregulated (red) and downregulated (blue) upstream regulators (including *PDGFBB*, *TGFΒ1*, *CD36*, *BMP4*, *VEGFA*, *FGF2*, and *HIF1A*). **c,** IPA shows genes of the NOTCH pathway for the Control group at day 16 of differentiation (compared to day 0) are downregulated (red). In contrast, most genes of the NOTCH pathway for the cVO group at day 16 of differentiation (compared to day 0) are upregulated (green). **d,** IPA shows most genes are upregulated (green) in the BMP pathway for both the Control and cVO groups at day 16 of differentiation (compared to day 0). *BMP-2* is more upregulated in cVOs vs Controls. **e,** IPA of the WGCNA Dark Grey module within the “Cardiomyocyte/Endothelial/Smooth Muscle/Fibroblast Genes” cluster in Fig. 4h confirming a “Cardiogenesis Regulator Effect Network” with upstream regulators including *BMP*, *TGFβ*, and *WNT11* activating genes including *MEF2C*, *NKX2-5*, *TBX5*, *MYH6*, and *NPPA* leading to downstream cardiogenesis. Green indicates measured upregulated genes (|log2FC| > 2)). **f,** IPA of the WGCNA Pale Turquoise module within the “Vascularization Genes” cluster in Fig. 4h confirming a “Vascularization Regulator Effect Network” with upstream regulators including *VEGF*, *CD36*, *JAG2*, and *JUNB* activating genes including *PDGFB*, *FLT1*, *MMP9*, *TGFβ1*, *HIF1A*, *FN1*, and *DLL4*, leading to downstream vascularization functions. Green indicates measured upregulated genes (|log2FC| > 2). **g,** Day 16 cVO vs Control IPA showing a heat map of “Vascularization Functions”. Functions > Z-score of 2 are shaded red. Size of rectangles are proportional to -log (p-value). **h,** Day 16 cVO vs Control IPA showing “*VEGF* Regulator Effect Network” on *ITGA1*, *MMP9*, *CDH5*, *F3*, *NOTCH4*, *CD34*, and *DLL4* leading to downstream development of vasculature. Green indicates measured upregulated genes (|log2FC| > 2).

**Extended Data Fig. 7.**
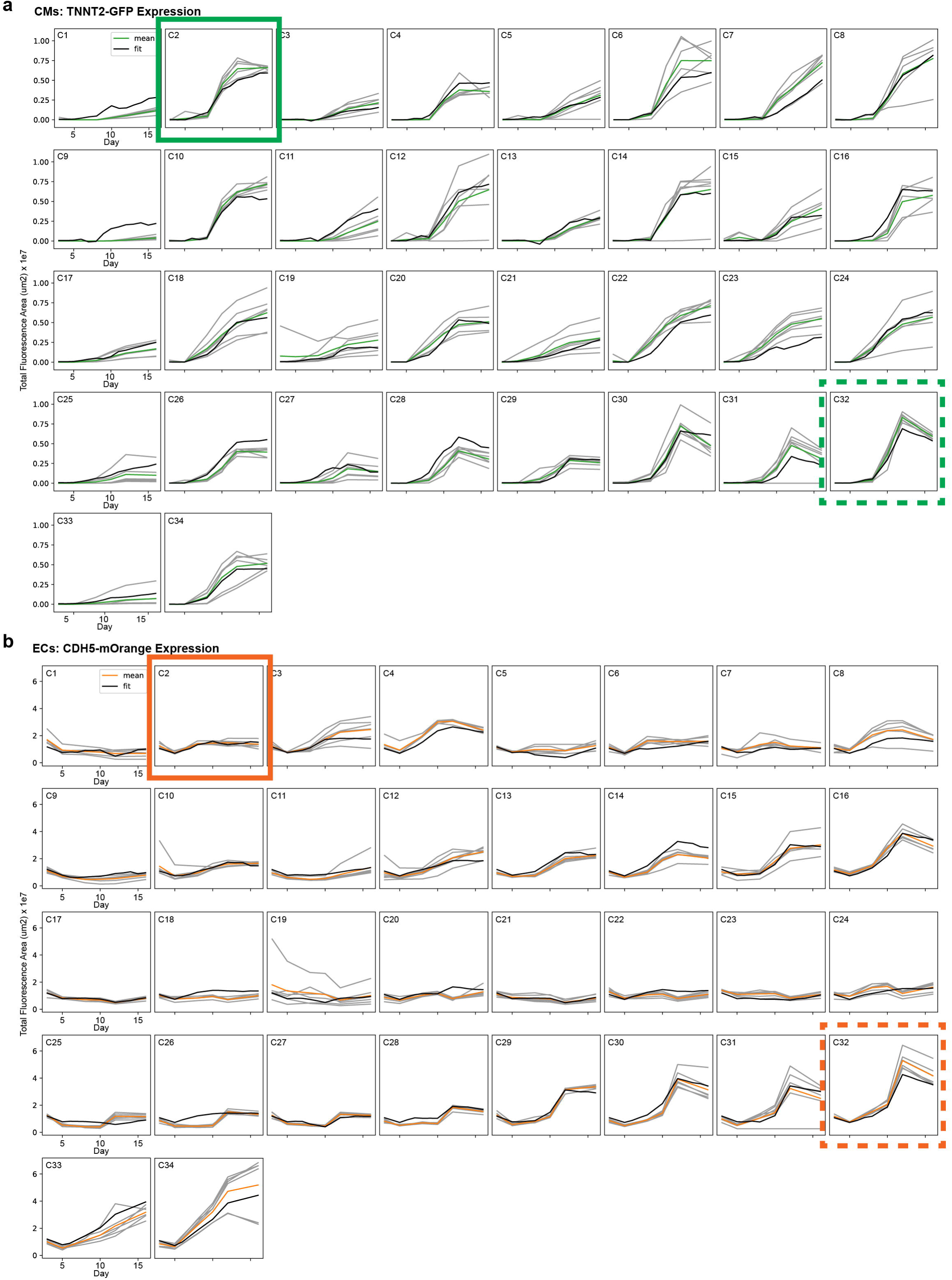
A machine learning multiple linear regression model created from screening conditions for cVO formation, related to Fig. 3. **a,** Multiple linear regression model created from 34 screening conditions (C1-C34) for cVO formation showing TNNT2-GFP total fluorescence area from days 2 to 16, indicating CM formation over time. Grey lines indicate 5-6 estimates from the original data, green lines indicate the mean of the estimates, and black lines indicate the fit of the model. The solid green square identifies C2 as the Control condition, and the dashed green square identifies C32 as the condition resulting in CMs within cVOs. **b,** Multiple linear regression model created from 34 screening conditions (C1-C34) for cVO formation showing CDH5-mOrange total fluorescence area from days 2 to 16, indicating EC formation over time. Grey lines indicate 5-6 estimates from the original data, green lines indicate the mean of the estimates, and black lines indicate the fit of the model. The solid orange square identifies C2 as the Control condition, and the dashed orange square identifies C32 as the condition resulting in ECs within cVOs.

**Extended Data Fig. 8.**
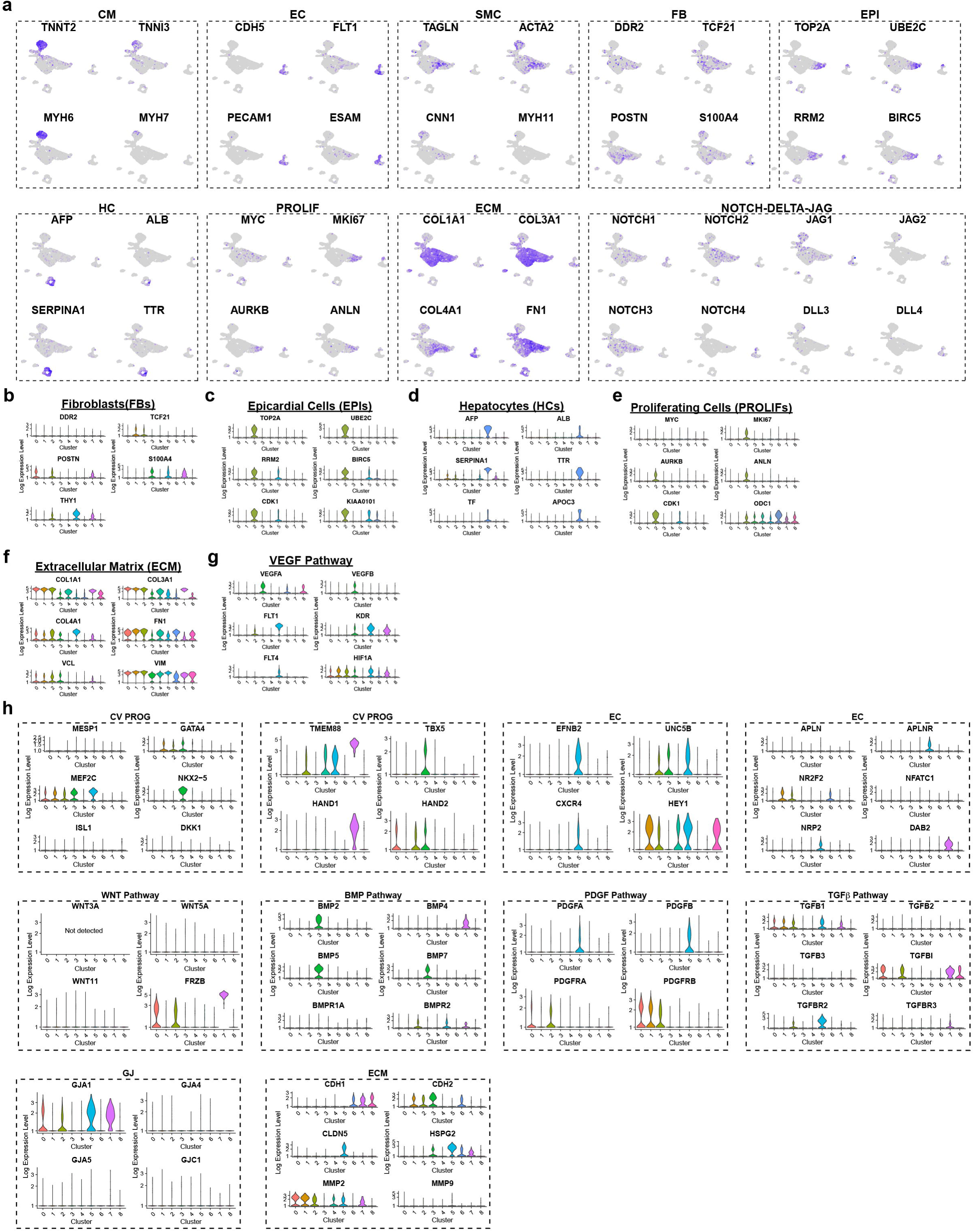
Single-cell RNA-seq reveals multiple vascular, endocardial, myocardial, and epicardial cell types in cVOs, related to Fig. 5. **a,** cVO violin plots showing FB (*DDR2, TCF21, POSTN, S100A4, THY1*). **b,** EPI (*TOP2A, UBE2C, RRM2, BIRC5, CDK1, KIAA0101*). **c,** HC (*AFP, ALB, SERPINA1, TTR, TF, APOC3*). **d,** PROLIFS (*MYC, MKI67, AURKB, ANLN, CDK1, ODC1*). **e,** ECM (*COL1A1, COL3A1, COL4A1, FN1, VCL, VIM*). **f,** VEGF Pathway (*VEGFA, VEGFB, FLT1, KDR, FLT4, HIF1A*). **g,** cVO UMAP feature plots showing CM (*TNNT2*, *TNNI3*, *MYH6*, *MYH7*), EC (*CDH5*, *FLT1*, *PECAM1*, *ESAM*), SMCs (*TAGLN*, *ACTA2*, *CNN1*, *MYH11*), FB (*DDR2*, *TCF21*, *POSTN*, *S100A4*), EPI (*TOP2A*, *UBE2C*, *RRM2*, *BIRC5*), HC (*AFP*, *ALB*, *SERPINA1*, *TTR*), PROLIF (*MYC*, *MKI67*, *AURKB*, *ANLN*), ECM (*COL1A1*, *COL3A1*, *COL4A1*, *FN1*), and NOTCH-DELTA-JAG pathway (*NOTCH1*, *NOTCH2*, *NOTCH3*, *NOTCH4*, *JAG1*, *JAG2*, *DLL3*, *DLL4*) gene expression. **h,** cVO violin plots showing CV PROG, EC, WNT pathway, BMP pathway, PDGF pathway, TGFβ pathway gap, junction (GJ), and ECM gene expression.

**Extended Data Fig. 9.**
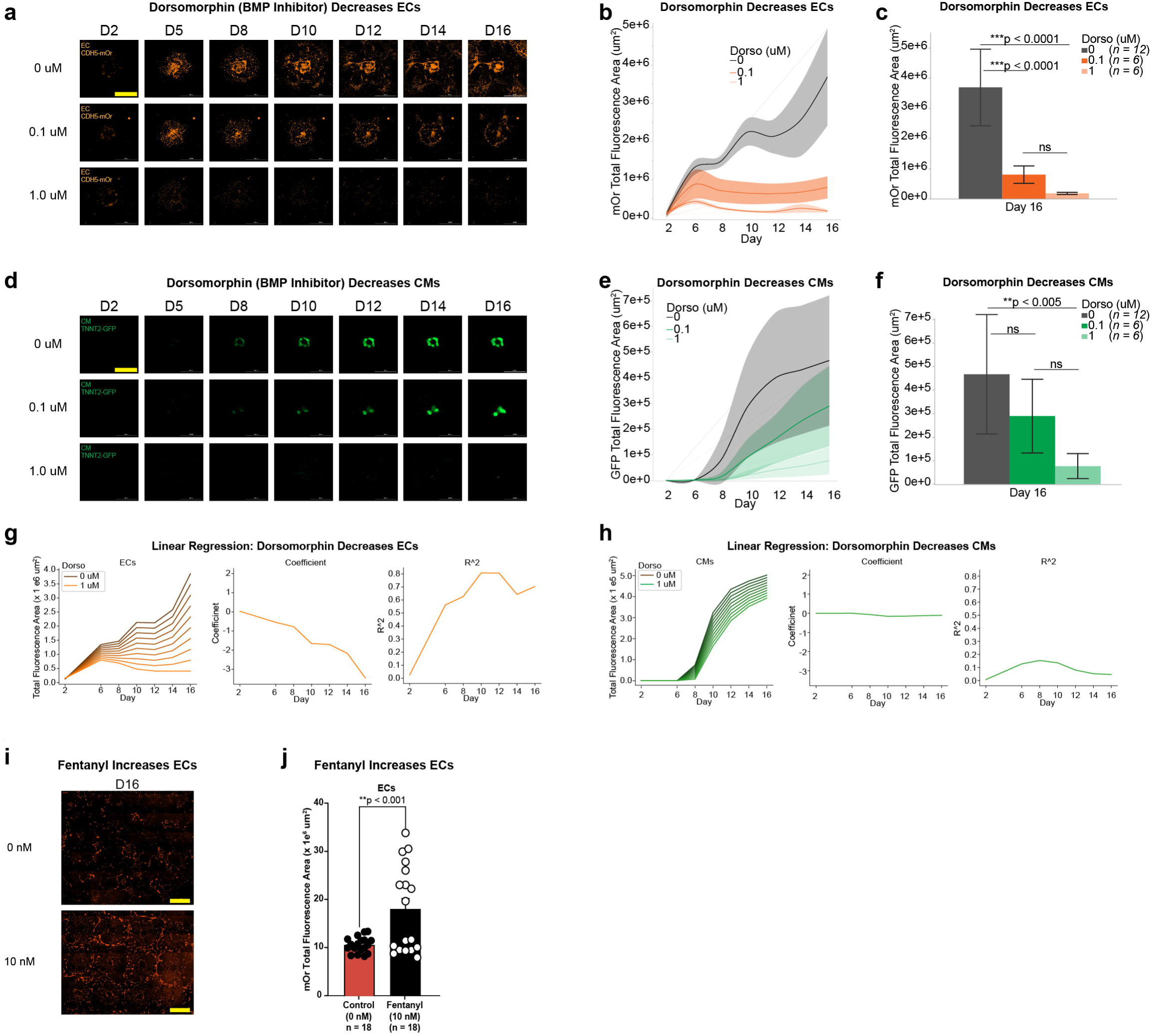
Inhibition of BMP pathway disrupts vasculature within cVOs, related to Fig. 6. **a-c,** Dorsomorphin, a BMP pathway antagonist, significantly decreased cVO EC formation over 16 days at 0.1 µM (n = 6, ***p < 0.0001) and 1.0 uM (n = 6, ***p < 0.0001) compared to 0 uM (Control) (n = 12). There was no significant (ns) difference between 0.1 and 1.0 µM. One-way ANOVA with Tukey’s test for multiple comparisons. Shaded bands and error bars ± 1 SD. Scale bar, 2 mm. Note, Dorsomorphin and DAPT (Fig. 6) were tested together and share the same Control. **d,** Multiple linear regression model effects of Dorsomorphin (between 0 and 1 uM) on cVO EC formation over 16 days (left) with linear regression coefficients (middle), and R^2^ (right). **e-g,** Dorsomorphin significantly decreased cVO CM formation over 16 days only at 1 µM (n = 6, **p < 0.005) compared to 0 µM (Control) (n = 12). There was no significant (ns) difference between the other two comparisons. One-way ANOVA with Tukey’s test for multiple comparisons. Shaded bands and error bars ± 1 SD. Scale bar, 2 mm. **h,** Multiple linear regression model effects of Dorsomorphin on cVO CM formation over 16 days (left) with linear regression coefficients (middle), and R^2^ (right). Based on the temporal graphs (**b, e, g, h**), the negative effect of Dorsomorphin on ECs was more than that on CMs; this was the opposite for DAPT. **i-j,** Fentanyl, a potent opioid agonist, significantly increased cVO EC formation over 16 days at 10 nM (n = 18, **p < 0.001) compared to 0 nM (Control) (n = 18). Student’s t-test. Error bars ± 1 SD. Scale bar, 2 mm.

**Extended Data Fig. 10.**
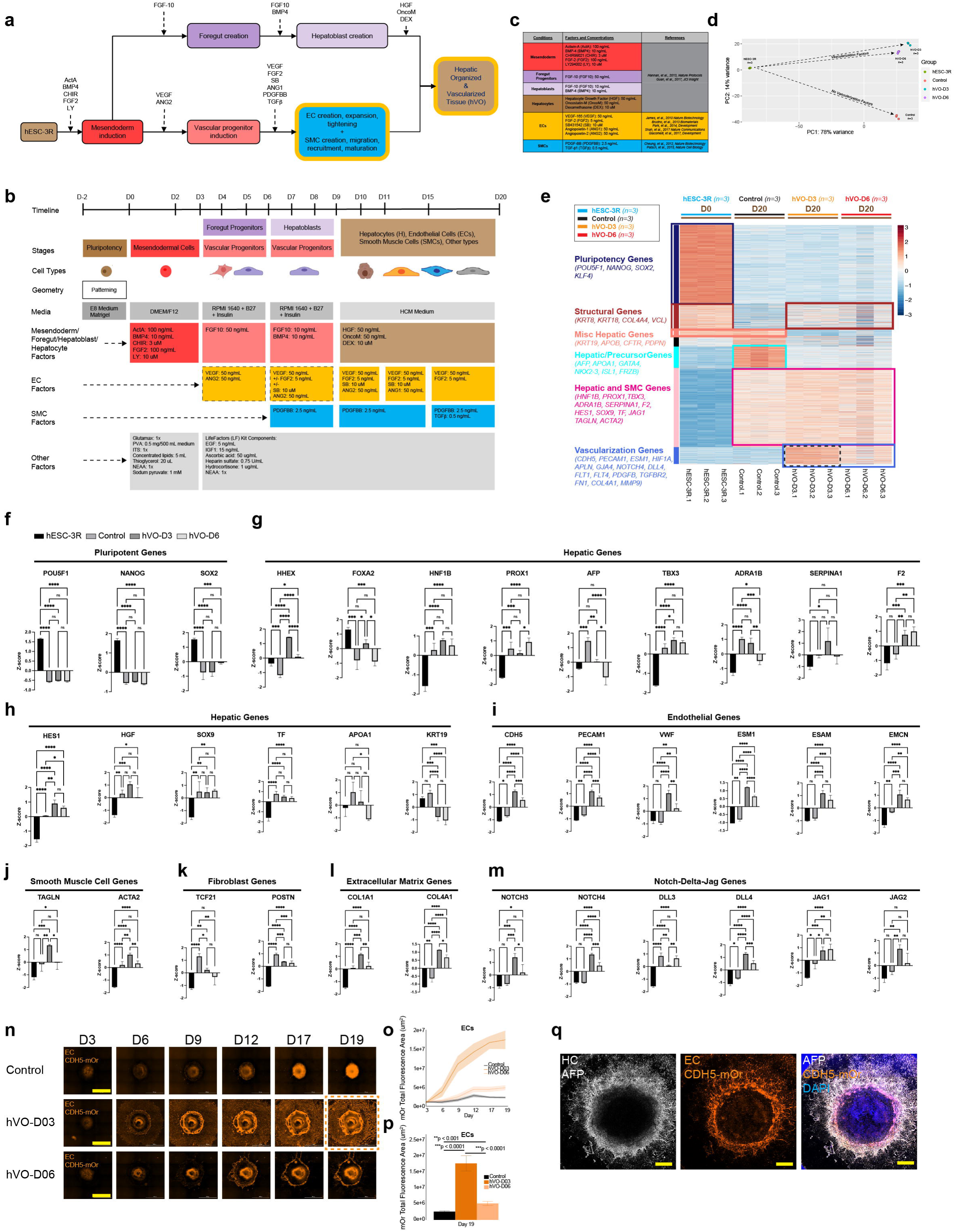
Vascularization factors used for creating cVOs enable creation of hVOs. **a,** Strategy for co-differentiating HCs, ECs, and SMCs to create hVOs. Key steps are inducing mesendoderm and then co-differentiating a vascular (CV) progenitor pool and hepatoblast pool that give rise to all three cell types. HC, hepatic cells. **b,** Schematic showing timeline, stage, cell types, geometry, media, and growth factors for creating HCs, ECs, SMCs, and resulting hVOs. **c,** Growth factor and small molecule characteristics for differentiation into HCs, ECs, and SMCs. Select references used as guidelines for concentrations and timing used in differentiation conditions are listed. Detailed conditions are listed in **Supplementary Table 2**. **d,** PCA of temporal bulk RNA-seq shows developmental differences of hESC-3R (day 0 undifferentiated hESC-3R micropatterns) (green, n = 3), Control (day 20 baseline hepatic differentiation, with no vascularization factors added) (red, n = 3), hVO-D3 (day 20 hVOs created by adding vascularization factors at day 3 of differentiation) (cyan, n = 3), compared to hVO-D6 (day 20 hVOs created by adding vascularization factors at day 6 of differentiation) (purple, n = 3). Note divergence between “Vascularization Factors” (D3 and D6) and “No Vascularization Factors” (Control) groups. **e,** bulk RNA-seq WGCNA heat map showing clusters of Pluripotency, Structural, Miscellaneous Hepatic, Hepatic/Precursor, Hepatic and Smooth Muscle Cell, and Vascularization Genes over twenty days (D0-20) for hESC-3R (blue, n = 3), Control (black, n = 3), hVO-D3 (orange, D3), and hVO-D6 (red, n = 3) groups. Black dashed rectangle shows vascularization genes are most upregulated in the hVO-D3 group. **f-m,** Comparison between hESC-3R, Control, hVO-D3, and hVO-D6 for select gene groups: Pluripotent, Hepatic, Endothelial, Smooth Muscle Cell, Fibroblast, Extracellular Matrix, and Notch-Delta-Jag. Overall, pluripotent genes are upregulated for hESC-3R and downregulated for Control, hVO-D3, and hVO-D6 groups. Several hepatic genes are upregulated for Control, hVO-D3, and hVO-D6 groups. Notably, all EC, SMC, ECM, and Notch-Delta-Jag (except *DLL3* and *JAG1*) genes are most upregulated for the hVO-D3 group. One-way ANOVA with Tukey’s test for multiple comparisons, *p < 0.05, **p < 0.01, ***p < 0.001, ****p < 0.0001, ns, not significant. **n-p,** Over 19 days, ECs increase most for the hVO-D3 group (n = 12) (dashed orange square in (N)) compared to the hVO-D6 (n = 10) and Control (n = 10) groups, indicating that the most ECs form when vascularization factors are added at day 3 to the baseline hepatic differentiation protocol. One-way ANOVA with Tukey’s test for multiple comparisons, **p < 0.001, ***p < 0.0001. Shaded bands and error bars +/- 1 SD. Scale bar, 2 mm. **q,** Individual confocal fluorescence images showing HCs (AFP, white) (left), ECs (CDH5-mOrange, orange) (middle), and merge (right). Note the endoderm-derived HC ring concentrically surrounding the mesoderm-derived EC ring. Nuclei are labeled with DAPI (blue). Scale bar, 2 mm.

