## Supplementary Material for "Micropatterned Organoids Enable Modeling of the Earliest Stages of Human Cardiac Vascularization"

### Supplementary Information

**Extended Discussion**

**Supplementary Videos 1-7**

**Supplementary Tables 1-4**

**Methods**

**References**

#### Extended Discussion

To capture the benefits of micropatterning, namely, to have a reproducible and scalable system that facilitates the identification of stereotypically-formed germ layers, progenitors, and derivative cell types from hPSCs, our system produces mostly 2D cardiac organoids. As such, our 2D organoids do not fully replicate the spatial relationships between endocardial, myocardial, and epicardial layers as recently shown in 3D cardiac organoids[1-5]. However, the 2D system allows for easier initial phenotypic screening with conventional microscopy, especially if combined with fluorescent reporter lines; for a 3D system, dedicated confocal or light sheet microscopy would be necessary to fully characterize entire organoid and vascularization volumetric characteristics such as vessel diameter distribution, branching orders, and growth over space and time.

We show proof-of-principle that our vascularization process could be applied to a 3D system and integration with the published 3D systems above could lead to recapitulating cardiogenesis with higher fidelity. Also, because our 2D cVOs contained "contaminating' hepatocytes, integration with the published 3D models above would minimize or eliminate endoderm-derived cell types. In addition, extending our 16 days of cVO differentiation to approximately 45 days would allow for closer comparison to the 6.5 PCW heart data and performing temporal single-cell transcriptomics would also give higher temporal resolution of vascular developmental processes. Despite these limitations, our prospective and directed differentiation of vascularized organoids represents a significant technical advance for exploring fundamental mechanisms of vascularization, which could be applied to other organ systems for understanding human development, disease modeling, drug toxicity testing, drug discovery, and regenerative medicine.

#### Supplementary Video Legends

**Supplementary Video 1.** Time-lapse of a 6 mm hESC-TNNT2-GFP micropattern over 9 days of differentiation.

**Supplementary Video 2.** Beating of a 2 mm hESC-NKX2-5-eGFP micropattern after 10 days of differentiation.

**Supplementary Video 3.** Time-lapse differentiation of single 2 mm hESC-3R micropatterns over 12 days showing formation of CMs (green), ECs (orange), and SMCs (blue) under 34 screening conditions. Condition 2 was Control and Condition 32 produced cVOs.

**Supplementary Video 4.** Time-lapse differentiation of a single cVO showing formation of CMs (green), ECs (orange), and SMCs (blue) over 12 days.

**Supplementary Video 5.** A cVO showing beating CMs (green) and SMCs (blue) intimately surrounded by branching ECs.

**Supplementary Video 6.** A 3D volume rendering showing cVO vasculature branching and lumen formation.

**Supplementary Video 7.** A cVO with ring-formed CMs showing beating in a rotating fashion and a different cVO showing calcium transients.

#### Supplementary Table Legends

**Supplementary Table 1. Cardiac Vascularized Organoid (cVO) Differentiation Screening Conditions**

**Supplementary Table 2. Hepatic Vascularized Organoid (hVOs) Differentiation Conditions**

**Supplementary Table 3. PCR Primers**

**Supplementary Table 4. Key Materials, Reagents, Software, and Equipment**

**Supplementary Table 1. Cardiac Vascularized Organoid (cVO) Differentiation Screening Conditions**

Plate 1

| Condition<br>Day | C1 | C2 | C3 | C4 | C5 | C6 | C7 | C8 |
|---|---|---|---|---|---|---|---|---|
| D0 | CHIR: 4<br>FGF2: 5 | CHIR: 5<br>FGF2: 5 | CHIR: 4<br>FGF2: 5 | CHIR: 5<br>FGF2: 5 | CHIR: 4<br>FGF2: 5 | CHIR: 5<br>FGF2: 5 | CHIR: 4<br>FGF2: 5 | CHIR: 5<br>FGF2: 5 |
| D1 | RB-I | RB-I | RB-I | RB-I | RB-I | RB-I | RB-I | RB-I |
| D3 | IWR: 5 | IWR: 5 | IWR: 5 | IWR: 5 | IWR: 5 | IWR: 5 | IWR: 5 | IWR: 5 |
| D5 | RB+I | RB+I | VEGF: 50 | VEGF: 50 | RB+I | RB+I | RB+I | RB+I |
| D7 | RB+I | RB+I | VEGF: 50 | VEGF: 50 | FGF2: 5 | FGF2: 5 | SB: 10 | SB: 10 |
| D9 | RB+I | RB+I | VEGF: 50 | VEGF: 50 | FGF2: 5 | FGF2: 5 | SB: 10 | SB: 10 |
| D11 | RB+I | RB+I | VEGF: 50 | VEGF: 50 | FGF2: 5 | FGF2: 5 | SB: 10 | SB: 10 |
| D13 | RB+I | RB+I | VEGF: 50 | VEGF: 50 | FGF2: 5 | FGF2: 5 | RB+I | RB+I |
| D16 | Stop | Stop | Stop | Stop | Stop | Stop | Stop | Stop |

Units CHIR, IWR, SB: uM; Units FGF2, VEGF: ng/mL
RB+/-I: RPMI-1640-B27 with/without insulin

Plate 2

| Condition<br>Day | C9 | C10 | C11 | C12 | C13 | C14 | C15 | C16 |
|---|---|---|---|---|---|---|---|---|
| D0 | CHIR: 4<br>FGF2: 5 | CHIR: 5<br>FGF2: 5 | CHIR: 4<br>FGF2: 5 | CHIR: 5<br>FGF2: 5 | CHIR: 4<br>FGF2: 5 | CHIR: 5<br>FGF2: 5 | CHIR: 4<br>FGF2: 5 | CHIR: 5<br>FGF2: 5 |
| D1 | RB-I | RB-I | RB-I | RB-I | RB-I | RB-I | RB-I | RB-I |
| D3 | IWR: 5 | IWR: 5 | IWR: 5 | IWR: 5 | IWR: 5 | IWR: 5 | IWR: 5 | IWR: 5 |
| D5 | ANG2: 50 | ANG2: 50 | RB+I | RB+I | VEGF: 50 | VEGF: 50 | VEGF: 50<br>ANG2: 50 | VEGF: 50<br>ANG2: 50 |
| D7 | ANG2: 50 | ANG2: 50 | FGF2: 5<br>SB: 10 | FGF2: 5<br>SB: 10 | VEGF: 50<br>FGF2: 5<br>SB: 10 | VEGF: 50<br>FGF2: 5<br>SB: 10 | VEGF: 50<br>FGF2: 5<br>SB: 10<br>ANG2: 50 | VEGF: 50<br>FGF2: 5<br>SB: 10<br>ANG2: 50 |
| D9 | ANG1: 50 | ANG1: 50 | FGF2: 5<br>SB: 10 | FGF2: 5<br>SB: 10 | VEGF: 50<br>FGF2: 5<br>SB: 10 | VEGF: 50<br>FGF2: 5<br>SB: 10 | VEGF: 50<br>FGF2: 5<br>SB: 10<br>ANG1: 50 | VEGF: 50<br>FGF2: 5<br>SB: 10<br>ANG1: 50 |
| D11 | ANG1: 50 | ANG1: 50 | FGF2: 5<br>SB: 10 | FGF2: 5<br>SB: 10 | VEGF: 50<br>FGF2: 5<br>SB: 10 | VEGF: 50<br>FGF2: 5<br>SB: 10 | VEGF: 50<br>FGF2: 5<br>SB: 10<br>ANG1: 50 | VEGF: 50<br>FGF2: 5<br>SB: 10<br>ANG1: 50 |
| D13 | RB+I | RB+I | FGF2: 5 | FGF2: 5 | VEGF: 50<br>FGF2: 5 | VEGF: 50<br>FGF2: 5 | VEGF: 50<br>FGF2: 5 | VEGF: 50<br>FGF2: 5 |
| D16 | Stop | Stop | Stop | Stop | Stop | Stop | Stop | Stop |

Units CHIR, IWR, SB: uM; Units FGF2, VEGF, ANG2, ANG1: ng/mL
RB+/-I: RPMI-1640-B27 with/without insulin

Plate 3

| Condition<br><br>Day | C17 | C18 | C19 | C20 | C21 | C22 | C23 | C24 |
|---|---|---|---|---|---|---|---|---|
| D0 | CHIR: 4<br><br>FGF2: 5 | CHIR: 5<br><br>FGF2: 5 | CHIR: 4<br><br>FGF2: 5 | CHIR: 5<br><br>FGF2: 5 | CHIR: 4<br><br>FGF2: 5 | CHIR: 5<br><br>FGF2: 5 | CHIR: 4<br><br>FGF2: 5 | CHIR: 5<br><br>FGF2: 5 |
| D1 | RB-I | RB-I | RB-I | RB-I | RB-I | RB-I | RB-I | RB-I |
| D3 | IWR: 5 | IWR: 5 | IWR: 5 | IWR: 5 | IWR: 5 | IWR: 5 | IWR: 5 | IWR: 5 |
| D5 | RB+I | RB+I | RB+I | RB+I | RB+I | RB+I | RB+I | RB+I |
| D7 | PDGFBB: 2.5 | PDGFBB: 2.5 | PDGFBB: 10 | PDGFBB: 10 | TGFβ: 0.5 | TGFβ: 0.5 | TGFβ: 2 | TGFβ: 2 |
| D9 | PDGFBB: 2.5 | PDGFBB: 2.5 | PDGFBB: 10 | PDGFBB: 10 | TGFβ: 0.5 | TGFβ: 0.5 | TGFβ: 2 | TGFβ: 2 |
| D11 | PDGFBB: 2.5 | PDGFBB: 2.5 | PDGFBB: 10 | PDGFBB: 10 | TGFβ: 0.5 | TGFβ: 0.5 | TGFβ: 2 | TGFβ: 2 |
| D13 | PDGFBB: 2.5<br><br>TGFβ: 0.5 | PDGFBB: 2.5<br><br>TGFβ: 0.5 | PDGFBB: 10<br><br>TGFβ: 2 | PDGFBB: 10<br><br>TGFβ: 2 | TGFβ: 0.5 | TGFβ: 0.5 | TGFβ: 2 | TGFβ: 2 |
| D16 | Stop | Stop | Stop | Stop | Stop | Stop | Stop | Stop |

Units CHIR, IWR: uM; Units FGF2, PDGFBB, TGFβ: ng/mL
RB+/-I: RPMI-1640-B27 with/without insulin

Plate 4

| Condition<br>Day | C25 | C26 | C27 | C28 | C29 | C30 | C31 | C32 |
|---|---|---|---|---|---|---|---|---|
| D0 | CHIR: 4<br>FGF2: 5 | CHIR: 5<br>FGF2: 5 | CHIR: 4<br>FGF2: 5 | CHIR: 5<br>FGF2: 5 | CHIR: 4<br>FGF2: 5 | CHIR: 5<br>FGF2: 5 | CHIR: 4<br>FGF2: 5 | CHIR: 5<br>FGF2: 5 |
| D1 | RB-I | RB-I | RB-I | RB-I | RB-I | RB-I | RB-I | RB-I |
| D3 | IWR: 5 | IWR: 5 | IWR: 5 | IWR: 5 | IWR: 5 | IWR: 5 | IWR: 5 | IWR: 5 |
| D5 | RB+I | RB+I | RB+I | RB+I | VEGF: 50<br>ANG2: 50 | VEGF: 50<br>ANG2: 50 | VEGF: 50<br>ANG2: 50 | VEGF: 50<br>ANG2: 50 |
| D7 | PDGFBB: 2.5<br>TGFβ: 0.5 | PDGFBB: 2.5<br>TGFβ: 0.5 | PDGFBB: 10<br>TGFβ: 2 | PDGFBB: 10<br>TGFβ: 2 | VEGF: 50<br>FGF2: 5<br>SB: 10<br>ANG2: 50<br>PDGFBB: 2.5 | VEGF: 50<br>FGF2: 5<br>SB: 10<br>ANG2: 50<br>PDGFBB: 2.5 | VEGF: 50<br>FGF2: 5<br>SB: 10<br>ANG2: 50<br>PDGFBB: 10 | VEGF: 50<br>FGF2: 5<br>SB: 10<br>ANG2: 50<br>PDGFBB: 10 |
| D9 | PDGFBB: 2.5<br>TGFβ: 0.5 | PDGFBB: 2.5<br>TGFβ: 0.5 | PDGFBB: 10<br>TGFβ: 2 | PDGFBB: 10<br>TGFβ: 2 | VEGF: 50<br>FGF2: 5<br>SB: 10<br>ANG1: 50<br>PDGFBB: 2.5 | VEGF: 50<br>FGF2: 5<br>SB: 10<br>ANG1: 50<br>PDGFBB: 2.5 | VEGF: 50<br>FGF2: 5<br>SB: 10<br>ANG1: 50<br>PDGFBB: 10 | VEGF: 50<br>FGF2: 5<br>SB: 10<br>ANG1: 50<br>PDGFBB: 10 |
| D11 | PDGFBB: 2.5<br>TGFβ: 0.5 | PDGFBB: 2.5<br>TGFβ: 0.5 | PDGFBB: 10<br>TGFβ: 2 | PDGFBB: 10<br>TGFβ: 2 | VEGF: 50<br>FGF2: 5<br>SB: 10<br>ANG1: 50<br>PDGFBB: 2.5 | VEGF: 50<br>FGF2: 5<br>SB: 10<br>ANG1: 50<br>PDGFBB: 2.5 | VEGF: 50<br>FGF2: 5<br>SB: 10<br>ANG1: 50<br>PDGFBB: 10 | VEGF: 50<br>FGF2: 5<br>SB: 10<br>ANG1: 50<br>PDGFBB: 10 |
| D13 | PDGFBB: 2.5<br>TGFβ: 0.5 | PDGFBB: 2.5<br>TGFβ: 0.5 | PDGFBB: 10<br>TGFβ: 2 | PDGFBB: 10<br>TGFβ: 2 | VEGF: 50<br>FGF2: 5<br>PDGFBB: 2.5<br>TGFβ: 0.5 | VEGF: 50<br>FGF2: 5<br>PDGFBB: 2.5<br>TGFβ: 0.5 | VEGF: 50<br>FGF2: 5<br>PDGFBB: 10<br>TGFβ: 2 | VEGF: 50<br>FGF2: 5<br>PDGFBB: 10<br>TGFβ: 2 |
| D16 | Stop | Stop | Stop | Stop | Stop | Stop | Stop | Stop |

Units CHIR, IWR, SB: uM; Units FGF2, VEGF, ANG2, ANG1, PDGFBB, TGFβ: ng/mL
RB+/-I: RPMI-1640-B27 with/without insulin

Plate 5

| Condition<br>Day | C33 | C34 |
|---|---|---|
| D0 | CHIR: 4<br>FGF2: 5 | CHIR: 5<br>FGF2: 5 |
| D1 | RB-I | RB-I |
| D3 | IWR: 5 | IWR: 5 |
| D5 | VEGF: 50<br>ANG2: 50<br>LF Kit | VEGF: 50<br>ANG2: 50<br>LF Kit |
| D7 | VEGF: 50<br>FGF2: 5<br>SB: 10<br>ANG2: 50<br>PDGFBB: 2.5<br>LF Kit | VEGF: 50<br>FGF2: 5<br>SB: 10<br>ANG2: 50<br>PDGFBB: 2.5<br>LF Kit |
| D9 | VEGF: 50<br>FGF2: 5<br>SB: 10<br>ANG1: 50<br>PDGFBB: 2.5<br>LF Kit | VEGF: 50<br>FGF2: 5<br>SB: 10<br>ANG1: 50<br>PDGFBB: 2.5<br>LF Kit |
| D11 | VEGF: 50<br>FGF2: 5<br>SB: 10<br>ANG1: 50<br>PDGFBB: 2.5<br>LF Kit | VEGF: 50<br>FGF2: 5<br>SB: 10<br>ANG1: 50<br>PDGFBB: 2.5<br>LF Kit |
| D13 | VEGF: 50<br>FGF2: 5<br>PDGFBB: 2.5<br>TGFβ: 0.5<br>LF Kit | VEGF: 50<br>FGF2: 5<br>PDGFBB: 2.5<br>TGFβ: 0.5<br>LF Kit |
| D16 | Stop | Stop |

Units CHIR, IWR, SB: uM; Units FGF2, VEGF, ANG2, ANG1, PDGFBB, TGFβ: ng/mL
RB+/-I: RPMI-1640-B27 with/without insulin
LifeFactors (LF) Kit: 5 ng/mL EGF, 15 ng/mL IGF-1, 50 ug/mL ascorbic acid, 0.75 U/mL heparin sulfate, 1 ug/mL hydrocortisone

##### Supplementary Table 2. Hepatic Vascularized Organoid (hVOs) Differentiation Conditions

Plate 1

| Condition | 1 (Control) | 2 (“hVO-D3”) | 3 (“hVO-D6”) |
|---|---|---|---|
| D0 | ActA: 100<br>BMP4: 10<br>CHIR: 3<br>FGF2: 100<br>LY: 10 | Same as Control | Same as Control |
| D3 | FGF10: 50<br><br>No vascular factors | Control +<br>VEGF: 50<br>ANG2: 50<br>LF Kit | Same as Control |
| D6 | FGF10: 50<br>BMP4: 10<br><br>No vascular factors | Control +<br>VEGF: 50<br>FGF2: 5<br>SB: 10<br>ANG2: 50<br>PDGFBB: 2.5<br>LF Kit | Control +<br>VEGF: 50<br>ANG2: 50<br>LF Kit |
| D9 | HGF: 50<br>OncoM: 50<br>DEX: 10<br><br>No vascular factors | Control +<br>VEGF: 50<br>FGF2: 5<br>SB: 10<br>ANG2: 50<br>PDGFBB: 2.5<br>LF Kit | Control +<br>VEGF: 50<br>FGF2: 5<br>SB: 10<br>ANG2: 50<br>PDGFBB: 2.5<br>LF Kit |
| D11 | HGF: 50<br>OncoM: 50<br>DEX: 10<br><br>No vascular factors | Control +<br>VEGF: 50<br>FGF2: 5<br>SB: 10<br>ANG1: 50<br>PDGFBB: 2.5<br>LF Kit | Control +<br>VEGF: 50<br>FGF2: 5<br>SB: 10<br>ANG1: 50<br>PDGFBB: 2.5<br>LF Kit |
| D13 | HGF: 50<br>OncoM: 50<br>DEX: 10<br><br>No vascular factors | Control +<br>VEGF: 50<br>FGF2: 5<br>SB: 10<br>ANG1: 50<br>PDGFBB: 2.5<br>LF Kit | Control +<br>VEGF: 50<br>FGF2: 5<br>SB: 10<br>ANG1: 50<br>PDGFBB: 2.5<br>LF Kit |
| D15, 17, 19 | HGF: 50<br>OncoM: 50<br>DEX: 10<br><br>No vascular factors | Control +<br>VEGF: 50<br>FGF2: 5<br>PDGFBB: 2.5<br>TGFβ: 0.5<br>LF Kit | Control +<br>VEGF: 50<br>FGF2: 5<br>PDGFBB: 2.5<br>TGFβ: 0.5<br>LF Kit 0.5 |
| D20 | Stop | Stop | Stop |

Units LY, CHIR, DEX, SB: uM
Units ActA, BMP4, FGF2, FGF10, HGF, OncoM, VEGF, ANG2, ANG1, PDGFBB, TGFβ: ng/mL
LifeFactors (LF) Kit: 5 ng/mL EGF, 15 ng/mL IGF-1, 50 ug/mL ascorbic acid, 0.75 U/mL heparin sulfate, 1 ug/mL hydrocortisone

#### Supplementary Table 3. PCR Primers

| Common Gene Name | NLM Gene Name | NLM Ascension Code | Forward Primer | Reverse Primer | Length (bp) | Ref |
|---|---|---|---|---|---|---|
| **PCR Primers for Lentiviral Plasmids Creation** | | | | | | |
| Adaptor-A/B | N/A | N/A | CTCTCTCGAGCGAGC GGCCGCTCTCTAGAT ATCCCGGGT | AATTACCCGGGATAT CTAGAGAGCGGCCGC TCGCTCGAGAGAGGG CC | N/A | This paper |
| SVPr-F/R | N/A | N/A | CTCTCTAGATATAGCA TTGAAAAAGGAAG | GTGAGTTTGGCCATA CCGGTCCGCCGAGAC AGCAA | N/A | This paper |
| Zeo-F/R | N/A | N/A | ACCGGTATGGCCAAA CTCACTTCTGCA | GTCCCCGGGACTAGT CCTGTTCCT | N/A | This paper |
| Bsd-F/R | N/A | N/A | CGGACCGGTATGCCC CTGAGCCAGGA | GCTCCCGGGTCAGCC CTCCCACACA | N/A | This paper |
| VmO-F/R | N/A | N/A | TCTCTCGAGATCGATG CTCATCCATGC | ATATCTAGACTTACTT GTACAGCTCGTCCAT G | N/A | This paper |
| TGN-F/R | N/A | N/A | AGCCTCGAGTCCGGA TCTTGGAA | GCGGTACCAAGCTTC TGACTGAGAGGGTGG GTTTCCCTAGCCAG | N/A | This paper |
| CFP-F/R | N/A | N/A | GTCAGAAGCTTGGTA CCGC | ATATCTAGATTAGCG GTACAGCTCGTCCAT G | N/A | This paper |
| **PCR Primers for Reporter Cell Line Creation** | | | | | | |
| GAPDH (GAPDH) | GAPDH | NM002046 | GGACTCATGACCACA GTCCATGCC | TCAGGGATGACCTTG CCCACAG | 152 | [6] |
| OCT-4 (POU5F1) | POU5F1 | BC117437 | CTTGCTGCAGAAGTG GGTGGAGGAA | CTGCAGTGTGGGTTTC GGGCA | 169 | [6] |
| Green Fluorescent Protein (GFP) | GFP | - | ACGTAAACGGCCACA AGTTC | AAGTCGTGCTGCTTCA TGTG | 187 | [6] |
| CDH5 (VE-Cadherin) Promoter-mOrange (CDH5) | CDH5 | NM001795.5 | CCAACGGAACAGAAA CATCC | AATCCAGAGGTTGAT TGTCG | 1672 | This paper |
| mOrange (mOrange) | mOrange | - | AGGGCTTTCAGACCG CTAAG | TCCAACTTGATGCCG ACGAT | 484 | This paper |

| | | | | | | |
|---|---|---|---|---|---|---|
| Zeocin-1 (Zeocin1) | ZEO | - | GGGCTGTAGAGTTCT GGACTGA | CTGCTGAGATGAAGA GGGTGA | 105 | This paper |
| Zeocin-2 (Zeocin2) | ZEO | - | CCGGTATGGCCAAAC TCACT | TCTAGGCCTCTGACCC AGAC | 220 | This paper |
| Cyan Fluorescent Protein (CFP) | CFP | - | TCCAGTGCTTCGCCCG CTAC | TTCGCCTCCAGGCCGT TGTT | 289 | This paper |
| Blasticidin (Blasticidin) | BSD | - | TAGCCGCAAACATAG TTCAATACA | TGTTCCCAGCACCACC AGTT | 234 | This paper |

NLM: National Library of Medicine

#### Supplementary Table 4. Key Materials, Reagents, Software, and Equipment

| Materials, Reagents, Software, and Equipment | Source | Identifier |
|---|---|---|
| Antibodies | | |
| Mouse Anti-Human Oct-3/4 mAb (Clone 40/Oct-3) | BD Biosciences | 611202; RRID:AB_398736 |
| Goat Anti-Human Nanog pAb | R & D Systems | AF1997; RRID:AB_355097 |
| Rabbit Anti-Human Sox2 XP® mAb (Clone D6D9) | Cell Signaling Technology | 3579S; RRID:AB_2195767 |
| Mouse Anti-Human CDX2 mAb (Clone CDX2-88) | Abcam | ab157524; RRID:AB_2721036 |
| Goat Anti-Human Brachyury pAb | R & D Systems | AF2085; RRID:AB_2200235 |
| Mouse Anti-Human Sox17 mAb (Clone # 245013) | R & D Systems | mab1924; RRID:AB_2195646 |
| Rabbit Anti-Human Cardiac Troponin-T pAb | Abcam | ab45932; RRID:AB_956386 |
| Mouse Anti-Human Alpha-Actinin mAb (Clone EA-53) | Sigma-Aldrich | A7811; RRID:AB_476766 |
| Rabbit Anti-Human VE-Cadherin mAb (Clone D87F2) | Cell Signaling Technology | 2500S; RRID:AB_10839118 |
| Mouse Anti-Human CD31 mAb (Clone JC70A) | Thermo Fisher Scientific | MA5-13188; RRID:AB_10982120 |
| Rabbit Anti-Human vWF pAb | Agilent Technologies | A0082; RRID:AB_2315602 |
| Rabbit Anti-Human TAGLN/SM22-alpha pAb | Abcam | ab14106; RRID:AB_443021 |
| Mouse Anti-Human Calponin mAb (Clone hCP) | Sigma-Aldrich | C2687; RRID:AB_476840 |
| Mouse Anti-Human SMA mAb (Clone 1A4) | Sigma-Aldrich | A2547; RRID: AB_476701 |
| Mouse Anti-Human Vinculin mAb (Clone 7F9) | Santa Cruz Biotechnology | sc-73614; RRID:AB_1131294 |
| Goat Anti-GFP Polyclonal Ab | Abcam | ab6673; RRID:AB_305643 |
| Rabbit Anti-Human Alpha-1-Fetoprotein pAb | Agilent Technologies | A0008; RRID:AB_2650473 |

| | | |
|---|---|---|
| Donkey Anti-Goat IgG Alexa Fluor 488 | Thermo Fisher Scientific | A-11055; RRID:AB_2534102 |
| Donkey Anti-Rabbit IgG Alexa Fluor 488 | Thermo Fisher Scientific | A-11055; RRID:AB_2535792 |
| Donkey Anti-Mouse IgG Alexa Fluor 488 | Thermo Fisher Scientific | A-21202; RRID:AB_141607 |
| Donkey Anti-Goat IgG Alexa Fluor 546 | Thermo Fisher Scientific | A-11056; RRID:AB_142628 |
| Donkey Anti-Goat IgG Alexa Fluor 555 | Thermo Fisher Scientific | A-21432; RRID:AB_141788 |
| Donkey Anti-Rabbit IgG Alexa Fluor 594 | Thermo Fisher Scientific | A-21207; RRID:AB_141637 |
| Goat Anti-Mouse IgG Alex Fluor 594 | Thermo Fisher Scientific | A-11005; RRID:AB_2534073 |
| Donkey Anti-Rabbit IgG Alexa Fluor 647 | Thermo Fisher Scientific | A-31573; RRID:AB_2536183 |
| Goat Anti-Rabbit IgG Alexa Fluor 647 | Thermo Fisher Scientific | A-21245; RRID:AB_2535813 |
| Donkey Anti-Mouse IgG Alexa Fluor 647 | Thermo Fisher Scientific | A-31571; RRID:AB_162542 |
| Donkey Anti-Goat IgG Alexa Fluor 647 | Thermo Fisher Scientific | A-21447; RRID:AB_2535864 |
| CD144 (VE-Cadherin) MicroBeads, human | Miltenyi Biotec | 130-097-857 |
| Bacterial and Virus Strains | | |
| pLV-hVPr-mOrange-SVPr-Zeo | This paper | N/A |
| pLV-hTAGLNPr-CFP-SVPr-Bsd | This paper | N/A |
| Chemicals, Peptides, and Recombinant Proteins | | |
| Y-27632 (dihydrochloride) | MedChem Express | HY-10583 |
| CHIR-99021 | Selleck Chemicals | S2924 |
| IWR-1 | Selleck Chemicals | S7086 |
| SB431542 | Selleck Chemicals | S1067 |
| FGF-2 | PeproTech | 100-18B |
| VEGF-165 | PeproTech | 100-20 |
| Angiopoietin-1 (ANG-1) | PeproTech | 130-06 |
| Angiopoietin-2 (ANG-2) | PeproTech | 130-07 |
| PDGF-BB | PeproTech | 100-14B |
| TGF-β1 | PeproTech | 100-21C |
| EGF | PeproTech | AF-100-15 |

| | | |
|---|---|---|
| LDN193189 | Selleck Chemicals | S2618 |
| Retinoic acid (RA) | Sigma-Aldrich | R2625 |
| Activin-A | R & D Systems | 338-AC |
| BMP-4 | R & D Systems | 314-BP |
| LY294002 | Cayman Chemical | 70920 |
| FGF-10 | PeproTech | 100-26 |
| HGF | PeproTech | 100-39H |
| Dexamethasone | Tocris | 1126 |
| Oncostatin M | PeproTech | 300-10 |
| DAPT (GSI-IX) | Selleck Chemicals | S2215 |
| Dorsomorphin | Sigma-Aldrich | P5499 |
| Zeocin | Thermo Fisher Scientific | R25001 |
| Blasticidin | Thermo Fisher Scientific | A1113903 |
| IC Fixation Buffer | Thermo Fisher Scientific | FB001 |
| DRAQ5™ Fluorescent Probe Solution (5 mM) | Thermo Fisher Scientific | 62251 |
| 4′,6-diamidino-2-phenylindole (DAPI) | Sigma-Aldrich | D8417 |
| Critical Commercial Assays | | |
| Fluo-4 Direct Calcium Assay Kit, Starter pack | Thermo Fisher Scientific | F10471 |
| FluoVolt™ Membrane Potential Kit | Thermo Fisher Scientific | F10488 |
| Nitric Oxide (total), Detection Kit | Enzo | ADI-917-020 |
| FluoSpheres™ Carboxylate-Modified Microspheres | Thermo Fisher Scientific | F8807 |
| Deposited Data | | |
| Bulk RNA-seq data of hESC, Control, cVO, and hVO samples | This paper | GEO: GSE185194 |
| Single-cell RNA-seq data of Control and cVO samples | This paper | GEO: GSE185194 |
| Experimental Models: Cell Lines | | |
| Human H9 hESC | WiCell | WA09 |
| Human H9 hESC-NKX2-5-eGFP | Stanley Lab (Elliott, et al., 2011) | N/A |
| Human H9 hESC-TNNT2-GFP | WiCell | H9-hTnnT2-pGZ-TD2 |
| Human H9 hESC-3R | This paper | N/A |
| Human hESC-RUES2-GLR | Brivanlou Lab (Martyn et al., 2018) | N/A |
| Human 113 hiPSC | Stanford CVI Biobank | 113 |
| HEK 293FT | Thermo Fisher Scientific | R70007 |
| Oligonucleotides | | |
| See Supplementary Table 3. PCR Primers | This paper | N/A |
| Recombinant DNA | | |
| pLV-hVPr-mOrange | Rafii Lab (James, et al. 2011) | N/A |
| pSicoR | Addgene | 11579 |

| | | |
|---|---|---|
| pGL3 Luciferase Reporter | Promega | E1741 |
| pGreenZeo Human Nanog Reporter | System Biosciences | SR10030VA-1 |
| pCMV/Bsd (BsdCassette™ Vector) | Thermo Fisher Scientific | V51020 |
| pEZX-TAGLN-GLuc | GeneCopoeia | HPRM24364-PG02 |
| pTagCFP-N | Evrogen | FP112 |
| psPAX2 | Addgene | 12260 |
| pMD2.G | Addgene | 12259 |
| Software and Algorithms | | |
| SnapGene Viewer v5.0.6 | SnapGene | https://www.snapgene.com |
| Gen5 v3.05 | BioTek | https://www.biotek.com |
| Zeiss ZEN (Blue Edition) v2.6 | Zeiss | https://www.zeiss.com/microscopy/int/home.html |
| LAS X | Leica | https://www.leica-microsystems.com |
| Imaris v9.6.1 | Oxford Instruments | https://imaris.oxinst.com |
| ImageJ (Fiji) v2.1.0 | NIH | https://imagej.nih.gov |
| Micromanager v1.4 | Vale Lab (Edelstein, et al., 2010) | https://micro-manager.org |
| BV-Ana v1604 | SciMedia | https://www.scimedia.com |
| SI8000R Analyzer | Sony Corporation | https://www.sonybiotechnology.com/us |
| DESeq2 | Bioconductor | https://bioconductor.org/packages/release/bioc/html/DESeq2.html |
| Ingenuity Pathway Analysis (IPA) v65367011 | Qiagen | https://www.qiagen.com |
| GraphPad Prism v9 | GraphPad | https://www.graphpad.com |
| JMP Pro 15 | SAS Institute | https://www.jmp.com/en_us/home.html |
| R v3.6.1 and 4.0.4 | R Core | https://cran.r-project.org/bin/macosx/ |
| RStudio v1.4.1106 | RStudio | https://www.rstudio.com/products/rstudio/download/ |
| Cell Ranger v3.0.1 | 10X Genomics | https://www.10xgenomics.com/software |
| Loupe Cell Browser v3.0.1 | 10X Genomics | https://www.10xgenomics.com/software |
| Seurat v3.1.5 and 4.0.0 | Satija Lab (Stuart, et al., 2019) | https://satijalab.org/seurat/ |
| Microsoft Excel 2016 | Microsoft | https://www.microsoft.com/en-us/ |

| | | |
|---|---|---|
| Adobe Illustrator 2021 | Adobe | https://www.adobe.com |
| Adobe Photoshop 2021 | Adobe | https://www.adobe.com |
| Biorender | Biorender | https://biorender.com |
| Other | | |
| Single-cell RNA sequencing of 6.5 PCW human heart | Lundeberg Lab (Asp et al., 2019) | EGAS00001003996 |
| Library QC and Bulk RNA-Sequencing (20M Raw Reads/Sample) | Illumina | N/A |
| Library QC and Single-Cell RNA-Sequencing (HiSeq PE150) | Illumina | N/A |
| Chromium Single-Cell 3' Library & Gel Bead Kit v2 | 10X Genomics | PN-120267 |
| Normal Goat Serum (10%) | Thermo Fisher Scientific | 50062Z |
| Alconox | Midland Scientific | Cat#ALC1104 |
| Pluronic® F-127, 0.2 µm filtered (10% Solution in Water) | Thermo Fisher Scientific | P6866 |
| 16% Paraformaldehyde | Electron Microscopy Sciences | 15713S |
| Ethanol, Absolute (200 Proof), Molecular Biology Grade | Fisher Scientific | BP2818500 |
| Triton X-100 | Sigma-Aldrich | X100 |
| Tween-20 | Sigma-Aldrich | P5927 |
| 40 um cell strainer, Corning 352340 | Fisher Scientific | 08-771-1 |
| 70 um cell strainers, Corning 352350 | Fisher Scientific | 08-771-2 |
| 6-well plates, Falcon 353046 | Fisher Scientific | 08-772-1B |
| 24-well plates, Falcon 353047 | Fisher Scientific | 08-772-1 |
| 48-well plates, Falcon 353078 | Fisher Scientific | 08-772-1C |
| 96-well plates-Greiner CELLSTAR® black wells 655090 | Sigma-Aldrich | M0562-32EA |
| 24-well plates-Softwell Easy Coat custom 16 kPa hydrogels | Matrigen | SW24-EC-C |
| Essential 8 (E8) | Thermo Fisher Scientific | A1517001 |
| CGMP MTESR 1 | Stemcell Technologies | 85850 |
| StemMACS™ iPS-Brew XF (Brew) | Miltenyi Biotec | 130-104-368 |
| RPMI 1640 | Thermo Fisher Scientific | 11875093 |
| RPMI 1640 Medium, no phenol red | Thermo Fisher Scientific | 11835030 |
| DMEM, high glucose, no glutamine | Thermo Fisher Scientific | 11960044 |
| Ham's F-12 Nutrient Mix | Thermo Fisher Scientific | 11765054 |
| DMEM/F-12, GlutaMAX™ supplement | Thermo Fisher Scientific | 10565042 |
| Medium 231 | Thermo Fisher Scientific | M231500 |
| Medium 199, Earle's Salts | Thermo Fisher Scientific | 11150059 |
| Leibovitz's L-15 medium, no phenol red | Thermo Fisher Scientific | 21083027 |
| KnockOut Serum Replacement | Thermo Fisher Scientific | 10828028 |
| IMDM | Thermo Fisher Scientific | 12440053 |
| B-27™ Supplement (50X), serum free | Thermo Fisher Scientific | 17504044 |

| | | |
|---|---|---|
| B-27™ Supplement, minus insulin | Thermo Fisher Scientific | A1895601 |
| GlutaMAX™ Supplement | Thermo Fisher Scientific | 35050061 |
| Smooth Muscle Growth Supplement (SMGS) | Thermo Fisher Scientific | S00725 |
| MEM Non-Essential Amino Acids (NEAA) Solution (100X) | Thermo Fisher Scientific | 11140050 |
| Poly(vinyl alcohol) (PVA) | Sigma-Aldrich | P8136 |
| Insulin | Sigma-Aldrich | I9278 |
| Transferrin | Thermo Fisher Scientific | 11107018 |
| Insulin-Transferrin-Selenium (ITS) | Thermo Fisher Scientific | 41400045 |
| Chemically Defined Lipid Concentrate | Thermo Fisher Scientific | 11905031 |
| 1-thyoglycerol >97% | Sigma-Aldrich | M6145 |
| L-Ascorbic acid BioXtra, ≥99.0%, crystalline | Sigma-Aldrich | A5960-25G |
| Penicillin-Streptomycin (10,000 U/mL) | Thermo Fisher Scientific | 15140122 |
| HBSS, calcium, magnesium, no phenol red | Thermo Fisher Scientific | 14025092 |
| Tyrode's Salt Solution | Sigma-Aldrich | T2397 |
| Live Cell Imaging Solution | Thermo Fisher Scientific | A14291DJ |
| DPBS, calcium, magnesium | Thermo Fisher Scientific | 14040133 |
| VascuLife® VEGF LifeFactors® Kit | Lifeline Cell Technology | LS-1020 |
| EGM-2 BulletKit | Lonza | CC-3162 |
| HCM Hepatocyte Culture Medium BulletKit | Lonza | CC-3198 |
| Corning Matrigel Basement Membrane Matrix, LDEV-free | Fisher Scientific | 354234 |
| Fibronectin from bovine plasma | Sigma Aldrich | F1141-2MG |
| Gelatin solution | Sigma Aldrich | G1393 |
| EDTA (0.5 mM) | Thermo Fisher Scientific | 15575-020 |
| Pierce™ Primary Cardiomyocyte Isolation Kit | Thermo Fisher Scientific | 88281 |
| TrypLE Select Enzyme (1X), w/o phenol red | Thermo Fisher Scientific | 12563011 |
| Lipofectamine 2000 | Thermo Fisher Scientific | 11668019 |
| PEG-it Virus Precipitation Solution | System Biosciences | LV810A |
| PureLink Genomic DNA Mini Kit | Thermo Fisher Scientific | K182000 |
| AccuPrime SuperMix I | Thermo Fisher Scientific | 12342 |
| 10-1500 bp DNA QuantLadder | Lonza | 50475 |
| 1.2 % FlashGel DNA Cassette | Lonza | 57029 |
| Direct-zol RNA Microprep w/ Zymo-Spin IC Columns (Capped) | Zymo Research | R2060 |
| RNaseZap® RNase Decontamination Solution | Thermo Fisher Scientific | AM9782 |
| Silhouette Cameo | Silhouette America | B007R83VKE |
| Silhouette Cameo Cutting Mat-12"X12" | Silhouette America | CUT-MAT-12-3T |
| Press-to-Seal™ Silicone Sheet | Thermo Fisher Scientific | P18178 |
| BD-20AC Laboratory Corona Treater | Electro-Technic Products | 12051A |
| MiniMACS™ separator | Miltenyi Biotec | 130-090-312 |

#### Methods

All experiments, methods, and protocols for this study were approved by the Stanford University Stem Cell Research Oversight (SCRO) committee. Key materials, reagents, software, and equipment are listed in **Supplementary Table 4** and will be provided upon reasonable request.

##### Cell Lines

We used the following hPSC (both hESC and hiPSC) lines. The human H9 WT hESC (WA09, WiCell) was previously used as the parental line to create the human H9 hESC-NKX2-5-eGFP line[7] and the human H9 hESC-TNNT2-GFP line (H9-hTnnT2-pGZ-TD2, WiCell[8]. In our study here, we used the hESC-TNNT2-GFP line to create the human hESC-triple reporter (3R) line (described in more detail in the Reporter Cell Line Creation section below). To fluorescently visualize mesoderm, endoderm, and ectoderm germ layers, we used the human hESC-RUES2 GLR line generously provided by the Brivanlou Lab[9]). Finally, we used the human 113 WT hiPSC line (113, Stanford CVI Biobank) to validate our hESC cVO results in a hiPSC line.

##### Relevant Biological Variables

The hESC-3R and hESC-RUES2-GLR lines are female cell lines (ethnicity unknown) while the 113 hiPSC line is a male and White (not Hispanic and Latino) cell line. In future studies, we will investigate the potential impact of gender and ethnicity on 3D cardiovascular development.

##### Construction of Lentiviral Plasmids

The two final lentiviral vector plasmids pLV-CDH5Pr-mOrange-SVPr-Zeo and pLV-TAGLNPr-CFP-SVPr-Bsd were constructed as described below. To facilitate selection and identification of cell lines transduced with lentivirus created with these vectors, we designed a series of double-cassette lentiviral vectors that feature cell-specific fluorescent reporter cassettes and constitutively expressed drug-selection cassettes. **Extended Data Fig. 3a-d** illustrates the overall design of these vectors.

First, a blank lentivirus vector pLV-B was derived from pSicoR (11579, Addgene). The plasmid pSicoR was digested by ApaI-EcoRI, and re-ligated with an oligo adaptor (Adaptor-A/B) containing restriction sites XhoI, NotI, XbaI, and XmaI. The oligo adaptor and PCR primers used in this study are listed in **Supplementary Table 3**.

Next, a cassette with the SV40 promoter (SVPr) controlling the expression of Zeocin (Zeo) was introduced into the blank lentivirus vector pLV-B to create the plasmid pLV-SVPr-Zeo. The template for the SVPr comes from the pGL3 luciferase reporter plasmid (E1741, Promega), and Zeo from the pGreenZeo human Nanog reporter plasmid (SR10030VA-1, System Biosciences). The cassette SVPr-Zeo was assembled by PCR with primer pairs SVPr-F/R and Zeo-F/R as previously described[10]. The derived fragment was restricted with XbaI-XmaI and ligated into pLV-B at the same sites to create pLV-SVPr-Zeo.

Then, the Blasticidin (Bsd) coding sequence (CDS) was introduced to replace the Zeocin CDS in pLV-SVPr-Zeo. The Bsd CDS was amplified from the pCMV/Bsd (BsdCassette™ Vector) plasmid (V51020, Thermo Fisher) with primer pair Bsd-F/R. The amplified fragment was digested with AgeI-XmaI and ligated into pLV-SVPr-Zeo at the same sites. Thus, the plasmid pLV-SVPr-Bsd was created.

The plasmid pLV-CDH5Pr-mOrange-SVPr-Zeo was constructed as follows. The cassette carrying the CDH5 (VE-Cadherin) promoter (CDH5Pr) controlling expression of the mOrange fluorescent protein was amplified from the plasmid pLV-CDH5Pr-mOrange[11], which is a gift from the Rafii Lab, Cornell, with primer pair VmO-F/R. The amplified fragment was restricted with XhoI-XbaI and ligated into pLV-SVPr-Zeo at the same sites to create pLV-CDH5Pr-mOrange-SVPr-Zeo.

The plasmid pLV-TAGLNPr-CFP-SVPr-Bsd was constructed by inserting a human transgelin (TAGLN) (SM22-alpha) promoter-directed CFP cassette (TAGLNPr-CFP) into pLV-SVPr-Bsd. This TAGLNPr-CFP cassette was obtained by PCR assembly of the TAGLNPr and the CFP CDS with primer pairs TGN-F/R and CFP-F/R as previously described[10]. The TAGLN promoter sequence comes from the pEZX-TAGLN-GLuc plasmid (HPRM24364-PG02GeneCopoeia), and the CFP CDS from the pTagCFP-N plasmid (FP112, Evrogen). The derived fragment was restricted with XhoI-XbaI and ligated into pLV-SVPr-Bsd at the same sites to create pLV-TAGLNPr-CFP-SVPr-Bsd. Verification of plasmid construction was done by Sanger sequencing.

**Lentivirus Production**

To package plasmid in lentivirus, HEK 293FT cells (R70007, Thermo Fisher) were transfected separately with target plasmids (pLV-CDH5Pr-mOrange-SVPr-Zeo or pLV-TAGLNPr-CFP-SVPr-Bsd, and the packaging plasmids psPAX2 (containing GAG and POL) (12260, Addgene) and pMD2.G (containing VSV-G) (12259, Addgene)) using Lipofectamine 2000 (11668019, Thermo Fisher). Transfected HEK293FT cells were incubated for 3 days and media collected daily. Media was then centrifuged at 3000 g for 15 min to remove cells and cell debris. The supernatant was concentrated using PEG-it Virus Precipitation Solution (LV810A, System Biosciences) according to the manufacturer's protocol.

**Reporter Cell Line Creation**

The human H9 hESC-TNNT2-GFP cell line[8] (H9-hTnnT2-pGZ-TD2, WiCell) was seeded on 6-well plates and first transduced with the lentivirus expressing CDH5Pr-mOrange-SVPr-Zeo. Cells were selected by Zeocin (0.5-3 μg/mL) treatment for 1-2 weeks. Genomic DNA was isolated from several clones, PCR was performed, and gel electrophoresis was used to verify expression of CDH5 (VE-Cadherin), mOrange, and Zeocin. This double reporter cell line was then transduced with the lentivirus expressing TAGLNPr-CFP-SVPr-Bsd to create the human H9 hESC-triple reporter (3R) cell line (hESC-3R). Cells were selected by Blasticidin (3 μg/mL) treatment for 1-2 weeks. Again, genomic DNA was isolated from several clones, PCR was performed, and gel electrophoresis was used to verify the presence of TAGLN (SM22-alpha), CFP, and Blasticidin (**Extended Data Fig. 3e-g**). PCR primers are listed in **Supplementary Table 3**.

**Non-quantitative PCR**

Non-quantitative PCR was performed using custom-designed PCR primers (**Supplementary Table 3**). Genomic DNA was isolated with the PureLink Genomic DNA Mini Kit (K182000, Thermo Fisher) and quantified using a Nanodrop 2000 Spectrophotometer (ND-2000, Thermo Fisher) per the manufacturer's instructions. For PCR amplification, AccuPrime SuperMix I (12342, Thermo Fisher), custom primers for pluripotency and lentiviral markers (**Supplementary Table 3**), and genomic DNA were combined. Non-template control (NT) reactions were prepared by substituting DNA with distilled water. Samples were transferred to a thermal cycler and the following cycling program was used: a) initial denaturation at 94 °C for 2 min; b) 30 cycles of 94 °C, 30 sec; 60 °C, 30 sec; 68 °C, 1 min; c) final extension at 68 °C for 5 min. Reactions were maintained at 4 °C after cycling and then stored at -20 °C. The PCR products and a 10-1500 bp DNA QuantLadder (50475, Lonza) were loaded in a 1.2% 16+1 double-tier FlashGel DNA Cassette (57029, Lonza), run, and visualized with the FlashGel System (57067, Lonza).

**Human Pluripotent Stem Cell (hPSC) Culture**

Human pluripotent stem cells (hPSCs) (both hESCs and hiPSCs) were maintained in the pluripotent state through daily feeding with Essential 8 (E8) medium (A1517001, Thermo Fisher Scientific). The medium was changed daily, and cells were passaged every 3-4 days with 0.5 mM EDTA (15575-020, Thermo Fisher Scientific) between a 1:6 and 1:12 split ratio. hPSCs were passaged on to tissue culture plates coated with Corning Matrigel membrane matrix (354234, Thermo Fisher Scientific) at a dilution of 1:100 for either maintenance or differentiation. To aid in passaging undifferentiated hPSCs by minimizing anoikis (dissociation-induced apoptosis), we cultured cells for 24 hours with 10 uM of the ROCK inhibitor Y-27632 (HY-10583, MedChem Express). Excess cells were frozen in 90% Knockout Serum Replacement (KOSR) (#10828010, Thermo Fisher Scientific) with 10% DMSO (#D2438, Sigma-Aldrich) in cryovials and frozen at -80 °C overnight and then subsequently transferred to liquid nitrogen storage.

**Pluripotency Markers**

Using standard protocols, undifferentiated hPSCs were labeled with primary antibodies for the pluripotency markers Oct-3/4 (mouse anti-human Oct-3/4 mAb (Clone 40), 1:400, 611202, BD Biosciences), Nanog (goat anti-human Nanog pAb, 1:100, AF1997, R&D Systems), and Sox2 (rabbit anti-human Sox2 mAb (Clone D6D9), 1:200, 3579S, Cell Signaling Technology). Secondary antibodies used were donkey anti-mouse IgG Alexa 488 for Oct-3/4, 1:500 (A21202,

Thermo Fisher Scientific); donkey anti-goat IgG Alexa 555 for Nanog, 1:500 (A21432, Thermo Fisher Scientific); and donkey anti-rabbit IgG Alexa 647 for Sox2, 1:500 (A31573, Thermo Fisher Scientific). Cell nuclei were counterstained with 4′,6-diamidino-2-phenylindole (DAPI) (#D8417, Sigma-Aldrich, 1 mg/mL + 0.1% Triton X-100 in PBS) or DRAQ5™ Fluorescent Probe Solution (5 mM) (62251, Thermo Fisher Scientific). Primary and secondary antibodies are listed in **Supplementary Table 4**.

**Stencil Creation for hPSC Colony Micropatterning**

Our previous micropatterning method[12] used a high-cost laser cutter that is not available in many labs. Here, to create stencils for hPSC colony micropatterning, we used a low-cost, widely available, table-top Silhouette Cameo cutting tool (B007R83VKE, Silhouette America, Amazon) to design and then cut circular patterns from a press-to-seal silicone sheet (P18178, Thermo Fisher Scientific). To fit in one well of a standard Falcon 48-well plate (353078, Thermo Fisher Scientific), the outer diameter of the circular patterns was cut to approximately 9.5 mm. To fit in one well of a standard Falcon 6-well plate (353046, Thermo Fisher Scientific), the outer diameter of the circular patterns was cut to approximately 34 mm. At the center of the 48-well pattern, a single 2 mm hole was cut to create a complete single-hole stencil; for the 6-well circular pattern, an evenly spaced array (3 x 3 to 7 x 7 with a pitch of 2 to 5 mm from center to center) of 2 mm holes were cut to create a complete multi-hole stencil array. The stencils were then washed 3 times with deionized (DI) water for 20 minutes. For previously used stencils, cleaning was performed 3 times for 30 minutes with Alconox anionic detergent (ALC1104, Midland Scientific) in an ultrasonic cleaning bath (Branson Ultrasonic Bath, Emerson Electric) to remove any residual cellular debris. After detergent cleaning, stencils were washed again 6 times with DI water for 15 minutes in the ultrasonic bath. Next, to render the silicone stencil surfaces hydrophilic, a handheld laboratory corona treater (12051A, Electro-Technic Products) was swept back and forth across the stencils approximately 2 times for 5 minutes. The stencils were then autoclaved at 15 psi at 121° C for 30 minutes. Finally, the stencils were kept submerged in 100% ethanol until they were ready to be placed in the multi-well plates.

**Single and Multiple hPSC Colony Micropatterning**

Stencils for either a 48-well plate or a 6-well plate were placed in each well and pressed onto the well surface with sterile forceps. Next, ice cold Corning Matrigel membrane matrix (354234, Thermo Fisher Scientific) at a dilution of 1:100 in 0.25 mL ice cold RPMI-1640 (11875-093, Thermo Fisher Scientific) was poured into each well containing a stencil and was gelled by incubating at 37° C for at least 1 hour. Undifferentiated hPSCs were dissociated with 0.5 mM EDTA (15575-020 Gibco/ Thermo Fisher Scientific) and passaged as a single cell suspension that included 10 μM of the ROCK inhibitor Y-27632 (HY-10583, MedChem Express) to prevent dissociation-induced apoptosis. The cells at a concentration of 1.0 – 1.5 x $10^6$ cells/mL were introduced into the Matrigel- and stencil-containing wells and were allowed to attach at 37° C overnight in 0.25 mL Essential 8 (E8) medium. The next day, stencils were removed with sterile forceps, leaving micropatterned cells attached on the 2 mm circular islands created by the 2 mm holes of the stencils. At this concentration, the 2 mm colonies were essentially 100 % confluent in 2-3 days, showing good border integrity and uniform cell density. When colonies were filled in, the cells were fed with ice cold E8 which contained Matrigel at a dilution of 1:100 and immediately placed back in the incubator to minimize cold shock. This was required to allow the colonies to migrate beyond their 2 mm initial boundaries upon differentiation. The differentiation protocol was initiated when the colonies were essentially 100 % confluent, typically 2-3 days after seeding. We observed successful differentiation on circular micropatterns at diameters of 2, 4, and 6 mm, but chose 2 mm because this size gave more repeatable results and allowed greater outward migration within the 48-well plates.

**Growth Factors and Small Molecules**

To induce mesoderm and subsequent cardiomyocyte differentiation, we used 4, 5, or 6 uM CHIR-99021 (CHIR) (S2924, Selleck Chemicals) and 5 uM IWR-1 (IWR) (S7086, Selleck Chemicals) and 5 ng/mL FGF-2 (FGF2) (100-18B, PeproTech). These two small molecules and one growth factor served as the basis for our baseline (Control group) differentiation method (described in more detail in the Baseline Cardiomyocyte Differentiation section).

To simultaneously induce endothelial cell vasculogenesis and angiogenesis along with cardiomyocyte differentiation (described in more detail in the Cardiovascular Differentiation section), we added 50 ng/mL VEGF (VEGF) (100-20, PeproTech); 5 ng/mL FGF-2 (FGF2) (100-18B, PeproTech); 10 uM SB431542 (SB) (S1067, Selleck Chemicals); 50 ng/mL Angiopoietin-2 (ANG2) (130-07, PeproTech); 50 ng/mL Angiopoietin-1 (ANG1) (130-06, PeproTech); and select components of the VascuLife® VEGF LifeFactors® Kit (LS-1020, Lifeline Cell Technology) (5 ng/mL EGF, 15 ng/mL IGF-1, 50 ug/mL ascorbic acid, 0.75 U/mL heparin sulfate, 1 ug/mL hydrocortisone, 5 ng/mL VEGF (not added), 5 ng/mL FGF-2 (not added), 2 % FBS (not added), 30 mg/mL/15 ug/mL gentamicin/amphotericin B (not added), 10 mM glutamine (not added)).

To simultaneously induce smooth muscle cell differentiation with cardiomyocyte and endothelial cell differentiation (described in more detail in the Cardiovascular Differentiation section), we added 2.5 or 10 ng/mL PDGF-BB (PDGFBB) (100-14B, PeproTech) and 0.5 or 2 ng/mL TGF-β1 (TGFβ) (100-21C, PeproTech).

Finally, to maintain isolated endothelial cells, we used the EGM-2 Bullet Kit (CC-3162, Lonza) (EGF, IGF-1, ascorbic acid, heparin sulfate, hydrocortisone, VEGF, FGF-2, FBS, and gentamicin/ amphotericin-B (manufacturer does not provide concentrations)).

To induce mesendoderm and subsequent hepatocyte differentiation (described in more detail in the Hepatovascular Differentiation section), we used 100 ng/mL Activin-A (338-AC, R & D Systems); 10 ng/mL BMP-4 (314-BP, R & D Systems); 3 uM CHIR-99021 (S2924, Selleck Chemicals); 10 uM LY294002 (PI3K-AKT inhibitor) (70920, Cayman Chemical Company); 50 ng/mL FGF-10 (100-26, PeproTech); 50 ng/mL HGF (100-39H, PeproTech); 50 ng/mL Oncostatin-M (300-10, PeproTech); 10 uM Dexamethasone (1126/100, Tocris); and HCM Hepatocyte Culture Medium Bullet Kit (CC-3198, Lonza).

**Baseline Cardiomyocyte (CM) Differentiation**

For baseline cardiomyocyte (CM) differentiation (Control group), hPSCs were transferred to Matrigel-coated non-micropatterned and micropatterned surfaces as described above and differentiated in RPMI-1640 media (61870, Thermo Fisher Scientific) supplemented with B27 without insulin (A1895601, Thermo Fisher Scientific). Our baseline differentiation method was based on a previously described small molecule-based monolayer method[13, 14]. Briefly, on the first day (Day 0) of differentiation, basal medium was supplemented with 4, 5, or 6 uM CHIR99021 (CHIR) (S2924, Selleck Chemicals) and 5 ng/mL FGF-2 (FGF2) (100-18B, PeproTech). On Day 2, the medium was replaced with basal medium containing B27 without insulin. On Day 3, the medium was replaced with basal medium containing B27 without insulin and containing 5 uM

IWR-1 (IWR) (S7086, Selleck Chemicals). On Day 5 the medium was replaced with basal medium containing B27 without insulin. From Days 7-16, the basal medium was changed to contain B27 with insulin (17504044, Thermo Fisher Scientific) and was replaced every 48 hours thereafter. Cardiomyocytes generally began spontaneously beating sometime between Days 7 and 10. hPSC-CMs were collected on Day 16 for further downstream studies. More complex cardiac vascularized organoid (cVO) differentiation described below was based on this baseline differentiation and contained several additional small molecules and growth factors to promote simultaneous cardiac and vascular co-differentiation.

**Endothelial Cell (EC) Differentiation**

We differentiated ECs from hPSCs as previously described[15]. Briefly, hPSCs were seeded on Matrigel-coated plates in E8 medium (A1517001, Thermo Fisher Scientific) to 50% confluency. At 80% confluency, hPSCs were differentiated towards the mesodermal lineage by treatment of 6 μM CHIR-99021 (S2924, Selleck Chemicals) in RPMI-1640 media (61870, Thermo Fisher Scientific) supplemented with B27 without insulin (A1895601, Thermo Fisher Scientific) for 2 days, followed by a second treatment of 2 μM CHIR-99021 in RPMI-B27 without insulin for 2 days. At day 4 of differentiation, the cells were subjected to a differentiation medium comprised of RPMI-B27 without insulin and the EGM-2 Bullet Kit (CC-3162, Lonza) supplemented with 20 ng/mL BMP-4 (314-BP, R & D Systems), 50 ng/mL VEGF (100-20, PeproTech), and 20 ng/mL FGF-2 (100-18B, PeproTech). Cells were subjected to medium change every 2 days, where the concentration of RPMI-B27 without insulin was gradually decreased and replaced by EGM-2 by day 10. On day 12, the differentiating hPSC-ECs were isolated using Magnetic-activated cell sorting (MACS), where cells were dispersed by TrypLE (12563011, Thermo Fisher Scientific), incubated with bead-conjugated anti-VE-Cadherin antibody (130-097-857, Miltenyi Biotec), and sorted with a MiniMACS™ separator (130-090-312, Miltenyi Biotec). Sorted VE-Cadherin+ cells were collected and seeded on 0.2% gelatin (G1393, Sigma-Aldrich)-coated plates and maintained in EGM-2 medium supplemented with 10 μM SB431542 (S1067, Selleck Chemicals), a TGF-β inhibitor.

**Smooth Muscle Cell (SMC) Differentiation**

We prepared basal, induction, and maintenance media and differentiated SMCs as previously described[16]. Briefly, basal chemically defined medium (CDM) consisted of 50% vol/vol IMDM (12440053, Thermo Fisher Scientific), 50% vol/vol Ham's F-12 Nutrient Mix (11765054, Thermo Fisher Scientific), 1% vol/vol chemically defined lipid concentrate (11905031, Thermo Fisher Scientific); 2 mM Glutamax supplement (35050061, Thermo Fisher Scientific), 1 mg/mL PVA (P8136, Sigma-Aldrich), 7 μg/mL insulin (I9278, Sigma-Aldrich), 15 μg/mL transferrin (11107018, Thermo Fisher Scientific), and 450 μM 1-thioglycerol (M6145, Sigma-Aldrich). Neural crest (NC) induction medium consisted of CDM, 1 μM CHIR99021 (S2924, Selleck Chemicals), 2 μM SB431542 (S1067, Selleck Chemicals), 250 nM LDN193189 (S2618, Selleck Chemicals), 15 ng/mL BMP4 (314-BP, R & D Systems). Neural crest (NC) maintenance medium consisted of CDM, 10 ng/mL FGF2 (100-18B, PeproTech), 10 ng/mL EGF (AF-100-15, PeproTech), and 2 μM SB431542 (S1067, Selleck Chemicals). SMC induction medium consisted of CDM, 2 ng/mL TGF-β1 (100-21C, PeproTech), and 10 ng/mL PDGF-BB (100-14B, PeproTech).

Cardiac neural crest (CNC) cell differentiation was performed as follows. On Day 0, ~80% confluent hPSCs were dissociated and seeded at a density of $2\times10^4$ cells/cm$^2$ onto new Matrigel-coated 6-well plates in StemMACS™ iPS-Brew XF medium ("Brew") (130-104-368, Miltenyi Biotec) + 10 uM Y-27632 (HY-10583, MedChem Express). On Days 1-4, NC induction medium was added and changed daily. On Days 5-6, NC induction medium + 1 uM retinoic acid (R2625, Sigma-Aldrich) was added daily. On Day 7, cells were dissociated and split 1:6 in NC maintenance medium + 1 uM retinoic acid + 10 uM Y-27632. On Days 8-9, NC maintenance medium + 1 uM retinoic acid was refreshed daily. On Day 10, cells were dissociated and split 1:6 on gelatin (G1393, Sigma-Aldrich)-coated 6-well plates at a density of $2 \times 10^4$ cells/$cm^2$. NC maintenance medium was refreshed every day and cells were passaged when they reached ~90% confluency. From Days 11-21, SMC induction medium was refreshed daily. On Days 22-28, cells were maintained in Medium 231 (M231500, Thermo Fisher Scientific) + Smooth Muscle Growth Supplement (SMGS) (S00725, Thermo Fisher Scientific). Media which was refreshed every two days and cells were split 1:2 when they became confluent.

**Cardiac Vascularized Organoid (cVO) Differentiation**

See **Fig. 3c**, **Extended Data Fig. 3h-i**, and **Supplementary Table 1** for an overview of our thirty-four (34) cardiovascular (CV) differentiation screening conditions to produce cVOs. The Control groups utilized only CHIR, FGF2, and IWR. To begin, hPSCs suspended in single cells were transferred to Matrigel-coated non-micropatterned or micropatterned surfaces as described above and differentiated in RPMI-1640 media (61870, Thermo Fisher Scientific) supplemented with B27 without insulin (A1895601, Thermo Fisher Scientific) (RB-I). The CV differentiation was based on the baseline differentiation described previously[13, 14] to promote CM differentiation and contained several additional small molecules and growth factors to promote simultaneous endothelial and smooth muscle cell co-differentiation. On Day 0, for primarily mesoderm induction[17], basal medium was supplemented with 4 or 5 uM CHIR99021 (CHIR) (S2924, Selleck Chemicals) and 5 ng/mL FGF-2 (FGF2) (100-18B, PeproTech). On Day 3, for CM induction, 5 uM IWR-1 (IWR) (S7086, Selleck Chemicals) was added.

On Days 5-16, to simultaneously induce endothelial cell vasculogenesis and angiogenesis[18-21] along with the cardiomyocyte differentiation, we added combinations of 50 ng/mL VEGF-165 (VEGF) (Days 5, 7, 9, 11, 13) (100-20, PeproTech); 5 ng/mL FGF-2 (FGF2) (Days 7, 9, 11, 13) (100-18B, PeproTech); 10 uM SB431542 (SB) (Days 7, 9, or 11) (S1067, Selleck Chemicals); 50 ng/mL Angiopoietin-2 (ANG2) (Days 5 and 7) (130-07, PeproTech); 50 ng/mL Angiopoietin-1 (ANG1) (Days 9 and 11) (130-06, PeproTech); and select components of the VascuLife® VEGF LifeFactors® Kit[22] (LS-1020, Lifeline Cell Technology) (Days 5, 7, 9, 11, and 13) (5 ng/mL EGF, 15 ng/mL IGF-1, 50 ug/mL ascorbic acid, 0.75 U/mL heparin sulfate, 1 ug/mL hydrocortisone).

On Days 7-16, to simultaneously induce smooth muscle cell differentiation[23, 24] with cardiomyocyte and endothelial cell differentiation, we added 2.5 or 10 ng/mL PDGF-BB (PDGFBB) (Days 7, 9, 11, 13) (100-14B, PeproTech) and 0.5 or 2 ng/mL TGF-β1 (TGFβ) (Days 7, 9, 11, 13) (100-21C, PeproTech).

From Days 5-16, unless otherwise noted, the basal medium was changed to contain B27 with insulin (RB+I) (17504044, Thermo Fisher Scientific) and was replaced every 48 hours thereafter. Cardiomyocytes within Control and cVO groups generally began spontaneously beating sometime between Days 7 and 10. Cells were collected at various time points between Days 0 and 16 for further downstream studies.

For 3D cVO creation, two methods were used. In the first method, stencils with single 2 mm center holes were placed in 24-well plates with Softwell Easy Coat hydrogels with a custom stiffness of 16 kPa (SW24-EC-C, Matrigen) and hESC-3R cells were micropatterned as described above. In the second method, to increase throughput and eliminate using stencils, each well of a black 96-well plate (M0562-32EA, Sigma-Aldrich) was freshly coated with a central, 2 uL drop of Matrigel (1:400 dilution) and was then seeded with 125,000 hESC-3R cells. Cells were allowed to attach and grow for 2-4 days forming a single, homogenous, round colony in the center of each well. The same differentiation protocol for 2D cVOs was then used to create 3D cVOs from days 0-14, using a range of CHIR from 6.0-8.5 uM.

**Hepatic Vascularized Organoid (hVO) Differentiation**

See **Fig. 7a-c** and **Supplementary Table 2** for an overview of our hepatovascular (HV) differentiation to produce hepatic vascularized organoids (hVOs). We tested 3 differentiation conditions as follows: i) Control (day 20 baseline hepatic differentiation, with no vascularization factors added); ii) hVO-D3 (day 20 hVOs created by adding vascularization factors at day 3 of differentiation); and iii) hVO-D6 (day 20 hVOs created by adding vascularization factors at day 6 of differentiation). To begin, hPSCs suspended in single cells were transferred to Matrigel-coated micropatterned surfaces as described above and differentiated in DMEM/F-12 media with GlutaMAX™ supplement (10565042, Thermo Fisher Scientific) supplemented with 0.5 mg/500 mL media of Poly(vinyl) alcohol (PVA) (P8136, Sigma-Aldrich), 1x Insulin-Transferrin-Selenium (ITS) (41400045, Thermo Fisher Scientific), 1x chemically defined lipid concentrate (11905031, Thermo Fisher Scientific), 20 uL/500 mL media of 1-thioglycerol (M6145, Sigma-Aldrich), and 1x non-essential amino acids (NEAA) (11140050, Thermo Fisher Scientific). The hVO differentiation was based on the baseline differentiation described previously[25, 26] to promote hepatic differentiation and contained concentrations of the small molecules and growth factors found to promote simultaneous endothelial and smooth muscle cell production from the results of our cVO differentiation method above. On Day 0, for mesendodermal induction, basal medium was supplemented with 100 ng/mL Activin-A (ActA) (338-AC, R & D Systems), 10 ng/mL BMP-4 (BMP4) (314-BP, R & D Systems), 3uM CHIR99021 (CHIR) (S2924, Selleck Chemicals), 100 ng/mL FGF-2 (FGF2) (100-18B, PeproTech) and 10 uM LY294002 (LY) (70920, Cayman Chemical Company). On Day 3, for foregut induction, 50 ng/mL FGF-10 (FGF10) (100-26, PeproTech) was used. On Day 6, for hepatoblast induction, 10 ng/mL FGF10 and 10 ng/mL BMP4 was used. Finally, to induce hepatocytes, 50 ng/mL Hepatocyte Growth Factor (HGF) (100-39H, PeproTech), 50 ng/mL Oncostatin-M (OncoM) (300-10, PeproTech), and 10 uM Dexamethasone (DEX) (1126, Tocris) was used.

Starting on Days 3 or 6 and continuing until Day 20, to simultaneously induce endothelial cell vasculogenesis and angiogenesis[18-22] along with the hepatocyte differentiation, we added combinations of 50 ng/mL VEGF-165 (VEGF) (Days 3 or 6, 9, 11, 13, 15, 17, and 19) (100-20, PeproTech); 5 ng/mL FGF-2 (FGF2) (Days 6 or 9, 11, 13, 15, 17, and 19) (100-18B, PeproTech); 10 uM SB431542 (SB) (Days 6 or 9, 11, 13, 15, 17, and 19) (S1067, Selleck Chemicals); 50 ng/mL Angiopoietin-2 (ANG2) (Days 3 or 6, and 9) (130-07, PeproTech); 50 ng/mL Angiopoietin-1 (ANG1) (Days 11 and 13) (130-06, PeproTech); and select components of the VascuLife® VEGF LifeFactors® Kit (LS-1020, Lifeline Cell Technology) (Days 3 or 6, 9, 11, 13, 15, 17, and 19) (5 ng/mL EGF, 15 ng/mL IGF-1, 50 ug/mL ascorbic acid, 0.75 U/mL heparin sulfate, 1 ug/mL hydrocortisone).

On Days 6-20, to simultaneously induce smooth muscle cell differentiation[23, 24] with hepatocyte and endothelial cell differentiation, we added 2.5 ng/mL PDGF-BB (PDGFBB) (Days

6, 9, 11, 13, 15, 17, and 19) (100-14B, PeproTech) and 0.5 ng/mL TGF-β1 (TGFβ) (Days 15, 17, and 19) (100-21C, PeproTech).

From Days 3-9, the basal medium was changed to contain B27 with insulin (17504044, Thermo Fisher Scientific) (RB+I) and was replaced every 48 hours. From Days 9-20, the basal medium was changed to HCM medium (CC-3198, Lonza) and was replaced every 48 hours. Cells were collected on Day 20 for endpoint analyses.

**Time-lapse Microscopy**

Initial time-lapse microscopy was performed with the Incucyte S3 Live-Cell Analysis Instrument (Essen Bioscience). Twenty-four (24)-well plates containing a single 2, 4, or 6 mm circular micropattern of undifferentiated hESC-TNNT2-GFP cells (H9-hTnnT2-pGZ-TD2, WiCell)[8] in the center of each well were subjected to the baseline cardiomyocyte differentiation condition and imaged at ten (10) timepoints from Days 0-9 (D0-D9). Phase contrast and GFP signals were obtained with a 4x objective in a square mosaic containing the entire surface area of each well. At all time points, plates were kept at $37^{o}$ C and 5 % CO2.

To increase throughput, screening time-lapse microscopy was performed with the Cytation 5 Cell Imaging Multi-Mode Reader (BioTek). Five (5) forty-eight (48)-well plates (204 out of 240 wells) were imaged over the differentiation time-period. The plates contained a single 2 mm circular micropattern of undifferentiated hESC-3R cells in the center of each well subjected to thirty-four (34) differentiation conditions, with nominally 4-6 replicates per condition, imaged at six (6) timepoints (Days 3, 5, 8, 10, 12, and 16 (D3-D16)).

Phase contrast, CFP, GFP, and mOrange signals in four (4) independent channels were obtained and the following excitation light emitting diodes (LEDs), excitation (Ex) filters, emission (Em) filters, and dichroic (Di) mirrors were used for the fluorescence channels: CFP: LED-465 nm, Ex-445/45 nm, Em-510/42 nm, Di-482 nm; GFP: LED-465 nm, Ex-469/35 nm, Em-525/39 nm, Di-497 nm; mOrange- LED: 523 nm, Ex-531/40 nm, Em-593/40 nm, Di-568 nm. Light intensity and exposure time were adjusted for each channel to minimize under- or overexposure. A 4x objective (NA 0.13) was used to acquire a 4 x 4 array in a single focal plane for all channels, resulting in a rectangular mosaic containing the circular micropattern of cells and an area surrounding each micropattern which allowed imaging of migrating cells off each micropattern over time. At all time points, plates were kept at 37$^o$ C and 5 % CO2 while imaging.

When using 48-well plates to screen 34 conditions and 6 replicates per condition, the number of wells needed was 204. Therefore, the theoretical number of acquired raw images was 78,336 images (16 images/channel/mosaic x 4 channels/mosaic x 6 days/well x 204 wells/day). The actual number of acquired raw images was 75,648, corresponding to 197 wells (out of 204 wells), with 7 wells excluded because of debris, detachment of micropatterns, and incomplete micropatterns at the start of differentiation (representing 3.4 % exclusion of wells).

After all raw images were acquired, image analysis was performed with Gen5 v3.05 software (BioTek). Images in each channel and each 4 x 4 array were stitched to create a single mosaic image; 4728 mosaics were created, and each mosaic image was downsized by 25%. Next, image processing of each mosaic was performed to reduce background independently in each channel. Finally, for each channel, thresholding was performed on fluorescence intensity along with minimum and maximum object size to calculate the total Phase, CFP, GFP, and mOrange total fluorescence area in each well for each condition at each timepoint. Each mosaic image was computationally masked by a circular "plug" to eliminate fluorescence artifact created by well edges. For each condition, values for replicate samples were summed, and the mean and standard deviation were calculated.

##### Widefield Microscopy

Either an AxioObserver Z1 (Zeiss) or a Keyence BZ-X700 (Keyence) inverted microscope was used to visualize undifferentiated hPSCs and their derivatives. The Zeiss microscope was equipped with 2.5x, 4x, 10x, 20x, 40x, and 63x objectives, a Lambda DG-4 300 W Xenon light source (Sutter Instruments), an ORCA-ER CCD camera (Hamamatsu) to visualize Phase, CFP, GFP, and mOrange channels, and ZEN (Blue Edition) v2.6 software (Zeiss) for image stitching, z-stack image capturing, and incubation control. The Keyence microscope was equipped with 2x, 4x, 10x, and 20x objectives, an 80 W metal halide lamp, a monochrome CCD camera to visualize Phase, CFP, GFP, and mOrange channels, and BZ-X Analyzer software for optical sectioning, image stitching, z-stack image capturing, and incubation control. Both microscope incubation systems were maintained at $37^{o}$ C and 5 % CO2.

##### Confocal Microscopy

Either a Zeiss LSM710 Inverted Confocal Microscope (Carl Zeiss) (Stanford Neuroscience Microscopy Service Core Facility) or a Leica SP8 White Light Confocal (Leica) (Stanford Cell Sciences Imaging Core Facility) confocal microscope was used to visualize live and fixed undifferentiated hPSCs, individual hPSC-derived cells, and cVOs. The Zeiss LSM710 confocal (AxioObserver Z1 inverted microscope base) is equipped with the following: environmental chamber to maintain samples at 37° C and 5% CO2; motorized X-Y stage and Z focus; 10x Plan Apochromat (NA 0.45); 20x Plan Apochromat (NA 0.45); 40x Plan Neofluar (oil) (NA 1.30); 63x

Plan Apochromat (oil) (NA 1.40); 405 nm diode laser (30 mW); 458/488/514 nm argon laser (35 mW); 561 nm diode-pumped solid-state (DPSS) laser (20 mW); 633 nm helium-neon (HeNe) laser (5 mW); temperature-stabilized VIS-acousto-optical tunable filter (AOTF) for simultaneous intensity control; and Zeiss ZEN Black v2.6 software. The Leica SP8 confocal (DMI 6000 inverted microscope platform) is equipped with the following: environmental chamber to maintain samples at 37° C and 5% CO2; motorized X-Y stage and galvanometer Z focus; 10x Plan Apochromat (NA 0.40); 20x Plan Apochromat (oil) (NA 0.75); 40x Plan Apochromat (oil) (NA 1.30); 63x Plan Apochromat (oil) (NA 1.40); 405 nm laser (50 mW); super continuum, white light (WLL) pulsed laser (avg. power 1.5 mW), 470 nm to 670 nm, 78 MHz; acoustical optical beam splitter (AOBS) for selection of up to 8 discrete laser lines (1 nm precision); 3 hybrid-GaAsP detectors; 2 standard fluorescent photomultiplier tubes (PMT); and Leica LAS AF software. Offline image analysis was also performed with Imaris v9.6.1 and ImageJ (Fiji) v2.1.0 software.

**Immunocytochemistry (ICC)**

Primary and secondary antibodies (**Supplementary Table 4**) were reconstituted in sterile PBS per manufacturer guidelines. Samples were fixed in IC Fixation Buffer (FB001, Thermo Fisher Scientific, diluted in PBS) or 4% paraformaldehyde (15713S, Electron Microscopy Sciences) for 10-15 minutes and washed with PBS for 10 minutes three times. To begin staining, samples were permeabilized with 0.1% Triton-X for 30 minutes at room temperature and then incubated in blocking buffer (10% normal goat serum and 0.05% Tween in PBS) for 1 hour at room temperature. Then the primary antibody (prepared in blocking buffer) was applied for 4° C overnight. Washing buffer (0.05% Tween in PBS) was applied to the samples for 5 minutes at room temperature three times. Secondary antibody (typically at 1:500 dilution) was applied for 1 hour at room temperature and followed by three 5 minutes washes in wash buffer. After a final 5-minute rinse with sterile PBS, samples were counterstained with either 4′,6-diamidino-2-phenylindole (DAPI) (D8417, Sigma) or DRAQ5™ Fluorescent Probe Solution (62251, Thermo Fisher Scientific). Samples were either analyzed immediately or stored at 4° C in PBS.

**Bulk RNA-Sequencing (bRNA-seq)**

hESC-3R (1 group, day 0, n = 3, 3 total samples; note, these samples were used for both cardiac and hepatic analyses), Control, and cVO samples (2 groups, days 2, 5, 8, 10, 12, and 16, n = 3 per sample, 36 total samples) were prepared with the Direct-zol RNA Microprep w/ Zymo-Spin IC Columns (R2060, Zymo Research) and sequenced. For hVO analysis, Control, hVO-D3, and hVO-D6 samples (3 groups, day 20, n = 3 per sample, 9 total samples) were prepared in the same fashion. Each replicate had an average of 24.4 million 150 bp long paired-end reads. FastQC v0.11.2 was used for sequencing quality assessment. Reads were then aligned to the GRCh38(hg38) reference genome using STAR v2.5.3a with splice junctions being defined in GTF file (obtained from GRCh38). An average of 85.3% of reads was aligned to the reference transcriptome.

Expression at the gene level was determined by read counts using RSEM 1.2.30. Differently expressed genes (DEGs) with fold-change were further detected by DEseq2 version 1.10.1 for comparable conditions. Based on the DEG data set, the Euclidean distances between samples, as calculated from the rlog transformation, were calculated and plotted as a heatmap.

To determine heatmap scale, the following was performed. After hierarchical clustering using Median Clustering or the Weighted Pair Group Method with Median Clustering (WPGMC) method together with the Spearman correlation (square of Euclidean distance) method for distance measurement on the log-transformed gene expression table, the values were further scaled in the row direction. This centers and standardizes each row separately to row Z-score.

Principal component analysis (PCA) was performed to extract the main information from the DESeq2 transformed (rlog) data set so that each successive axis was ordered by decreasing order of variance. Approximately 60% of all transcripts (15994 of 26470 total genes) after minimal pre-filtering to keep only rows which contained at least 10 total read/raw counts were used for PCA. PCA plots were used to visualize the batch effects and overall effect of experimental covariates. Numbers on each plot axis indicate frequency of transcripts described by each principal component.

Weighted Gene Co-Expression Network Analysis (WGCNA) was used for hierarchical clustering with dynamic tree cut to identify gene co-expression modules; the modules were further analyzed using Ingenuity Pathway Analysis (IPA) (Qiagen), described below. RNA-seq data of hESC, Control, cVO, and hVO samples are deposited in the NIH Gene Expression Omnibus (GEO) under GSE185194.

**Ingenuity Pathway Analysis (IPA)**

We used Ingenuity Pathway Analysis (IPA) (v65367011, Qiagen) of the temporal bulk-RNA-sequencing (RNA-seq) weighted gene co-expression network analysis (WGCNA) of Control and cVO samples to show vascularization and cardiogenesis regulator effect networks. In addition, we used IPA to show predicted top canonical pathways, upstream regulators, functions, regulator effect networks, and networks of Day 16 cVOs compared to D16 Control samples. WGCNA Module 16 (Pale Turquoise) within the "Vascularization Genes" cluster (**Fig. 4l**) contained 1258 analysis-ready genes with a Log2FoldChange ranging from -0.9 to 15.1 (13 down regulated and 1245 upregulated genes) and revealed regulator effect networks related to vascularization. WGCNA Module 12 (Dark Grey) within the "Cardiomyocyte/Endothelial/Smooth Muscle/Fibroblast Genes" cluster (**Fig. 4l**) contained 3719 analysis-ready genes with a Log2FoldChange ranging from -1.2 to 17.8 (46 down regulated and 3673 upregulated genes) and revealed regulator effect networks related to cardiogenesis. For analysis of Day 16 cVOs compared to D16 Control samples, 15,671 differentially expressed genes (Log2FoldChange ranging from -7.2 to 8.3) were reduced to 1001 analysis-ready genes by analyzing only differentially expressed genes with Log2FoldChange cutoff values of below -2 and above +2 and padj value equal to or greater than 0.05.

**Single-Cell RNA-Sequencing (scRNA-seq)**

Control and cVO samples (day 16, n = 6 pooled organoids per sample) were dissociated using the Pierce™ Primary Cardiomyocyte Isolation Kit (88281, Thermo Fisher Scientific) according to the manufacturer's instructions. A 10 uL sample of dissociated single cells was visualized with a fluorescent microscope (Zeiss) to verify the presence of GFP+, mOrange+, and CFP+ cells in the sample. Next, single cells (with an end-target of ~4000 cells per sample) were used to create Gel Bead-In EMulsions (GEMs) by using the Chromium Single Cell 3' Library & Gel Bead Kit v2 kit (PN-120267, 10X Genomics). Libraries were generated according to the manufacturer's instructions and sequenced using the HiSeq2500 instrument (Illumina) with a read length of PE150 base pairs at each end and ~50 M reads per sample.

Sequences were demultiplexed into FASTQ files using Cell Ranger 3.0.0 (l0x Genomics) and then aligned to the GRCh38(hg38) reference genome. Count matrices were generated using the count function with default settings. Results were initially visualized with Loupe Cell Browser v3.0.1 (10X Genomics) and then detailed downstream analysis was performed with the Seurat package[27] (v.3.1.5 and v.4.0.0) in RStudio (v1.4.1106, RStudio) using R (v3.6.1 and 4.0.4, R Core).

Barcodes, features (genes), and matrix files were loaded into RStudio as a Seurat object using CreateSeuratObject with a minimum of 5 cells and 200 genes, and percent mitochondrial genes were identified. Cells with less than 200 or greater than 5000 unique genes, or mitochondrial content over 10% were discarded. Using NormalizeData, the data was then log-normalized and scaled with a factor of 10,000. FindVariableFeatures was then used to identify the top 2,000 highly variable genes using vst as the selection method. ScaleData was then used to apply a linear transformation. Next RunPCA was used to perform linear dimensional reduction by principal component analysis (PCA). JackStraw with 100 iterations was then used to identify statistically significant ($P < 0.001$) principal components and ElbowPlot identified an 'elbow' at approximately 15 principal components (PCs). Clustering was performed using FindNeighbors and FindClusters using 15 PC dimensions and 0.2 resolution. Non-linear dimensional reduction was performed using RunUMAP and Uniform Manifold Approximation and Projection (UMAP) plots were visualized using DimPlot. Finally, differentially expressed genes were identified with FindAll Markers and violin, feature plots, and heat maps were visualized with VlnPlot, FeaturePlot, and DoHeatmap. scRNA-seq data of Control and cVO samples are deposited in the NIH Gene Expression Omnibus (GEO) under GSE185194.

**scRNA-seq Public Data**

We compared our Control and cVO scRNA-seq data to a publicly available scRNA-seg dataset from a 6.5 post-conception week (PCW) human heart[28]. The accession number for that raw sequencing dataset is European Genome-phenome Archive (EGA): EGAS00001003996. Count matrices were downloaded from https://www.spatialresearch.org/resources-published-datasets/doi-10-1016-j-cell-2019-11-025/ and analyzed initially with Seurat v3.1.5 and reanalyzed with v4.0.0.

**Cell-Cell Communication Analysis**

As we have previously described[15], to quantify the putative cell-cell communication in the cardiovascular cellulome, we obtained human ligand-receptor pairs compiled by Ramilowski et al[29]. Briefly, we defined a ligand or receptor as "expressed" in a particular cell type if its expression is higher than 0 in more than 10% of cells in that cell type for the gene encoding the ligand or receptor. We linked any two cell types where the ligand was expressed in the former cell type and the receptor in the latter to define networks of cell-cell communication. The directionality in this relationship is visible where lines connecting cell populations are colored according to the population broadcasting the ligand and connect via an arrowhead to the population expressing the receptor, thus demonstrating a capability to receive the signal. In addition, line thickness correlates with the interaction number. To plot networks, we used the igraph and circlize R packages.

**Nitric Oxide (NO) Testing**

To determine the nitric oxide (NO) producing ability of hPSC-ECs in cVOs, we used the Nitric Oxide (total) detection kit (ADI-917-020, Enzo). The enzymatic conversion of nitrate to nitrite by nitrate reductase, followed by the Griess reaction to form a colored azo dye product, allowed assessment of NO produced by the hPSC-ECs. Quantification of NO produced was achieved by measuring absorption at 540-570 nm with the use of the Cytation 5 Cell Imaging Multi-Mode Reader (BioTek), averaged from four independent experiments.

**Electrical Stimulation**

For electrical stimulation studies, multi-well plates containing Control or cVO samples were placed in a customized upright optical imaging system (121517T1UP, SciMedia) and paced in Tyrode's solution (T2397, Sigma). Samples were then point stimulated with a stimulus generator (SIU-102, Warner Instruments). The electrical stimulation consisted of a biphasic waveform with peak-to-peak amplitude of 5-10 V, pulse width of 2 to 5 ms, and frequency of 1 or 2 Hz; point stimulation was delivered through a custom-made 1 mm two-wire platinum anode/cathode electrode.

**Calcium Dye Imaging (CDI)**

For CDI, the Fluo-4 Direct Calcium Assay kit was used (F10471, Thermo Fisher) as per the manufacturer's instructions as previously described[30]. Briefly, Fluo-4 loading solution was incubated with the samples at 37° C for 30 min, washed twice with HBSS (14025092, Thermo Fisher), and resuspended Tyrode's solution. Fluorescence was measured at 495+/- 20 nm excitation and 515 +/- 20 nm emission. Videos were taken with an Evolve 512 Delta EMCCD camera (Photometrics) using Micromanager v1.4 software (Vale Lab, UCSF) at 90 fps for 10 seconds of Control or cVO samples beating spontaneously or electrically stimulated at 1 or 2 Hz. In each video frame, regions of interest (ROIs) were analyzed for changes in dye intensity f/f0, with the resting fluorescence value f0 determined at the first frame of each video. Background intensity was subtracted from all values, and plots were normalized to zero. BV-Ana software (SciMedia) was used to quantify conduction velocity and beating frequency.

**Contractility Imaging**

We assessed the contractility of Control and cVO groups in a similar manner as we have previously described[31]. Contraction of beating CMs within the micropatterns was recorded with high-resolution motion capture tracking (75 fps) using the SI8000 Live Cell Motion Imaging System (Sony Corporation). During data collection, cells were maintained under controlled humidified conditions at 37° C with 5% CO2 and 95% air in a stage-top microscope incubator (Tokai Hit). Functional parameters were assessed from the averaged contraction-relaxation waveforms from 10 sec recordings, using the SI8000C Analyzer software (Sony Corporation). The software was used to detect motion vectors and quantify beating rate, contraction velocity, relaxation velocity, and contraction-relaxation peak interval of cVOs.

**Microsphere Vascular Lumen Identification**

For vascular lumen identification, dark red (Ex/Em 660/680 nm) 0.2 um FluoSpheres™ carboxylate-modified microspheres (2% v/v) (F8807, Thermo Fisher Scientific) were diluted 1:1000 in culture medium (RPMI+B27), then added to live cardiac vascularized organoids (cVOs) and incubated at 37° C and 5% CO2 on a rocker at 15 rpm for 1 hour. cVOs were then rinsed with culture medium 3 times before confocal imaging with appropriate laser excitation and fluorescence detection to detect microspheres within cVO vascular lumen.

**Signaling Pathway Inhibition**

For NOTCH signaling pathway inhibition we used 0, 1, and 10 uM DAPT (GSI-IX) (S2215, Selleck Chemicals) added at day 0 of differentiation. For BMP signaling pathway inhibition, we used 0, 0.1, and 1 uM Dorsomorphin (P5499, Sigma-Aldrich) added at day 0 of differentiation.

**Teratogen Drug Testing**

For teratogen drug testing, we used 0 and 10 nM of fentanyl added at day 0 of differentiation.

**Machine Learning**

The general linear model and multiple regression fits were performed using scikit-learn. The general linear models had 100 coefficients that accounted for 9 growth factor/small molecule combinations and 1 bias term at 10 lags and was fit to the mean fluorescence area across 6 replicates in each condition. The multiple regression models had 7 coefficients accounting for the 7 measured days and was fit to all measured data points.

**Statistical Analysis**

Statistical analyses were performed using Prism 9 (GraphPad Software, LLC) and JMP Pro 15 (SAS Institute, Inc.) software. Data was first analyzed for normality and lognormality using the Shapiro-Wilk and D'Agostino & Pearson tests in Prism 9. If data comprised two normally distributed groups, a parametric unpaired two-tailed Student's t test was performed to determine significant differences. If data comprised two non-normally distributed groups, a nonparametric two-tailed Wilcoxon/Mann-Whitney U test was performed. If data comprised greater than two normally distributed groups, a parametric one-way or two-way ANOVA was performed. If data comprised greater than two non-normally distributed groups, a nonparametric Kruskal-Wallis test was performed. Subsequent multiple comparisons correction analysis was performed using the parametric Tukey's test for normally distributed data and the nonparametric Dunn's test for non-normally distributed data. Data are expressed as mean ± standard deviation (SD). The p-values for significance differences are as follows: *$p < 0.05$, **$p < 0.01$, ***$p < 0.001$, ****$p < 0.0001$, not significant (ns). Statistical methods were not used to predetermine sample size. Samples were randomized with respect to applied differentiation conditions and assays. The investigators were not blinded to samples, experiments, and assessments of outcome. Detailed information regarding sample size, statistical analyses, and statistical significance is included in each figure.

#### Data Availability

Bulk RNA-sequencing and single-cell RNA-sequencing data sets generated for this study are deposited in the Gene Expression Omnibus (GEO) under accession number [GSE185194].
